# Analysis of movement recursions to detect reproductive events and estimate their fate in central place foragers

**DOI:** 10.1101/562025

**Authors:** Simona Picardi, Brian J. Smith, Matthew E. Boone, Peter C. Frederick, Jacopo G. Cecere, Diego Rubolini, Lorenzo Serra, Simone Pirrello, Rena R. Borkhataria, Mathieu Basille

## Abstract

Recursive movement patterns have been used to detect behavioral structure within individual movement trajectories in the context of foraging ecology, home-ranging behavior, and predator avoidance. Some animals exhibit movement recursions to locations that are tied to reproductive functions, including nests and dens; while existing literature recognizes that, no method is currently available to explicitly target different types of revisited locations. Moreover, the temporal persistence of recursive movements to a breeding location can carry information regarding the fate of breeding attempts, but it has never been used as a metric to quantify recursive movement patterns. Here, we introduce a method to locate breeding attempts and estimate their fate from GPS-tracking data of central place foragers. We tested the performance of our method in three bird species differing in breeding ecology (wood stork (*Mycteria americana)*, lesser kestrel (*Falco naumanni*), Mediterranean gull (*Ichthyaetus melanocephalus*)) and implemented it in the R package ‘nestR’. We identified breeding sites based on the analysis of recursive movements within individual tracks. Using trajectories with known breeding attempts, we estimated a set of species-specific criteria for the identification of nest sites, which we further validated using non-reproductive individuals as controls. We then estimated individual nest survival as a binary measure of reproductive fate (success, corresponding to fledging of at least one chick, or failure) from nest-site revisitation histories during breeding attempts, using a Bayesian hierarchical modeling approach that accounted for temporally variable revisitation patterns, probability of visit detection, and missing data. Across the three species, positive predictive value of the nest-site detection algorithm varied between 87-100% and sensitivity between 88-92%, and we correctly estimated the fate of 86-100% breeding attempts. By providing a method to formally distinguish among revisited locations that serve different ecological functions and introducing a probabilistic framework to quantify temporal persistence of movement recursions, we demonstrated how the analysis of recursive movement patterns can be applied to estimate reproduction in central place foragers. Beyond avian species, the principles of our method can be applied to other central place foraging breeders such as denning mammals. Our method estimates a component of individual fitness from movement data and will help bridge the gap between movement behavior, environmental factors, and their fitness consequences.

## Background

A major goal of movement ecology is to uncover behaviors underlying movement trajectories (Gurarie et al. 2016). Knowing what animals are doing when moving a certain way can improve our understanding of the links between movement and resource dynamics, species interactions, distribution, and individual fitness (Mueller and Fagan 2008, Schick et al. 2008, Morales et al. 2010). In the past decade, the availability of animal tracking data has increased exponentially and, in parallel, methodological approaches to infer behavior from movement have proliferated (Wilmers et al. 2015).

One way to detect behavioral structure in movement trajectories is to analyze movement recursions (Berger-Tal and Bar-David 2015). Recursive movement patterns are common across taxa, as animals exhibit periodic returns to places of ecological significance such as nests, dens, foraging patches, etc. (Riotte-Lambert et al. 2013, Berger-Tal and Bar-David 2015, Bracis et al. 2018). By analyzing recursive movement patterns, researchers can identify the location of ecologically relevant places and unlock the signature of underlying ecological processes, such as responses to temporally variable resources (Bar-David et al. 2009, Berger-Tal and Bar-David 2015). In this paper, we demonstrate the use of movement recursions to detect reproductive events and estimate their fate from the trajectories of central place foragers.

Berger-Tal and Bar-David (2015) reviewed the literature on recursive movement patterns and identified three main lines of research: traplining behavior, path recursions, and the ecology of fear. While these lines of research focus on different spatial scales and have traditionally focused on different taxa, the common thread among them is the analysis of the relationship between movement recursions and temporal variation in resource availability, alone or in trade-off with predator avoidance (Berger-Tal and Bar-David 2015). For example, Garrison and Gass (1999) showed how the long-billed hermit (*Phaethornis longirostris*), a traplining hummingbird, adjusts the rate of revisitation to feeders according to changes in food delivery rates; Bar-David et al. (2009) analyzed path recursions to identify individual responses to cycles of vegetation depletion and replenishment in African buffalos (*Syncerus caffer*), and English et al. (2014) in forest elephants (*Elephas maximus borneensis*); Riotte-Lambert et al. (2013) linked recursive movement patterns of an impala (*Aepyceros melampus*) to open areas with temporally varying predation risk. As Berger-Tal and Bar-David’s (2015) review illustrates, most of the existing research on recursive movement patterns is deeply rooted in foraging ecology, with ties to optimal foraging theory, home-ranging behavior, and predator-prey interactions.

From a technical standpoint, methods to identify places of ecological relevance along individual trajectories are often based on the use of spatial buffers around visited locations (Barraquand and Benhamou 2008, Bracis et al. 2018). Other approaches that do not entail the use of spatial buffers exist, such as detecting recursions as closed paths within a trajectory (Bar-David et al. 2009) or using utilization-distribution based methods (Benhamou and Riotte-Lambert 2012). The advantage of buffer-based approaches is that they allow simple computation of metrics describing the temporal patterns of site revisitation (Bracis et al. 2018).

Temporal patterns of revisitation are used to uncover behavioral signatures underlying movement recursions (Barraquand and Benhamou 2008, Riotte-Lambert et al. 2013, Bracis et al. 2018). Different temporal metrics have proven useful to quantify recursive movement patterns for different objectives. Residence time, i.e., the cumulative time an animal spends at a certain location or area, has been used to quantify the profitability of resource patches, with high-quality patches attended for longer stretches of time (Barraquand and Benhamou 2008). Similarly, the frequency of visits has also been used to quantify the intensity of use of different areas within animals’ home ranges, with individuals visiting profitable patches more often (Benhamou and Riotte-Lambert 2012). Conversely, shorter times between visits (or time-to-return) characterize preferred areas within an animal’s home range (Van Moorter et al. 2016), as well as lower speeds (Nelson et al. 2015). The periodicity of movement recursions, i.e., the temporal repeatability of visits to a certain location, detected using Fourier- and wavelet-based methodologies, has been used to quantify responses of animals to cyclical variation in resource availability and/or predation risk (Riotte-Lambert et al. 2013). More complex analyses of movement recursions involve quantifying repetitiveness in the sequence of returns to multiple locations (Riotte-Lambert et al. 2017). Recent software developments, such as the ‘recurse’ R package (Bracis et al. 2018), have made it easier for researchers to quantify spatio-temporal patterns of movement recursions along individual tracks by identifying recurrently visited locations and calculating metrics such as number of visits, visit duration, intervals between visits, and residence time.

Important gaps remain in our current approach to the study of movement recursions. First, from a conceptual standpoint, most of the existing work has used movement recursions to gain insight on foraging ecology, home-ranging behavior, and responses to resource dynamics (Berger-Tal and Bar-David 2015). However, recursive movement patterns may lock away the signatures of other behavioral processes as well, including reproduction; yet, this application remains unexplored. For example, in central place foragers (Orians and Pearson 1979), the central return location is often associated with reproductive activities, like in the case of nests for birds (Andersson 1981, Kacelnik 1984, Burke and Montevecchi 2009) or dens for some mammals (Frame et al. 2004, Castillo et al. 2011, Olson et al. 2011). The potential for recursive movement patterns to provide information on reproduction has been so far neglected. Current literature acknowledges that, besides foraging patches, recurrently visited locations also include nests and dens (Bracis et al. 2018), but it makes no formal effort to provide ways to explicitly distinguish between different types of revisited locations. Without this ability, current approaches cannot take advantage of recursive movement patterns to specifically detect reproductive events, which is nonetheless a relevant application in the case of central place foragers. Other studies have analyzed movement tracks with the objective of detecting reproductive events along them in ungulates (caribous (*Rangifer tarandus)*, DeMars et al. 2013, Bonar et al. 2018; moose (*Alces alces*) Nicholson et al. 2019). However, these species do not necessarily exhibit movement recursions because of reproductive behavior; rather, the signature of reproductive events consists in a slow-down of movements due to limited motion capacity of calves, and thus the appropriate diagnostic metric used to detect reproduction in these species was movement rate (DeMars et al. 2013, Bonar et al. 2018, Nicholson et al. 2019). Second, while researchers have devoted effort to calculating and interpreting temporal patterns of movement recursions in terms of metrics such as residence time and periodicity (e.g., Barraquand and Benhamou 2008, Riotte-Lambert et al. 2013), little or no attention has been given to another temporal metric that can prove informative: the persistence of movement recursions, i.e., the time until the recursive movement behavior to a certain location ceases. In the case of reproduction in central place foragers, the temporal persistence of returns to a nest or den may inform us of the fate of reproductive attempts. For example, many bird species act as central place foragers while breeding, leaving the nest to go forage and returning to incubate their eggs and bring back food for their offspring (Kacelnik 1984, Alonso et al. 1994, Burke and Montevecchi 2009). Altricial bird species keep returning to the nest (often with varying frequency throughout the development of nestlings, e.g., Wiley and Wiley 1980) until their young are independent (i.e., until fledging or for some time after that; Grüebler and Naef-Daenzer 2008, Tarwater and Brawn 2010). If their attempt fails, i.e., the eggs are lost or nestlings die before fledging, altricial bird parents abandon the nest and stop visiting its location (Calder 1973, Stake and Cimprich 2003, Zangmeister et al. 2009); in some cases, nest abandonment itself can be the cause of failure (Nelson and Hamer 1995, Sabine et al. 2006, Becker et al. 2010). Thus, the persistence of recursive movements around the nest site can inform us of the fate of breeding attempts in altricial bird species, based on whether returns to the nest site persist for a duration compatible with the full developmental cycle of nestlings or cease before it is completed. The challenge with using the persistence of recursive movement patterns to pinpoint the cessation of an underlying behavior lies in the fact that movement data are collected as discrete locations rather than continuously, and thus visits to a location of interest may be missed if they happen between fixes. Moreover, in the case of breeding birds, nest attendance often decreases as the nestlings grow older (Clark 1980, Cadiou and Monnat 1996, Weimerskirch et al. 2001), which may make the daily detectability of nest visits lower later in the attempt. To account for imperfect detection of visits, the persistence of movement recursions needs to be conceptualized in a probabilistic framework.

In this paper, we present a method to apply the analysis of movement recursions to 1) detect reproductive events along the trajectories of central place foragers and 2) estimate their fate. The first step is necessary for the second, but the method serves both purposes. Unlike other available methods for the identification of revisited sites (e.g., Bracis et al. 2018), our method allows to distinguish between different types of revisited locations and isolate those of interest, thus addressing an important shortcoming of current approaches. Moreover, our method introduces the persistence of movement recursions as a useful metric to detect the cessation of behaviors of interest and allow inference on latent processes. Specifically, we use the persistence of movement recursions to estimate the fate of reproductive events in central place foragers, accounting for imperfect detection of visits to the central location. Our method thus demonstrates the use of recursive movement patterns to infer a component of individual fitness. Given the central role of fitness in ecological and evolutionary processes, estimating components of it from movement data is a long-sought goal in movement ecology (Morales et al. 2004, Singh and Ericsson 2014). As an illustration, we apply our method to identify nest sites and estimate the fate of individual breeding attempts using real GPS-tracking data for three avian species differing in their breeding habitat and ecology: wood storks (*Mycteria americana*), lesser kestrels (*Falco naumanni*), and Mediterranean gulls (*Ichthyaetus melanocephalus*).

## Methods

### Method description

Our method is composed of two parts: first, the detection of breeding sites, and second, the estimation of reproductive fate (Fig. 1). The workflow is implemented in the R package ‘nestR’, which was designed with avian species in mind (https://github.com/picardis/nestR; Picardi et al. 2019). Although the purpose of this paper is to introduce the method *per se*, we included a synthetic description of the package structure and functions in Additional file 1; Additional file 2 provides the complete package documentation; moreover, the ‘nestR’ package vignette includes worked examples of our study cases (Additional file 3).

**Figure 1.**
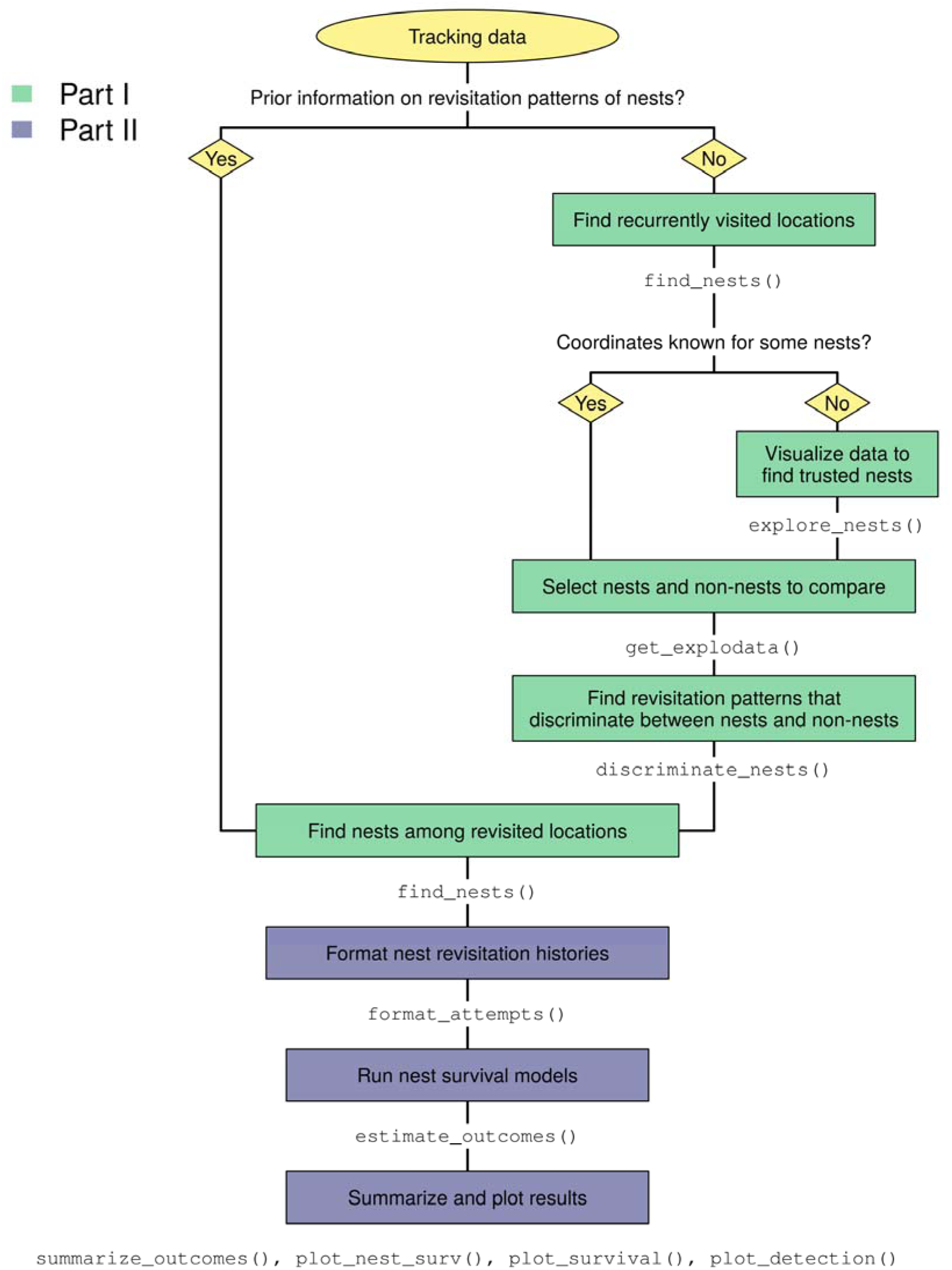
Method workflow. Workflow of the analysis to identify nest sites (Part I) and estimate reproductive fate (Part II) from telemetry data. The R package ‘nestR’ includes functions to tackle each of the steps depicted in the boxes.

#### Nest-site detection

The objective of Part I is to identify nest sites (and thus, breeding events) along individual trajectories (Fig. 1). Nest sites are identified as repeatedly visited locations along individual trajectories (Fig. 1, “Find recurrently visited locations”). Returns to a location are defined as returns to a circular area of a user-defined radius buffering each point of the trajectory (Fig. 2A). Using buffers accounts for the spatial scattering of GPS points around the actual central location due to both behavior and GPS error (Frair et al. 2010). The buffer size sets the spatial scale at which revisitation patterns will be calculated and should be chosen according to the expected scale of movements, which should be small in the case of a nest (compared, for example, to returns to a same foraging area but not exact location; Bracis et al. 2018). We recommend using this rationale to choose a reasonable buffer size for the initial round of analysis; then, we recommend performing a sensitivity analysis on the effect of buffer size on nest-detection performance and refining the spatial scale of the analysis to optimize performance, if necessary (Additional file 4).

**Figure 2.**
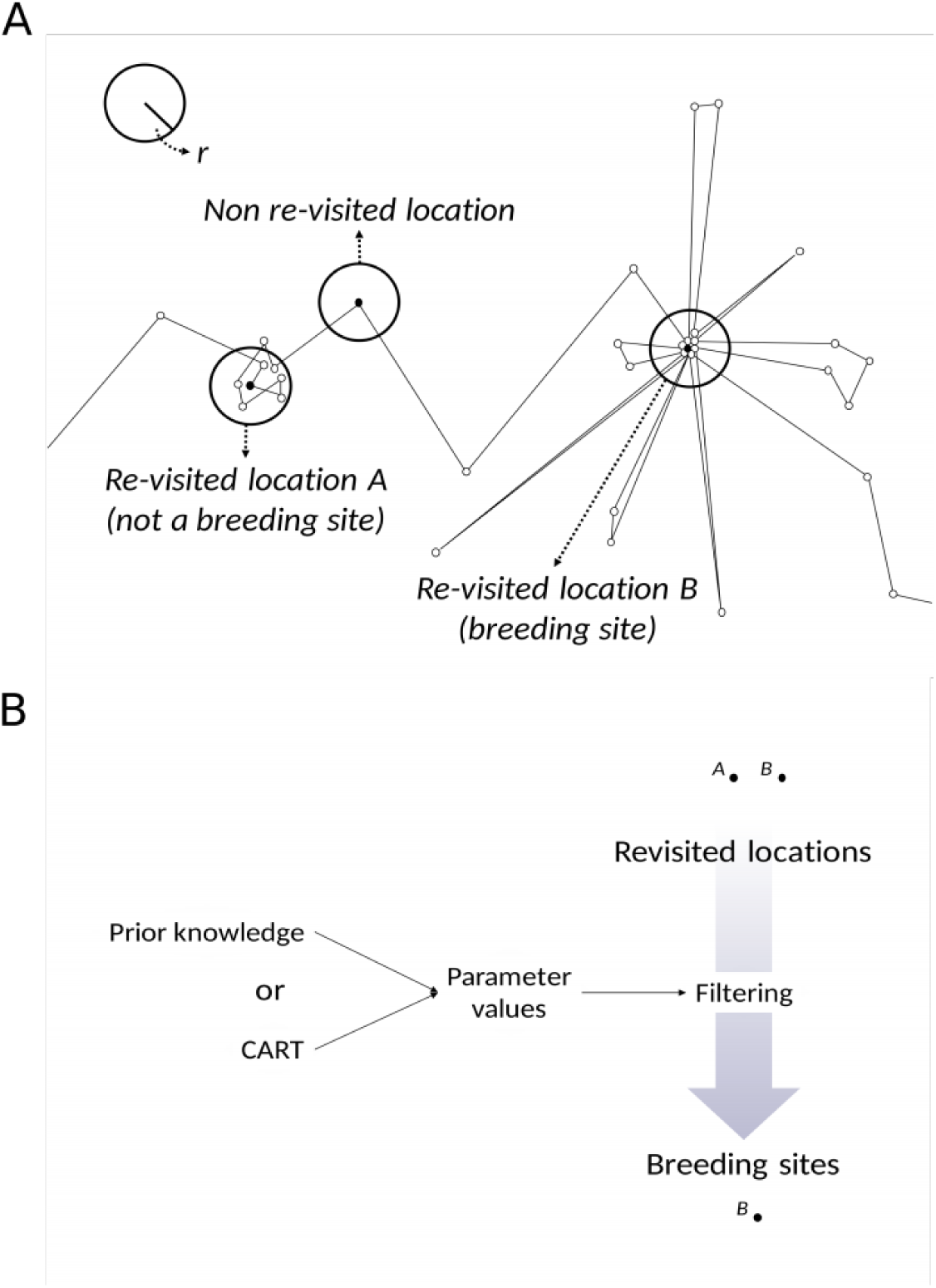
Identification of revisited locations and breeding sites among them. Schematic diagram depicting how revisited locations (and breeding sites among them) are identified along individual tracks. A) A buffer of user-defined radius *r* is placed around each location of the trajectory. Buffers that contain any other locations in addition to the one they are centered on are revisited locations. A fraction of these will be breeding sites. B) To detect breeding sites specifically, revisited locations are filtered based on their temporal patterns of revisitation using parameter values that are diagnostic of breeding sites. Parameter values can be chosen based on prior knowledge or based on the comparison of known breeding and non-breeding sites with CART.

For the purpose of distinguishing between different types of revisited locations, we describe revisitation patterns using the following set of temporal parameters: the maximum number of consecutive days a location is visited; the percentage of days it is visited between the first and last visit; and the percent fixes at a location on the day with longest attendance (we later refer to this as the “top day”, in short). We calculate these parameters for every revisited location that we initially detect, each of which is a potential nest site. Parameter values are then used as diagnostic features to filter actual nest sites among revisited locations (Fig. 1, “Find nests among revisited locations”), based on the rationale that revisitation patterns differ between nest and non-nest sites (i.e., other types of revisited locations such as foraging patches, roosts, etc.). Nest sites are often visited for longer stretches of consecutive days, on more days, and more frequently or for longer within a day than other types of revisited locations; the parameters we use to describe revisitation patterns are meant to capture these diagnostic behaviors. The method can be tailored to different case studies by restricting the analysis within the breeding season for a given species and accounting for data sampling rate and fix failure rate (Additional files 2 and 3).

Several approaches are possible to identify sets of parameter values to distinguish nest from non-nest sites. The simplest option is to use prior knowledge of the behavior of the focal species or expert opinion (Fig. 1, “Prior information on revisitation patterns of nests?”). If such information is available, users can directly skip to the final step of the first part of the workflow (Fig. 1, “Find nests among revisited locations”). Alternatively, we propose a data-driven approach to agnostically identify optimal parameter values based on Classification And Regression Trees (CART; De’ath and Fabricius 2000). CART can be used to find the optimal set of parameter values that discriminate between nest sites and other types of revisited locations, based on the comparison of known nest and non-nest sites (Fig. 1, “Find revisitation patterns that discriminate between nests and non-nests”). If the true location of nests is known for a subset of the data (Fig. 1, “Coordinates known for some nests?”), researchers can use these known nests and compare their revisitation patterns with those of other locations that are not nests. If on-ground data are not directly available, an alternative is to visually explore the data and identify trusted nest sites, where possible (Fig. 1, “Visualize data to find trusted nests”). For example, likely nest sites may be recognized in some species based on habitat features or proximity to known breeding colonies. Once known or trusted nest sites are identified, non-nest sites can be selected based on a criterion of temporal overlap (Fig. 1, “Select nests and non-nests to compare”); locations revisited simultaneously with a breeding attempt can be considered non-nest sites, assuming birds cannot breed in two places at the same time (which may not be true in all study systems). Once a dataset of known nest and non-nest sites is built, users can apply CART to this dataset and prune the tree to the optimal number of nodes based on a minimum relative error criterion (De’ath and Fabricius 2000). More sophisticated classification tools, such as random forests (Breiman 2001), may also be appropriate for this task, but CART has the advantage of providing outputs that are easy to interpret biologically, as it returns numeric cut-offs for each of the input parameters that most efficiently split the data between the groups of interest.

The (one or more) sets of parameters identified by CART as best discriminating between nest and non-nest sites can be manually applied to the complete set of revisited locations to identify nest sites among them (Fig. 1, “Find nests among revisited locations”; Fig 2B). In case multiple nests that satisfy the criteria are identified for one individual, if any of them temporally overlap with each other (and again assuming birds cannot breed in two places at the same time), the recommended option is to pick the most visited candidate and discard those that temporally overlap with it.

Because subsequent analyses rely on accurate nest-detection results, it is of paramount importance that users verify that performance of the nest-detection algorithm is satisfactory before proceeding with the estimation of reproductive fate. Both failing to identify some nests or identifying spurious ones might result in mis-estimating reproductive fate down the line. On one hand, failure to identify a nest in an individual’s track would lead to concluding that that individual did not attempt to breed; the consequences of this wrong conclusion might weigh more or less according to, first, whether the missed attempt was successful or not, and second, whether we are interested in assessing net reproductive success or relative to effort. For instance, if we are not interested in measuring reproductive fate relative to effort, concluding that an individual did not attempt to breed or attempted and failed does not make a difference in terms of net output, but it does when considered in light of return per energy investment. Both behavior and data quality might affect the probability to miss a nest: for example, the risk of failing to detect a nest is higher if data resolution is low or if nest visits are infrequent. In most cases, we would expect successful attempts to not be missed except in case of persistent malfunctioning of a GPS tag; thus, missed detections are likely to be biased towards failed attempts – especially early failures, where the nest was not visited for long enough to be picked up by the algorithm. On the other hand, identifying as a nest a revisited location which is actually not a nest might lead to concluding that an individual bred when it actually did not or failed at breeding when it actually did not attempt. This type of error is affected by behavior more than data characteristics, and it may be more likely for some species than others: for example, the signature of recursive movements around a nest may be more likely to be confounded with non-breeding movements in perch-and-wait predators, who spend long and frequent stretches of time on the same perch, than in seabirds, who spend most of their time wandering at sea when not breeding. The gravity of detection errors depends on specific study objectives (e.g., measuring net fate vs. relative to effort) and committing one type of error might be more undesirable than committing the other; we recommend that researchers think critically about how much they care about minimizing each and interpret performance metrics accordingly.

We propose four metrics to quantify nest-detection performance: positive predictive value, i.e., the proportion of true nest sites among those that are found; sensitivity, i.e., the proportion of true nest sites that are successfully identified; false negative rate, i.e., the proportion of true nest sites that the algorithm failed to identify; and false positive rate, i.e., the proportion of non-nest sites that are identified as nests. The first three metrics can be evaluated using data on known or trusted nest-site locations; the fourth metric can be assessed using non-breeder trajectories, for example outside of the breeding season or for individuals under reproductive age.

#### Reproductive fate estimation

Part II of the workflow addresses the second objective of our method: estimating the fate of the reproductive events identified in Part I (Fig. 1). The fate of breeding attempts is estimated based on the temporal persistence of returns to the nest: essentially, we are asking whether the tracked individual kept revisiting the nest for long enough to carry the attempt to term, based on how long the developmental cycle is in the focal species. A breeding attempt is considered successful if the nest site was visited until the end of a complete breeding cycle for the focal species, which includes nest-building, egg-laying, incubation, and chick-rearing until the nestlings reach autonomy and no longer receive parental care. Thus, we define success of a breeding attempt as survival of at least one nestling until fledging and failure to none. First, for each nest site identified in Part I, we compile a history of nest revisitation (Fig. 1, “Format nest revisitation histories”), by converting the movement track to a presence/absence time series at the nest (0/1 for each GPS point). This presence/absence time series will then be treated as capture-recapture data and used to estimate the fate of each breeding attempt using a survival analysis in a Bayesian hierarchical modeling framework (Fig. 1, “Run nest survival models”). Modeling persistence of nest revisitation with a hierarchical modeling approach allows us to take into account imperfect detection of nest visits and missing GPS fixes, as described in the following paragraph.

The nest survival model specification includes two processes: the survival process, which is not directly observable, and the observation process, which is the revisitation history. Much like a Bayesian implementation of a Cormack-Jolly-Seber capture-mark-recapture model (Lebreton et al. 1992, Schaub and Royle 2014), the latent nest survival variable, *z*, is modeled as a Bernoulli variable at the daily scale as a function of survival status and daily survival probability, *φ*, at the previous time-step:

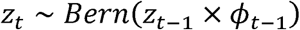

Observed visits on a given day are modeled as a binomial variable as a function of current nest survival status, probability of visit detection, *p*, and number of GPS fixes available, *N*, on day *t*:

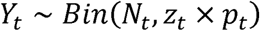

Where the probability of detection is conditional to *N* and to the nest being alive on that day:

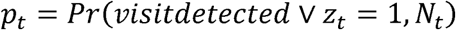

Reproductive fate is defined as the probability *P* that the nest was still surviving on the last day of the theoretical duration of a complete breeding attempt, *T*:

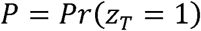

Both survival and detection probability are modeled using a binomial Generalized Linear Model (GLM) as a function of the day of the attempt:

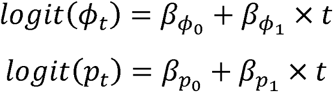

The model is fully specified by using uninformative priors on the *β* parameters, in this case a normal distribution with a mean of 0 and precision of 1e-5. In the current implementation, daily survival and detection are assumed to be the same for all nests in the population. The model outputs daily estimates of survival and detection probability at the population level, as well as daily survival estimates for each breeding attempt along with credible intervals. From these values, results can be summarized in different ways: for instance, they could be expressed in terms of value of *P* on the final day of the breeding cycle for each attempt, or the last day when *P* was greater than a chosen threshold (e.g., 90%), and so on. The MCMC (Markov Chain Monte Carlo) algorithm is implemented in JAGS (Just Another Gibbs Sampler; Plummer 2003) via the R package ‘rjags’ (Plummer 2018).

Assumptions underlying the nest survival model include: 1) birds are tracked for the entire duration *T* of a complete nesting attempt (if birds were tagged part-way through an attempt, *T* needs to be adjusted by subtracting the age of the nest (in days) at tagging); 2) the GPS tag does not permanently fail before the end of the attempt; 3) parents visit the nest until fledging, or nestling mortality is negligible between the time when parental care is interrupted and fledging; 4) parents stop visiting a nest after failing.

### Study cases

We applied our method to GPS-tracking data for 148 individual-years for wood storks (henceforth storks), 56 for lesser kestrels (henceforth kestrels) and 29 for Mediterranean gulls (henceforth gulls; Table 1). All tags were solar-powered and recorded fixes primarily during daytime. Details about devices, settings, harnesses, and study areas regarding storks and kestrels can be found in Borkhataria et al. (2008) and Cecere et al. (2018), respectively. To find nest sites, we restricted the analysis to the breeding season only for each species (Table 1). While both kestrels and gulls have a well-defined breeding season between April and August in our study areas (Snow et al. 1997), storks in the southeastern U.S. can breed at slightly different times of the year depending on latitude (Coulter et al. 1999; Table 1). In this case, we used a conservative approach and only excluded the period where no breeding activities were expected to occur anywhere in the range.

**Table 1.**
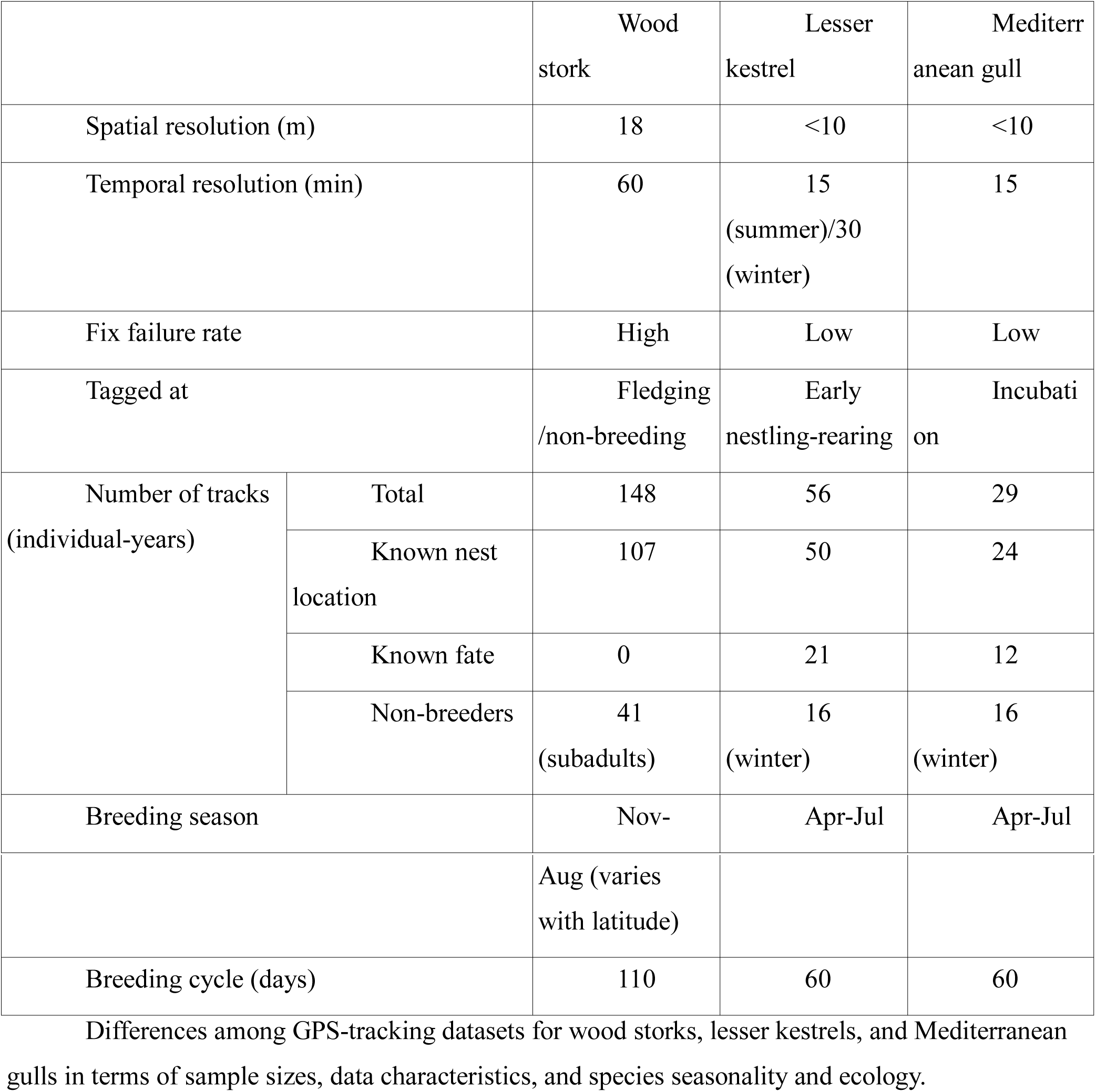
Datasets.

Given the spatial resolution of the GPS data (Table 1) and the expected scale of movements around the nest site for all three species, we initially ran the nest-detection algorithm using a buffer of 40 m around each GPS position. We then ran a sensitivity analysis on buffer size using values between 10 and 100 m (Additional file 4), picked the radius that maximized performance for each species, and re-ran the analysis to refine results.

We initially screened trajectories for any revisited locations using non-constraining values for parameters describing revisitation patterns (thus not applying any filtering). We then used on-ground data on known nest locations to select true nests and non-nest sites from the revisited locations. Kestrels and gulls were captured at the nest site (Table 1), so the location of the nest was known. For storks, on-ground data on nest locations was available for 10 individual-years (Bear D., unpublished data). We explored the remaining stork trajectories and identified those for which the top visited location was at a known breeding colony (data from U.S. Fish and Wildlife Service 2018). We marked these as trusted and treated them as known nest sites for the rest of the analysis.

We used CART to compare revisitation patterns between nest and non-nest sites, and used the resulting sets of parameter values to filter nest sites among revisited locations in the trajectories of breeding individuals. We only retained individual-years where data exceeded the minimum number of consecutive days visited indicated by CART (Table 1). Even when CART did not suggest that the number of consecutive days visited was an important predictor of true nest sites, we chose a reasonable value to use as a threshold for this parameter, as we did not expect to have enough power to discern nest from non-nest sites for attempts that failed in the first handful of days. We only retained the candidate with the most visits among any sets of breeding attempts that were temporally overlapping. We used non-breeder trajectories (sub-adults in the case of storks, non-breeding season data in the case of kestrels and gulls) to validate our results against false positives. We calculated positive predictive value of our algorithm as the percentage of nest sites that were known among those we found for each species; sensitivity as the percentage of known nest sites that were identified; false negative rate as the percentage of known nest sites that we failed to identify; and false positive rate as the percentage of non-breeding individual-years for which we erroneously identified a nest site.

We fit the nest survival model described above to estimate the fate of identified breeding attempts, using only individual-years for which the tag was active throughout the attempt to meet model assumptions (Table 1). Since kestrels and gulls were captured after they had already started breeding (immediately after hatching and in late incubation, respectively, although the exact age of the nest at tagging was unknown), the initial part of every breeding attempt was missing from the movement data. To account for this, we subtracted the theoretical number of days until hatching (for kestrels, 25 days) and late incubation (for gulls, 20 days) from the value of *T* (Table 1). We evaluated performance of the method by comparing survival estimates to known fates.

## Results

The initial screening with no filtering identified 9871 revisited locations (i.e., potential nest sites) for storks, 511 for kestrels, and 1379 for gulls. Results from CART showed that the optimal set of parameter values to discriminate nest from non-nest sites was 14 minimum consecutive days visited and 79% minimum nest attendance on the top day for storks, 7 minimum consecutive days visited for kestrels, and 26% minimum attendance on the top day for gulls (Fig. 3). Because CART did not indicate a minimum number of consecutive days visited for gulls, we added a reasonable constraint for this value by exploring the data and determining which value would allow us to rule out most non-nest sites while retaining most nest sites (8 days). Based on results of the sensitivity analysis, we used buffers of 30, 60, and 20 m for storks, kestrels, and gulls, respectively (Additional file 4). By filtering revisited locations using these optimal buffer sizes and the parameter values indicated by CART, we identified 108 nest sites for storks, 46 for kestrels, and 24 for gulls, which closely matched the number of nest sites we were expecting to find (Table 1). As a consequence, the positive predictive value of the algorithm ranged between 87-100%, the sensitivity between 88-92%, and the false negative rate between 8-12% for the three species (Table 2). The false positive rate was 2% for storks and 0% for gulls but reached 50% for kestrels (Table 2). The probability of detecting nest visits decreased throughout the breeding attempt for all three species, while survival remained constant (Fig. 4). We correctly estimated the fate of 100% of breeding attempts for gulls and 86% for kestrels (1 failure and 2 successes incorrectly estimated; Fig. 5). No data on true fates were available for storks, therefore we were unable to verify survival estimates for this species.

**Table 2.** Nest-detection performance metrics.

|  | Wood stork | Lesser kestrel | Mediterranean<br>gull |
| --- | --- | --- | --- |
| Positive<br>predictive value | 87.04% | 100.00% | 91.67% |
| Sensitivity | 87.85% | 92.00% | 91.67% |
| False negative<br>rate | 12.15% | 8.00% | 8.33% |
| False positive<br>rate | 2.45% | 50.00% | 0.00% |

Performance metrics of the nest identification algorithm (see Methods for definitions).

**Figure 3.**
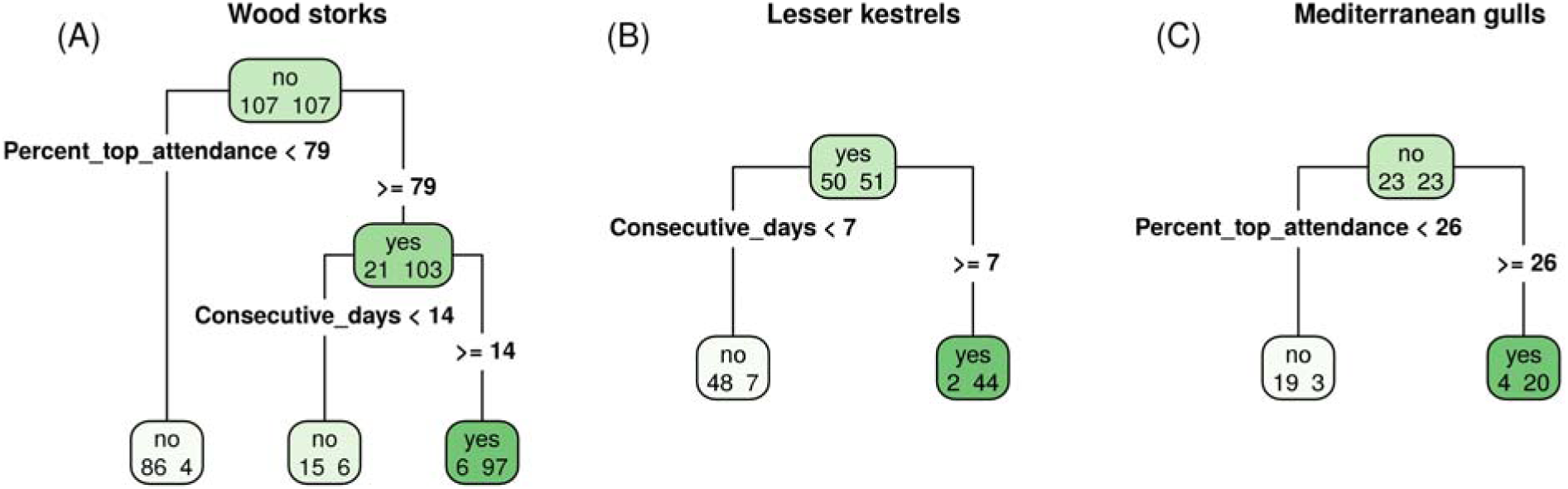
Output of Classification and Regression Trees on parameters describing nest revisitation patterns. Output of CART to discriminate nest and non-nest sites in A) wood stork, B) lesser kestrel, C) Mediterranean gull. Within each node (box), the number of known non-nest and nest sites are reported on the left and right, respectively. The root node (top) is recursively split into two until the terminal nodes (bottom). The criterion used to split each node is shown on the corresponding stems (bold font). The label on each node represents the class that was assigned to the content of that node (nest site for “yes” boxes, non-nest site for “no” boxes). Thus, the number on the right in “yes” terminal nodes and the number on the left in “no” terminal nodes correspond to correct classifications, while the number on the left in “yes” terminal nodes and on the right in “no” terminal nodes correspond to incorrect classifications.

**Figure 4.**
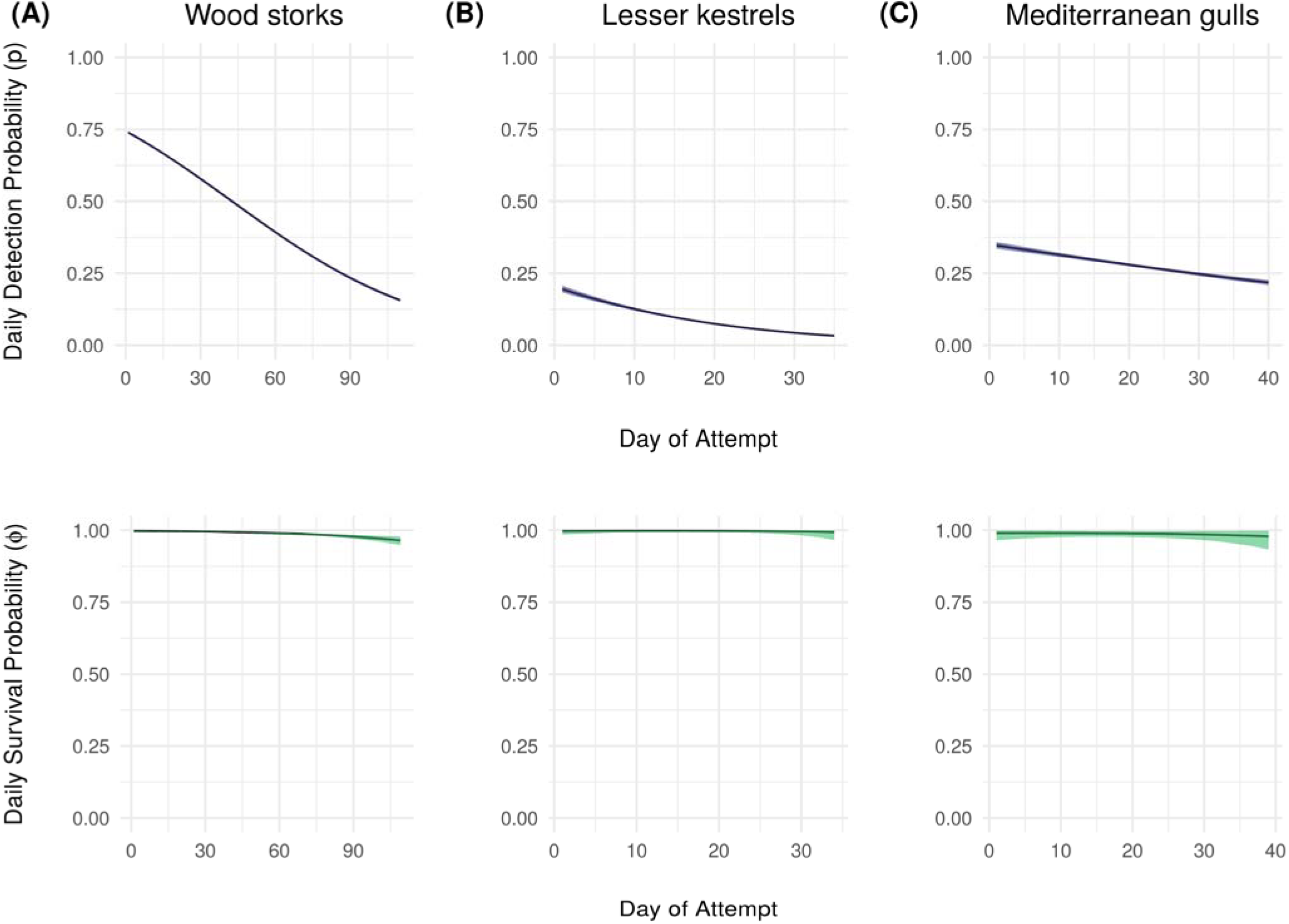
Probability of survival and visit detection. Probability of visit detection (top row) and survival (bottom row) through time (days of the attempt) estimated at the population level for A) wood stork, B) lesser kestrel, C) Mediterranean gull. Ribbons indicating 95% credible intervals are shaded.

**Figure 5.**
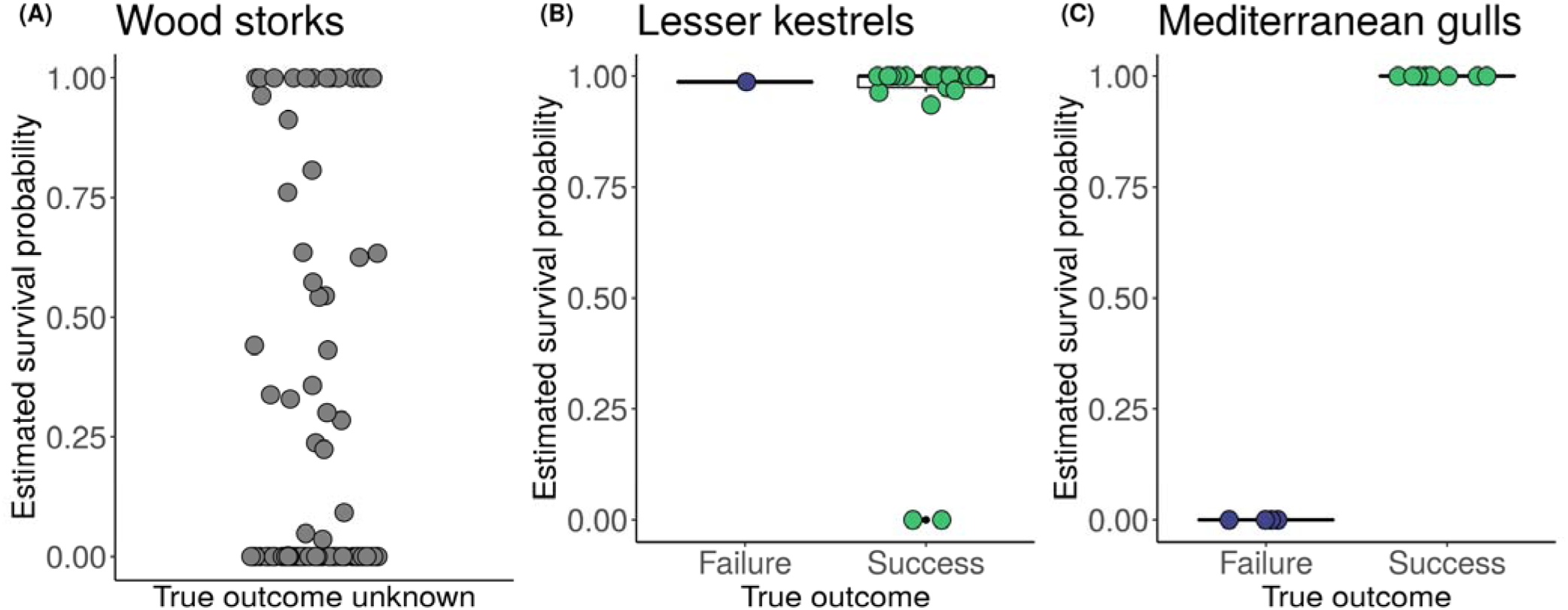
Nest-survival estimates. Estimates of probabilities of success, i.e., *P(z*_*T*_ *= 1)* for breeding attempts of A) wood stork, B) lesser kestrel, C) Mediterranean gull. For kestrels and gulls, estimates are plotted in relation to their true fate. True fate was unknown for storks. Raw data points are shown as dots (purple for failures, green for successes, gray when true fate is unknown) overlaid to boxplots (black).

## Discussion

We presented a method to detect reproductive events and estimate their fate from recursive movement patterns of central place foragers. As an illustration, we applied our method to identify nest-site locations and estimate the fate of breeding attempts from GPS-tracking data for three avian species differing in breeding behavior and ecology (Fasola and Canova 1992, Coulter et al. 1999, Gustin et al. 2017). To our knowledge, this is the first time the analysis of movement recursions is applied to infer reproduction; more broadly, ours is also among the first attempts to infer the reproductive component of fitness from telemetry data (but see DeMars et al. 2013, Bonar et al. 2018, Nicholson et al. 2019), and to our knowledge, the first applied to birds. Our method broadens the applicability of current approaches and tools for the analysis of movement recursions, opening the possibility of inferring reproduction in addition to traditionally targeted latent processes such as foraging behavior and home-ranging (Berger-Tal and Bar-David 2015). The application of recursive movement analysis to estimating the fate of reproduction is made possible by introducing persistence of movement recursions as a metric to detect the cessation of latent behaviors. Conservation and management applications may both benefit from the availability of our method and its implementation in an open-access, user-friendly R package, ‘nestR’. Knowledge of the biology and ecology of the target species and careful consideration of data characteristics and limitations are critical for successful use of the tools we presented.

### Performance of nest-site detection

Our nest-site detection method performed well for all three species, allowing us to correctly identify most or all known nest sites from movement trajectories of breeding individuals. We achieved high positive predictive value (87-100%) and sensitivity (88-92%) for all species. Importantly, the positive predictive value quantifies how many of the nest sites we found were known, which does not necessarily imply that the remaining were non-nest sites: it is possible that those we were unable to confirm for storks and gulls included second attempts (true but unknown nest sites) in addition to non-nest sites, as both species may attempt to breed again at a different location if their first clutch fails early in the season (Rodgers et al. 1987, Cecere pers. obs.). In support of this possibility, all unknown nest sites we found for gulls were from birds whose known attempt failed early on, and they were thus plausible second attempts. The same might be true for storks, although we did not have on-ground data to confirm it. False negative rates were low for all species (8-12%) and mostly associated with early failures: 2 out of 2 nest sites that we failed to identify for gulls and 2 out of 5 for kestrels corresponded to attempts that failed before the enforced limit of consecutive days visited (as early as the day after tagging in the case of gulls). This may be true for storks as well, where the breeding attempts we were unable to identify might have failed before the 14-day mark. Not identifying breeding attempts whose duration does not exceed the minimum constraint applied is a logical implication of the approach rather than a failure of the algorithm. The remaining 3 nest sites that we were unable to identify for kestrels did not fail within the first week, but were never visited for 7 consecutive days. False positives were none or negligible for gulls and storks (0% and 2% respectively), but reached 50% for kestrels. This is likely explained by species-specific behavior: kestrels spend long stretches of time and consecutive days on a perch while scanning for prey or resting (Hernández-Pliego et al. 2017). Distinguishing these patterns of attendance and revisitation from those of a nest might be challenging without applying restrictions based on seasonality and geographical area (e.g., breeding versus wintering range).

Error rates for nest-site identification vary in importance depending on the study objectives. If the objective is to estimate reproductive fate (especially if regardless of effort), ensuring that attempts are not missed should receive priority over avoiding the selection of non-nest sites. Any revisited location that gets erroneously identified as a nest site would likely be classified as a failed attempt eventually anyway. In this case, we suggest that researchers may want to focus on minimizing false negatives. Conversely, if the objective of a study is, for instance, to analyze factors associated with nest-site selection, minimizing false positives should be the priority.

Once on-ground data on nest locations are used to identify parameter values to distinguish nests among revisited locations, these parameter values can then be applied to new individuals of the same species for which on-ground information is not available, assuming other data characteristics are the same. If CART is the tool of choice to inform the choice of parameter values, we recommend that parameter values suggested by the algorithm should be critically evaluated for their biological significance before use, and that adjustments should be made as needed based on knowledge of the species biology. Future efforts to improve our method for the identification of nest locations will include incorporating uncertainty in our estimates of nest sites, allowing us to interpret classification results in a probabilistic framework.

### Performance of reproductive fate estimation

We correctly estimated reproductive fate of 100% of breeding attempts for gulls and 86% for kestrels, with probability of success estimated as *P* > 0.97 for true successes and as *P* = 0 for true failures. The remaining attempts were two successes that we estimated as failures (*P* <= 0.3) and one failure that we estimated as a success (*P* = 0.98). The two attempts that we erroneously estimated as failures corresponded to one male and one female kestrel whose original clutch included four eggs and was partially lost, leading to two and one fledglings, respectively. Partial brood loss may lead to a faster completion of the breeding cycle (Stenning 1996), which may have compromised our ability to detect these attempts as successful as they did not reach the benchmark *T* = 60. Specifically, one of the two attempts was completed within 27 days of tagging, which corresponds to *T* = 52. However, the other attempt was completed within 33 days of tagging, corresponding to *T* = 58, which is a similar duration to other successful attempts that we estimated correctly. In this case, our inability to recognize the attempt as successful might have depended on behavioral differences between parents, whereby the male we were tracking might have interrupted parental care before the female did. This result highlights the importance of taking into account sex differences in breeding behavior, where that applies (Hernández-Pliego et al. 2017). For example, in species exhibiting uniparental care, inference should only be based on the sex that carries out parental care (e.g., Birks 1997). The failed attempt that we erroneously estimated as successful corresponded to a male that occasionally visited the nest site after failing, thus violating one of the assumptions of our model. Unfortunately, this was the only failed attempt for kestrels in our dataset, which makes it difficult to generalize our ability to estimate nest failures for this species. Overall, the three instances of incorrect estimation might suggest that model assumptions, such as interruption of nest visits after failure, might not always hold across species; or that the duration of a complete breeding cycle may be too variable to lend itself to generalizations in some species; or that not knowing the exact age of the nest at tagging might have reduced our power to distinguish late failures from successful attempts that were completed in less-than-average time.

We did not have on-ground data to validate our estimation of reproductive fate for storks; however, most attempts were estimated as either *P* = 1 or *P* = 0, while intermediate values (between 0.25 and 0.75) were relatively rare (14 out of 109). This is an important result given that data for storks were at lower temporal resolution compared to kestrels and gulls (Table 1). Low temporal resolution of data in combination with decreasing frequency of nest visits can, in principle, increase the uncertainty of fate estimation by reducing probability of visit detection especially towards the end of a breeding attempt (Fig. 4). Thus, the higher proportion of intermediate values for estimates of breeding success probabilities we observed in storks compared to kestrels and gulls was to be expected, but results were still rather polarized, suggesting that the method is largely able to distinguish between successes and failures at this temporal resolution, given the frequency of nest visits in storks.

#### Innovations in the analysis of movement recursions

Our method introduces key innovations compared to current approaches to the analysis of recursive movement patterns. First, compared to other point-based methods, our method provides the ability to distinguish between locations that are revisited for different purposes and are thus functionally different from one another from an ecological point of view. Bracis et al. (2018) acknowledge that revisited locations can include foraging patches, roosts, nests, dens, etc. and they speculate post-hoc on what type of revisited locations are those they identify for the vulture in their study, but their method does not provide a way to explicitly tell them apart. Similarly, Riotte-Lambert et al. (2017) recognize that foraging patches are not the only type of revisited location, but they do not attempt to distinguish between these types *a-priori*, rather they interpret the ecological function of recursive movements *post-hoc*. Other approaches (e.g., Bar-David et al. 2009, English et al. 2014) focus on quantifying individual foraging behavior while implicitly assuming that any revisited location is a foraging patch (which may be a valid assumption in species that do not exhibit recursive movements when reproducing). Finally, approaches that aim to quantify the intensity of use of different areas within individual home ranges usually do not attempt to discriminate between areas used for different functions (e.g., Barraquand and Benhamou 2008, Benhamou and Riotte-Lambert 2012, Van Moorter et al. 2016). Second, our method draws attention to a neglected temporal metric, i.e., the persistence of movement recursions. Previous studies focused on inferring latent processes from recursive movement patterns use residence time (Barraquand and Benhamou 2008), time-to-return (Moorter et al. 2016), periodicity (Riotte-Lambert et al. 2013), or repeatability of visitation sequences (Riotte-Lambert et al. 2017) as temporal metrics, but not the persistence of movement recursions. We introduced a probabilistic approach to quantify the persistence of recursions to a central location (nest site) and to estimate the cessation of a latent behavior (administering parental care to nestlings) using a survival analysis in a Bayesian hierarchical modeling framework. This approach allowed us to account for imperfect detection of visits to the central location, thus incorporating uncertainty into our estimates of temporal persistence. Combined, the two elements of novelty that we described unlocked the possibility of using recursive movement patterns to infer reproduction from movement data in central place foragers.

Among birds, important exceptions to the applicability of our method are precocial species and nest parasites, where parental care is limited (Starck et al. 1998) or absent (Lyon and Eadie 1991). Another limitation of our approach is that it does not provide estimation of reproductive success in terms of number of offspring, but only in terms of overall success or failure. Under this aspect, our method does not compare to the level of detail obtainable with conventional field methods (e.g., Coulson and Porter 1985, Mayer and Ryan 1991, Bruant et al. 2019).

We demonstrated an application of our method to three avian species, but the method can in principle be adapted to other central place foragers too. For example, some carnivores such as wolves (*Canis lupus*, Frame et al. 2004), Canada lynx (*Lynx canadensis*; Olson et al. 2011), and Pampas foxes (*Lycalopex gymnocercus*, Castillo et al. 2011) also act as central place foragers when rearing their young, performing trips to and from the den until the offspring are weaned. Some denning mammals, including Canada lynx, use multiple dens sequentially within a single breeding season (e.g., Olson et al. 2011), so the method may need to be adjusted to sequentially identify these, and the persistence of movement recursions would then be analyzed in relation to multiple locations. The persistence of returns to the den (or dens) could be used like in the present study to estimate whether the young survived until weaning. More broadly, the principle of distinguishing types of revisited locations that serve different ecological functions can be applied across species and contexts.

### Synthesis and significance

Our method can appeal to researchers with different objectives. First, it can be used by researchers who aim to detect reproductive events from movement data in central place foragers, such as any non-parasitic altricial bird species. Researchers may want to identify reproductive events for analyses of reproductive-site fidelity or selection or to discover the geographic location of unknown breeding sites. Researchers interested in this application of our method can stop at the first part of the workflow. Second, the method can serve researchers who aim to estimate whether individual reproductive attempts were successful or not. This application of our method can be especially useful when collecting field data on reproductive fate is impractical or risky, for example because of inaccessible locations or high risk of disturbing reproductive activities (Götmark 1992, Mayer-Gross et al. 1997, Etterson et al. 2011, Wilmers et al. 2015). Researchers interested in this application of our method will use both parts of our method sequentially. Finally, having pinpointed the location of reproductive sites and estimated the fate of reproductive events, researchers can pair this information with environmental conditions experienced by breeding individuals at locations visited away from the reproductive site (Cagnacci et al. 2010, Pettorelli et al. 2014), which are usually unknown when data on reproduction is collected in the field and individuals are not tracked. Knowledge of sites visited by breeding animals when not at the reproductive site can be used, for example, to test for foraging conditions that may enhance or hamper reproductive success. Our method can be applied both in situations of opportunistic use of historical tracking data or in cases where the study is explicitly designed with these objectives in mind. Our work pioneers the use of recursive movement analysis to obtain estimates of reproductive events from tracking data in central place foragers. Reproductive fate is an important component of fitness, and estimating it from tracking data will help establish the long-sought bridge between movement and fitness at the individual level (Nathan et al. 2008, Morales et al. 2010).

## Conclusions

We presented a method to locate breeding sites and estimate the fate of breeding events from the movement trajectories of central place foragers. Our approach complements the existing literature on recursive movement patterns by allowing the distinction of functionally different types of revisited locations, and by introducing the temporal persistence of recursions as a metric to detect the cessation of latent behaviors. In our case, the focal type of revisited location are breeding sites and the latent behavior of interest is the administration of parental care, which informs us of the fate of breeding events. By estimating reproductive fate from movement trajectories, our method helps bridge the gap between movement and individual fitness.

## List of abbreviations

GPS: Global Positioning System
CART: Classification And Regression Trees
MCMC: Markov Chain Monte Carlo
JAGS: Just Another Gibbs Sampler
GLM: Generalized Linear Model

## Declarations

### Ethics approval and consent to participate

Not applicable.

### Consent for publication

Not applicable.

### Availability of data and material

The GPS-tracking datasets generated and analyzed during the current study will be made publicly available on MoveBank. The datasets on nest locations and fates are available from the corresponding author on reasonable request.

### Authors’ contributions

S. Picardi conceived the idea. S. Picardi., B. J. S., and M. E. B. implemented the analyses. S. Picardi wrote the paper. M. B. and P. C. F. supervised research. J. G. C., D. R., L. S., S. Pirrello, and R. R. B. collected the data. All authors contributed to revisions.

## Acknowledgments

We thank D. Bear for providing data on known nest locations for wood storks in northern Florida. We thank anonymous reviewers, as well as J. Hightower and C. Poli, for their constructive criticism that greatly helped to improve previous drafts of the manuscript. The Mediterranean gull tracking project was carried out with the support of Ente Parco Delta del Po Emilia-Romagna, Reparto Carabinieri per la Biodiversità di Punta Marina, R. Nardelli, A. Andreotti, G. Meneghini, F. Spina. Lesser kestrel tracking project activities were carried out with the support of S. Podofillini, E. Fulco, P. Giglio, S. C. Pellegrino, M. Lorusso (Comune di Altamura), F. Parisi (Comune di Gravina in Puglia), and D. Ciampanella (project manager). Any use of trade, firm, or product names is for descriptive purposes only and does not imply endorsement by the U. S. Government.

