## Additional file 4 for "Analysis of movement recursions to detect reproductive events and estimate their fate in central place foragers"

### Sensitivity analysis on buffer size for breeding-site detection

After running an initial round of analysis with a 40 m buffer for all three species (wood storks, lesser kestrels, and Mediterranean gulls) and identifying optimal values for the revisitation parameters using CART, we performed a sensitivity analysis of nest-detection performance according to a range of possible buffer radii. We included buffers of size ranging from 10 to 100 m (this is a reasonable range for the behavior that we are targeting, i.e., returns to an exact location (nest) versus, for example, returns to a general area (foraging patch)). We calculated three performance metrics (positive predictive value, sensitivity, false negative rate) using each of these buffer sizes while keeping revisitation parameters constant. We limited our assessment to the three performance metrics that are verified using known nests (thus excluding false positive rate which is calculated using non-breeding individuals). We defined positive predictive value as the percentage of nest sites that were known among those we found for each species; sensitivity as the percentage of known nest sites that were identified; false negative rate as the percentage of known nest sites that we failed to identify.

We found an optimal buffer size of 30 m for wood storks, 60 m for lesser kestrels, and 20 m for Mediterranean gulls. Optimal buffer size is affected by both data resolution and behavior. Spatial resolution of the GPS data was approximately 17 m for wood storks and 10 m for the other two. Nonetheless, the optimal buffer for kestrels was larger than for gulls. This might depend on differences in terms of cover between kestrel and gull nests: the first nest in cavities, where the GPS signal might have been poor and affected spatial accuracy of fixes more so than for gulls, which nest in the open in salt pans in our study area. Additionally, behavior might have played a role, with kestrels possibly spending more time in the close proximity of the nest (but not in it) than gulls. Storks, like gulls, nest in the open where GPS signal should be good; however, the lower spatial resolution of the storks data resulted in a slightly larger optimal buffer size than for gulls.

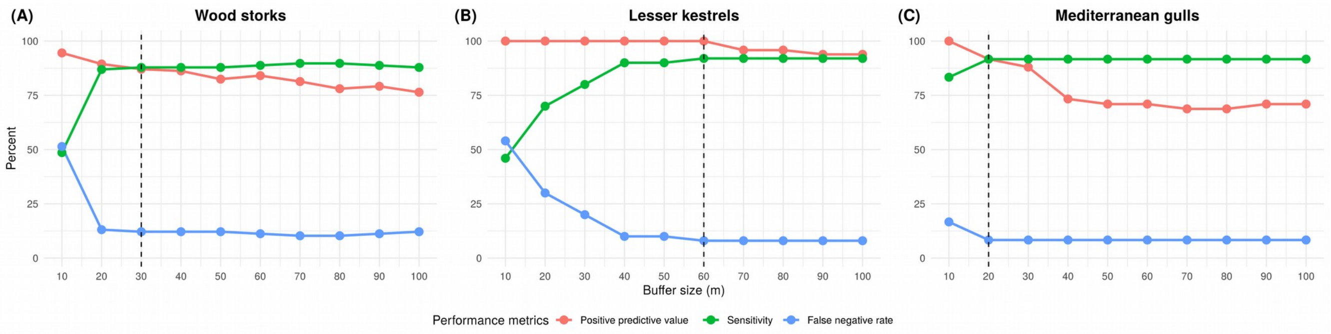


Figure 6 – **Results of sensitivity analysis on buffer size.** Nest-detection performance metrics as a function of buffer size for A) wood storks, B) lesser kestrels, and C) Mediterranean gulls. Positive predictive value in red, sensitivity in green, false negative rate in blue. The dashed vertical lines indicate the optimal value we chose for each species.
