## Additional file 3 for "Analysis of movement recursions to detect reproductive events and estimate their fate in central place foragers"

### Analysis of bird nesting from GPS data in R: the `nestR` package

**Simona Picardi**

**January 2019**


- Introduction
- TL;DR
- Data format
- Part I: Finding nest locations
  - Background
  - Introducing the `find_nests()` function
  - Step 1: Identifying recurrently visited locations
  - Step 2: Discriminating between nests and non-nests
  - Step 3: Identifying nests among revisited locations
- Part II: Estimating reproductive outcome
  - Background
  - Formatting nest revisitation histories
  - Nest survival model specification
  - Model fitting and results
- Case studies: applying the method to different species and data
- Conclusions


#### Introduction

The `nestR` package provides functions to locate nesting attempts and estimate their outcome from bird GPS-tracking data. Being able to estimate reproductive outcome from GPS data bridges the gap between movement and an important component of individual fitness.

This vignette presents the workflow of the `nestR` package. The workflow can be conceptually divided in two parts: first, the identification of nesting attempts along individual movement trajectories, and second, the estimation of the outcome of nesting attempts.

The following diagram summarizes the workflow and provides a roadmap for the reader:

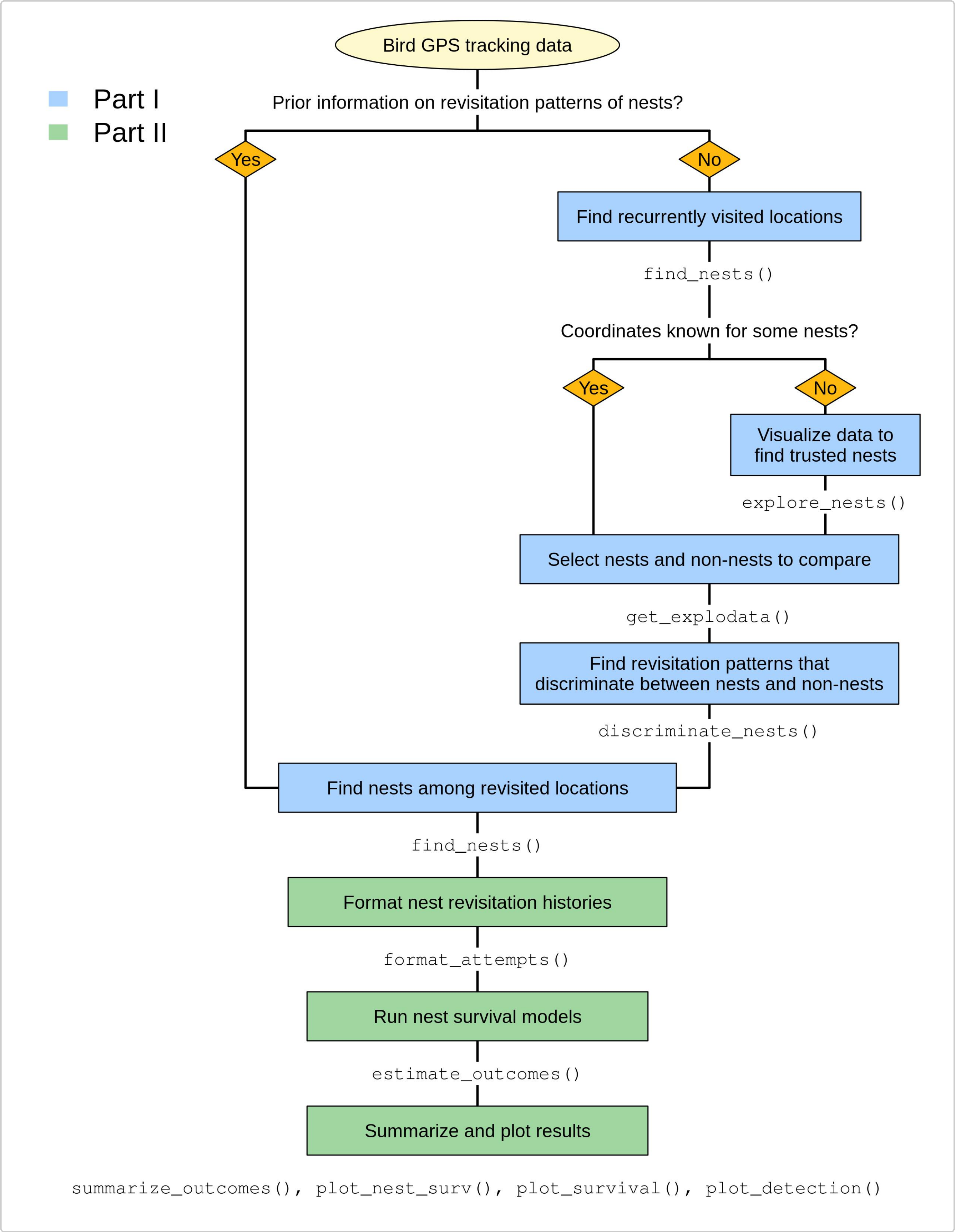


For illustration purposes, this vignette uses example datasets on three bird species: wood storks (*Mycteria americana*), lesser kestrels (*Falco naumanni*), and Mediterranean gulls (*Ichthyaetus*

*melanocephalus*). Each dataset includes real trajectories for two individual-years. All example datasets are available within the package.

## TL;DR

This section is meant to provide a concise and self-sufficient overview of the capabilities of the `nestR` package. Readers that want to get familiar with `nestR` without delving too deep into the technical aspects can stop at the end of this section. The next sections provide a more detailed guide to the usage of `nestR` functions.

- **Context –** The rationale behind the development of `nestR` is that bird movement patterns during nesting differ from those they exhibit when non nesting, and that this behavioral signal is observable in tracking data and can be used to infer the location of nests and the outcome of nesting attempts (see conceptual figure below). When birds are nesting, they behave as central place foragers, exhibiting back-and-forth movements to and from their nest while they provide food for their nestlings and themselves. Parents keep returning to their nest to feed their offspring until fledging. Repeatedly visited locations can be identified along movement trajectories, and patterns of revisitation can be used to determine whether a location that gets visited repeatedly is likely to be a nest. Histories of nest revisitation can then be used to estimate the outcome of nesting attempts. Specifically, we can estimate if an attempt was successful or not according to whether it lasted as long as the theoretical duration of a complete nesting cycle for the focal species. The underlying assumption is that the nest stops being revisited after an attempt fails, which is true for many bird species.

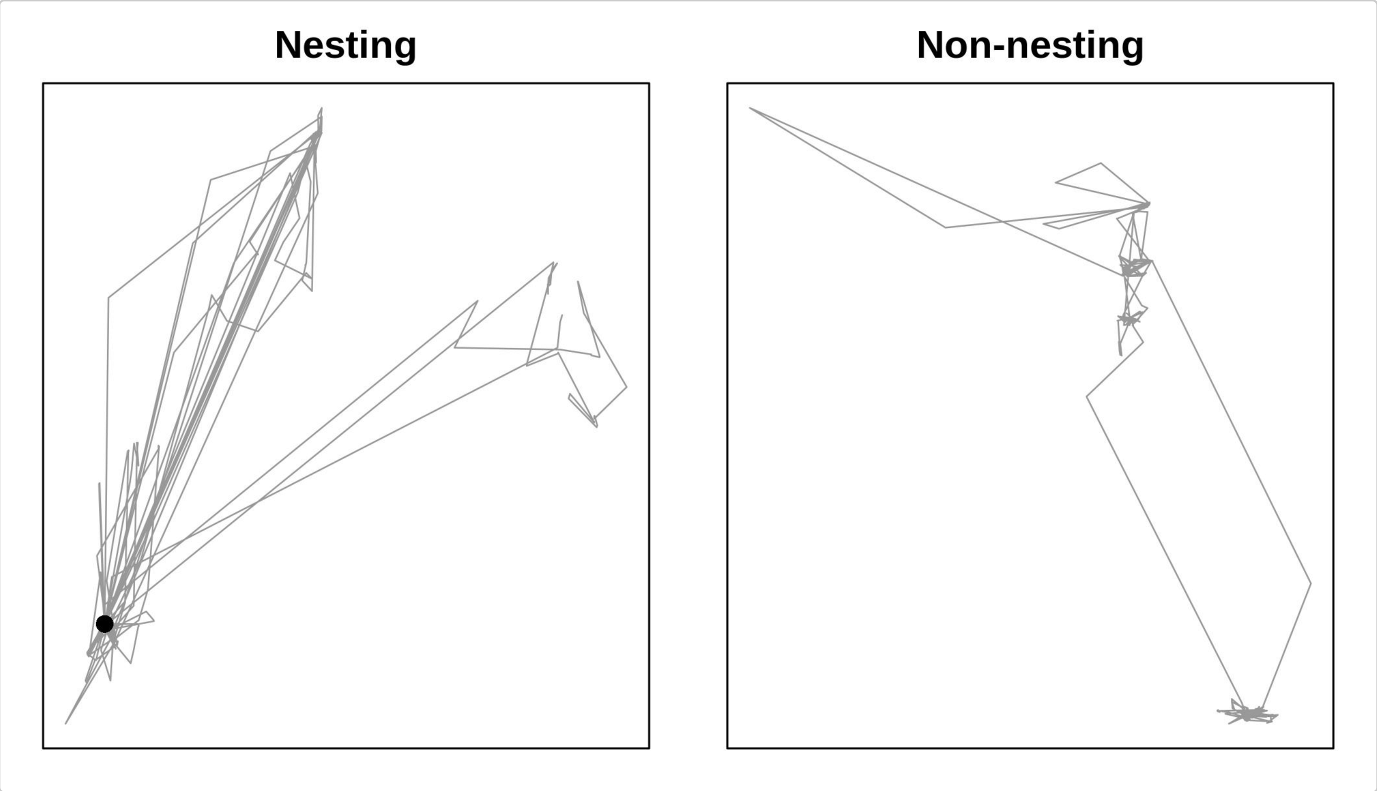


- **Is prior information on revisitation patterns of nest available? –** Nests are expected to be visited for longer stretches of consecutive days, more consistently, or for longer/more times during a day compared to other types of revisited locations. If prior information on revisitation patterns of nests is available, the user can skip directly to finding nests along bird trajectories using the `find_nests()` function. This function takes as input the tracking data and a series of species- and data-specific parameters that are used to describe revisitation patterns. The function filters nests among revisited locations based on the parameter values in input. However, it will often be the case that the user has only a vague idea of what revisitation patterns distinguish nests from non-nests, and making a decision on which parameter values to use will not be straightforward. In this case, data exploration tools in `nestR` can help, as illustrated in the following bullet points.

- **Find recurrently visited locations** – Conceptually, `find_nests()` can be used to find any type of revisited location. Using non-constraining parameter values as input in `find_nests()` allows the user to obtain a list of revisited locations that includes both nests and non-nests. Then, comparing revisitation patterns of nests and non-nests can inform the choice of parameter values to filter nests only among revisited locations.
- **Are coordinates known for some nests?** – Comparing nests and non-nests requires having prior information on the location of *some* nests. In the best-case scenario, this information is available to the user in the form of on-ground data (e.g., geographic coordinates of nest locations for a subset of the data). If this is not the case, an alternative is to visually explore the data and identify "trusted" nest locations (for example, because they fall within known colonies).
- **Visualize data to find trusted nests** – `nestR` includes an interactive visualization tool that can be used to identify "trusted" nests: `explore_nests()` launches a Shiny app that allows to interactively explore data on revisited locations by manipulating input parameters on the fly.
- **Select nests and non-nests to compare** – Based on the information on the location of true nests, the user can select a balanced sample of nests and non-nests to compare among revisited locations using the function `get_explodata()`.
- **Find revisitation patterns that discriminate between nests and non-nests** – Once the user has obtained a dataset of nests and non-nests, `discriminate_nests()` will run a CART (Classification and Regression Trees) algorithm to identify the set of filtering parameter values that best discriminate between the two. This can inform the choice of parameter values to input in `find_nests()`.
- **Find nests among revisited locations** – By inputting the parameters resulting from the previous step in `find_nests()`, the user can identify nests among revisited locations. The described workflow for finding nests is essentially a two-step feedback loop where parameter values used to find nests are recursively tuned: first, running `find_nests()` with non-constraining parameters identifies any revisited location along a bird's trajectory; then, using exploratory tools, the user finds a set of parameter values to tell nests apart from non-nests; finally, running `find_nests()` again, this time with the parameter values obtained from the exploratory analysis, narrows down the selection to nests only.
- **Format nest revisitation histories** – Once nests are identified, the user can analyze patterns of revisitation to infer whether a nesting attempt was completed or not. Conveniently, together with the set of nests found, `find_nests()` outputs a history of revisitation for each of them. The function `format_attempts()` formats revisitation histories into a matrix of daily fixes and a matrix of daily visits for each attempt. This will allow us to estimate the probability of having missed nest visits on days where they were not observed.
- **Run nest survival models** – `estimate_outcomes()` fits a Bayesian hierarchical model to the histories of nest revisitation to estimate daily nest survival proability until the end of the attempt. The function supports model structures where both the probability of detecting a nest visit and the probability of survival can be either constant or vary in time.
- **Summarize and plot results** – Finally, `nestR` includes functions to summarize and plot results of the nest survival models both at the population and at the individual level (`summarize_outcomes()`, `plot_nest_surv()`, `plot_detection()`, `plot_survival()`).

#### Data format

Functions in the `nestR` package require GPS data to be formatted as a `data.frame` including four columns:

- `burst` - this is a unique identifier of each individual-year. We recommend the user split individual data into bursts on a date that does not overlap with nesting activities. The full breeding cycle of an individual on a given year needs to be contained within a single burst.
- `date` - this is a date-time object of class `POSIXct`.
- `long` and `lat` - the longitude and latitude of the locations in decimal degrees.

Additional columns will not interfere with the functioning of the package, as long as these four fields are present and named with the terms above.

We recommend using the package `adehabitatLT` to cut the raw trajectories into bursts and get the data in the required format for analysis.

Let's load `nestR` and take a look at the wood stork dataset, for example:

```
# Load the package
library(nestR)

# Load wood stork dataset
data(woodstorks)

# Take a look
head(woodstorks)
#>          burst                date      long      lat
#> 1 1134370-2013 2012-10-02 07:00:00 -79.23650 33.23600
#> 2 1134370-2013 2012-10-02 08:00:00 -79.22867 33.24000
#> 3 1134370-2013 2012-10-02 09:00:00 -79.22983 33.24067
#> 4 1134370-2013 2012-10-02 10:00:00 -79.23133 33.23583
#> 5 1134370-2013 2012-10-02 11:00:00 -79.23817 33.22667
#> 6 1134370-2013 2012-10-02 12:00:00 -79.23800 33.22667

# Data structure
str(woodstorks)
#> 'data.frame':    9488 obs. of  4 variables:
#>  $ burst: chr  "1134370-2013" "1134370-2013" "1134370-2013" "1134370-2013" ...
#>  $ date : POSIXct, format: "2012-10-02 07:00:00" "2012-10-02 08:00:00" ...
#>  $ long : num  -79.2 -79.2 -79.2 -79.2 -79.2 ...
#>  $ lat  : num  33.2 33.2 33.2 33.2 33.2 ...
#>  - attr(*, "spec")=
#>   .. cols(
#>   ..   X1 = col_double(),
#>   ..   X1_1 = col_double(),
#>   ..   X = col_double(),
#>   ..   date = col_datetime(format = ""),
#>   ..   id = col_double(),
#>   ..   burst = col_character(),
#>   ..   pkey = col_double(),
#>   ..   year = col_double(),
#>   ..   id_nestyr = col_character(),
#>   ..   long = col_double(),
#>   ..   lat = col_double()
#>   .. )
```

### Part I: Finding nest locations

#### Background

When birds are nesting, they exhibit specific movement patterns that can allow us to identify the location of their nest. Specifically, birds perform repeated back-and-forth movements to and from their nest while they provide food for their nestlings and themselves. Repeatedly visited locations can be identified along movement trajectories, and patterns of revisitation can be used to determine whether a location that gets visited repeatedly is likely to be a nest.

The process of identifying nesting locations from GPS data relies on two components: knowledge of the biology of the study species and characteristics of the data at hand. Both of these aspects are fundamental in driving the patterns we observe in movement trajectories and require careful consideration by the user.

The datasets included in the `nestR` package differ both in terms of species biology and data characteristics. The following table summarizes the main differences:

| | Data characteristics | | | | Species biology | | | |
|---|---|---|---|---|---|---|---|---|
| **Dataset** | **Time resolution** | **Spatial resolution** | **Fix failure rate** | **Tagged at** | **Nesting cycle** | **Nesting season** | **Behavioral dimorphism** | **Repeated attempts** |
| woodstorks | 1 hour | 18 meters | High | Birth/non-breeding | 110 days | Nov-Aug (varies with latitude) | No | Yes |
| kestrels | 15 minutes | ~10 meters | Low | Early chick-rearing | 60 days | Apr-Jul | Yes | No |
| gulls | 15 minutes | ~10 meters | Low | Incubation | 60 days | Apr-Jul | No | Yes |

Throughout this vignette, we will refer back to specific features of each dataset and illustrate how these can and need to be accounted for when using functions in the `nestR` package.

#### Introducing the `find_nests()` function

The central function for the first part of the `nestR` workflow is `find_nests()`. This function identifies potential nests based on patterns of location revisitation. It takes as input the GPS data and returns any revisited locations that match a set of user-defined parameter values that characterize revisitation patterns, along with their revisitation history. Input parameters can be conceptually organized in three groups:

- Parameters related to basic information on the biology of the species;
- Parameters related to data characteristics;
- Parameters used for discriminating nests from other repeatedly visited locations that are not nests.

Typically, while prior information on the first two is available to the user, this might not be the case for the third group of parameters. Therefore, our recommended workflow for finding nests involves the following steps:

1. Identifiying any recurrently visited locations, including potential nests, along individual movement trajectories;
2. Comparing revisitation patterns at nests versus non-nests and establish a way to tell them apart;
3. Based on what found in step 2, identifying nests among revisited locations.

The function `find_nests()` will be used to tackle steps 1 and 3.

To accomplish step 2, some information on the location of nests for a subset of the data is necessary. In an ideal situation, the user has access to prior information on the location of nests for some individual-years in the dataset. Otherwise, a viable alternative is for the user to identify “trusted nests” among revisited locations through visual inspection, for example because they fall within known nesting colonies. Both cases will be illustrated later on.

Here’s the full list of arguments of the function `find_nests()`:

- `gps_data`
- `sea_start`
- `sea_end`
- `nest_cycle`
- `buffer`
- `min_pts`
- `min_d_fix`
- `min_consec`
- `min_top_att`
- `min_days_att`
- `discard_overlapping`

The first argument is the GPS data, formatted as described in Data format.

The next three arguments describe basic information of the biology of the species:

- `sea_start` and `sea_end` delimit the start and end of the breeding season for the focal species. Restricting the scope of the analysis to the breeding season only is especially important for reducing computation time and to avoid memory issues when running `find_nests()`.
- `nest_cycle` is the duration (number of days) of a complete nesting attempt for the focal species. Typically, it counts how many days go by, on average, between the moment an individual starts building its nest and the moment its chicks fledge.

The next three arguments are related to data characteristics:

- `buffer` defines the spatial scale at which revisitation patterns will be calculated. Returns to a location are defined as returns to a circular area of radius = `buffer` (in meters). The use of this parameter is meant to account for the spatial scattering of GPS points around a nest due to both behavior (sometimes the tag will happen to record a point when the bird was in the proximity of the nest, possibly arriving or departing, and not exactly on it) and GPS error. The value of `buffer` needs to be set to a number greater or equal to the spatial resolution of the GPS data.
- Revisitation stats will be computed on each area of size `buffer` that is visited multiple times. To speed up calculations, the user can discard isolated points from the get-go and not bother calculating revisitation stats on those. The parameter `min_pts` allows the user to specify a minimum number of points that need to fall within a point’s buffer for it to be retained in the calculation of revisitation stats.
- Some of the revisitation stats included in the output of `find_nests()` are based on the number of consecutive days a location is visited. For tags that have a high fix failure rate and a relatively low temporal resolution, it is easy to miss a nest visit even though it happened. This can break an otherwise continuous stretch of days where visits were recorded. The argument `min_d_fix` allows

the user to counteract the effect of missed visits, by setting a minimum number of fixes that need to be available in a day when no visit was detected for that day to be truly counted as non visited. In the next section, we provide an example of when this is especially needed.

The remaining arguments are used for filtering locations based on revisitation patterns:

- Revisitation patterns include the number of consecutive days a location is visited, the percent of fixes at the location on the day with maximum attendance, and the percent of days a location is visited between the first and last visit. The user can set minimum values for each of these using `min_consec`, `min_top_att`, and `min_days_att`, respectively. Any location for which revisitation patterns do not exceed these user-defined thresholds will not be returned. Prior information regarding these parameters is likely not available to the user, and finding the best set of values to specify for these arguments will be the focus of Step 2.
- Finally, `discard_overlapping` specifies whether the function should return all the revisited locations that match the specified criteria, or avoid returning locations for which the time ranges of revisits are overlapping. Assuming that a bird cannot nest in two places at the same time, if `discard_overlapping` is set to `TRUE`, only the most likely nest location is returned among temporally overlapping ones. However, we will later illustrate the case where it is useful to set this argument to `FALSE` instead.

The output of `find_nests()` is a list with two elements: `nests` and `visits`. The first element, `nests`, is a `data.frame` including any revisited locations that match the criteria specified in the function arguments, along with their revisitation stats. For example, here is a preview of an output (the following paragraphs will illustrate in detail how to get there):

```
#>          burst loc_id    long      lat first_date  last_date attempt_start
#> 1 1134370-2013   2170 -80.851 25.46283 2013-02-24 2013-05-02    2013-02-24
#> 2  721290-2010    594 -81.645 30.40483 2009-11-26 2010-08-16    2010-03-24
#>   attempt_end tot_vis days_vis consec_days perc_days_vis perc_top_vis
#> 1  2013-05-02     589       61          37         89.71          100
#> 2  2010-07-11    1066      143          37         54.17          100
```

Each row of the `data.frame` corresponds to one revisited location. The columns are:

- `burst`, the burst id of the individual-year;
- `loc_id`, the unique identifier of the location at the center of the buffer;
- `long` and `lat`, its coordinates;
- `first_date` and `last_date`, the days when the location was first and last visited;
- `attempt_start` and `attempt_end`, the estimated start and end dates of the nesting attempt;
- `tot_vis`, the total number of visits (fixes);
- `days_vis`, the total number of days when the location was visited;
- `consec_days`, the duration of the longest stretch of consecutive days when the location was visited;
- `perc_days_vis`, the percentage of days when the location was visited between the first and last visits;
- `perc_top_vis`, the percentage of fixes at the location on the day with maximum attendance.

Note the direct correspondence between the last three parameters and the three filtering arguments in input (`min_consec`, `perc_days_vis`, and `perc_top_vis`).

The days of first and last visit do not necessarily correspond to the start and end of a nesting attempt. If a bird visits the nest location before actually starting to nest, or returns to the nest after having completed the attempt, using the days of first and last visits as temporal limits of the attempt is going to be misleading. Since estimating the start and end dates of an attempt correctly will be critical for successfully estimating its outcome, `find_nests()` takes into account the possibility of visits outside of the attempt and cuts them out. The implementation is based on the value of `min_consec`: the attempt is estimated to start at the first occurrence of a stretch of at least `min_consec` consecutive days. Then, if the nest was visited for longer than the duration of a complete nesting attempt after that, the end date gets set at `attempt_start` + `nest_cycle` (that is, the latest date possible given the start date). If the last visit occurs before `attempt_start` + `nest_cycle`, then `attempt_end` is set to `last_date`. We stress the importance of being aware that the value of `min_consec` has an effect on the estimation of the temporal limits of nesting attempts.

The second element of the list that `find_nests()` returns as output is a `data.frame` named `visits`: this is essentially an updated version of the GPS data provided in input, with an additional column that flags any records taken at a nest with the location ID of that nest. Locations away from the nest are assigned zeroes. Therefore, `visits` contains the revisitation history of all nests listed in `nests`. This is what it looks like:

```
#>            burst                date      long      lat loc_id
#> 1  1134370-2013 2012-11-12 14:00:00 -80.43233 25.29583      0
#> 2  1134370-2013 2012-11-12 15:00:00 -80.57600 25.44033      0
#> 3  1134370-2013 2012-11-12 16:00:00 -80.43200 25.37900      0
#> 4  1134370-2013 2012-11-12 17:00:00 -80.41683 25.37350      0
#> 5  1134370-2013 2012-11-12 18:00:00 -80.41600 25.37250    644
#> 6  1134370-2013 2012-11-12 19:00:00 -80.41617 25.37283    644
#> 7  1134370-2013 2012-11-12 20:00:00 -80.41617 25.37283    644
#> 8  1134370-2013 2012-11-12 21:00:00 -80.41617 25.37267    644
#> 9  1134370-2013 2012-11-13 06:00:00 -80.41617 25.37267    644
#> 10 1134370-2013 2012-11-13 07:00:00 -80.41667 25.37333      0
#> 11 1134370-2013 2012-11-13 08:00:00 -80.40967 25.36783      0
#> 12 1134370-2013 2012-11-13 09:00:00 -80.40967 25.36783      0
#> 13 1134370-2013 2012-11-13 10:00:00 -80.41000 25.36783      0
```

#### Step 1: Identifying recurrently visited locations

Animals exhibit recursive movement patterns for a variety of reasons. In the case of birds, locations that get visited recurrently can include nests as well as roosting sites, favorite foraging spots, and so on. However, patterns of revisitation likely differ for locations that a bird visits for different reasons. Nests are expected to be visited for longer stretches of consecutive days, more consistently, or for longer/more times during a day compared to other types of revisited locations. But *how much* more? `nestR` provides tools to figure that out.

As a first step, we suggest using the `find_nests()` function to screen the data to identify recurrently visited locations, regardless of whether they are nests or not. To do so, the user can specify low values for the arguments involved with the filtering, so that the constraints applied are loose.

Here is an example with wood storks. Wood storks nest at different times of the year in different parts of their range, so the first challenge with this species is that there isn't a well-defined nesting season. For example, wood storks in southern Florida usually breed between January and May, but those that nest in some areas farther north can start as late as March or April. Overall, we cannot exclude the possibility of observing nesting events any time between November and August. Therefore, we set `sea_start` and `sea_end` to November 1st and August 31st, respectively. The time required for an individual wood stork to complete its nesting cycle is 110 days. We set `nest_cycle` to this value.

The spatial accuracy of the GPS data for wood storks is 18 meters according to the tag user manual. We set the value of `buffer` to 40 meters to allow some extra room for spatial scattering of points around a central location. In general, we found that buffer sizes between 20 and 50 meters return comparable results in our three case studies. We encourage the user to explore results obtained with different buffer sizes and possibly fine-tune the value to one that well captures the spatial scale of recursive movements given the species and data at hand. We set `min_pts` to 2 points within the buffer to avoid calculating revisitation patterns for points that are relatively isolated. This is still a very low, conservative value that nevertheless substantially helps reduce computation time.

We set `min_d_fix` to 5, meaning that any day with no visit does not interrupt a stretch of consecutive days if it has fewer than 5 fixes. This is especially important for wood storks because the time interval between fixes is 1 hour, the tags are solar powered and therefore only take fixes during the daytime (maximum 16 fixes per day), and the fix failure rate is high. These factors combined mean that often there are only a handful of fixes collected on a day, and the probability of missing nest visits is high. If a stretch of consecutive days is interrupted by one or more days with no recorded visit but that only include 1 to 4 fixes, we do not have enough information to truly determine if the nest was not visited, and we assume that a visit was likely missed instead. As we will mention later, this is not as much of a problem in the case of both kestrels and gulls, because those tags were set to collect data every 15 minutes and have a low fix failure rate, so that the probability of missing visits to the nest is lower. There is no obvious rule to pick an adequate value for `min_d_fix`. We recommend that the user explores the data to assess fix failure rate and chooses a value of `min_d_fix` that they are comfortable with given temporal resolution and fix failure rate.

We set the three filtering arguments to low values, meaning that the constraints we enforce are as loose as they can be. We set `min_consec` to 2 days, and `min_top_att` and `min_days_att` to 1%. This will return any location that is visited for at least 2 consecutive days, regardless of how often it is visited between the first and last visit and regardless of how much is it visited for on the day with the most visits. We also set `discard_overlapping` to `FALSE` so that the function will not discard locations that are visited simultaneously to a likely nest. This will be useful for Step 2.

Fair warning: depending on the amount of data, `find_nests()` can be computationally intensive and take some time to run (typically <1 minute per burst).

```
ws_output_1 <- find_nests(gps_data = woodstorks,
                          sea_start = "11-01",
                          sea_end = "08-31",
                          nest_cycle = 110,
                          buffer = 40,
                          min_pts = 2,
                          min_d_fix = 5,
                          min_consec = 2,
                          min_top_att = 1,
                          min_days_att = 1,
                          discard_overlapping = FALSE)
```

Here is what the output looks like:

```
head(ws_output_1$nests)
```

```
#>           burst loc_id      long      lat first_date  last_date
#> 1 1134370-2013   2170 -80.85100 25.46283 2013-02-24 2013-05-02
#> 2 1134370-2013   3270 -79.39017 33.14267 2013-05-12 2013-06-03
#> 3 1134370-2013   1023 -80.57167 25.48183 2012-11-27 2012-12-20
#> 4 1134370-2013   2364 -80.84683 25.46033 2013-02-02 2013-04-18
#> 5 1134370-2013   1391 -80.54367 25.24817 2012-12-23 2013-01-17
#> 6 1134370-2013   1317 -80.54383 25.24900 2012-12-23 2013-01-19
#>   attempt_start attempt_end tot_vis days_vis consec_days perc_days_vis
#> 1    2013-02-24  2013-05-02     589       61          37         89.71
```

```
#> 2    2013-05-12  2013-06-03     219       22          20         95.65
#> 3    2012-12-07  2012-12-20      97       15          14         62.50
#> 4    2013-03-08  2013-04-18      87       37          17         48.68
#> 5    2012-12-23  2013-01-17      60       17           4         65.38
#> 6    2012-12-23  2013-01-19      54       16           8         57.14
#>   perc_top_vis
#> 1       100.00
#> 2       100.00
#> 3        87.50
#> 4        35.71
#> 5        83.33
#> 6        80.00
```

Let's check how many revisited locations were returned with this set of parameters for each of the two individual-years:

```
table(ws_output_1$nests$burst)
#>
#> 1134370-2013  721290-2010
#>          140           86
```

Results are automatically sorted by `tot_vis` within each burst, so for each individual the location that was visited the most is at the top of the list.

#### Step 2: Discriminating between nests and non-nests

The set of revisited locations obtained as output of the screening in Step 1 likely includes both nests and non-nests. The objective of this section is to identify the best set of parameter values to discriminate between these. Specifically, the parameters that we want to tune are those that describe patterns of revisitation: `min_consec`, `min_top_att`, and `min_days_att`. To inform our choice of values for these parameters, we want to compare the values obtained in output for nests and non-nests for `consec_days`, `perc_top_vis`, and `perc_days_vis`.

The function `get_explodata()` automates the process of selecting nests and non-nests to compare. Given the output of `find_nests()` and information on known nest locations, the function extracts the true nest and another location that is not a nest from the set of revisited locations. To illustrate how the function works, let's first step through the process manually on the two individuals in the wood stork example dataset.

##### Case A: coordinates of true nest are known

For one of the two wood storks, the location of the nest is known. This individual bred at a colony located at the Jacksonville Zoo, and we are lucky enough to have the GPS coordinates of its nest. For this individual, thus, we can identify the true nest among the set of revisited locations by comparing coordinates to those of the known nest. We can then select another location among the remaining to serve as a non-nest counterpart, and compare how values of the parameters describing revisitation patterns differ between the two.

Let's look at the first few revisited locations identified for this individual in the previous section:

```
head(ws_output_1$nests %>% filter(burst == "721290-2010"))
```

```
#>         burst loc_id      long      lat first_date  last_date
#> 1 721290-2010    594 -81.64500 30.40483 2009-11-26 2010-08-16
#> 2 721290-2010    771 -81.72133 30.07050 2010-02-06 2010-03-21
#> 3 721290-2010   3116 -81.58433 30.39700 2010-07-20 2010-08-29
#> 4 721290-2010   1246 -81.64517 30.40517 2010-01-22 2010-08-08
#> 5 721290-2010   2941 -81.55367 30.36583 2010-05-26 2010-08-15
#> 6 721290-2010   1770 -81.61717 30.40450 2010-04-20 2010-05-23
#>   attempt_start attempt_end tot_vis days_vis consec_days perc_days_vis
#> 1    2010-03-24  2010-07-11    1066      143          37         54.17
#> 2    2010-02-06  2010-03-21      76       26          10         59.09
#> 3    2010-08-10  2010-08-29      72       17           5         41.46
#> 4    2010-01-24  2010-05-13      60       32           4         16.08
#> 5    2010-07-18  2010-08-15      60       18           5         21.95
#> 6    2010-04-29  2010-05-23      42       13           4         38.24
#>   perc_top_vis
#> 1       100.00
#> 2        38.46
#> 3        78.57
#> 4        42.86
#> 5        91.67
#> 6        61.54
```

The first location was visited a total of 1066 times, for 143 days (of which 37 consecutive). The bird went back to the location on 54.17% of days between the day of first and last visit. On the day the bird spent the most time at the location, 100% of the fixes were at the location. This seems like a good candidate nest. Let's compare its coordinates to the ones of the known nest and see if they match.

```
data(jax_known_nest)

jax_known_nest
#>         burst      long      lat
#> 1 721290-2010 -81.64525 30.40485
```

The coordinates match until the third (longitude) and fourth (latitude) decimal places. Let's see what distance that corresponds to:

```
coords_cand <- ws_output_1$nests %>%
  filter(burst == "721290-2010") %>%
  slice(1) %>%
  select(long, lat)

coords_known <- jax_known_nest %>%
  select(long, lat)

geosphere::distGeo(coords_cand, coords_known)
#> [1] 24.0195
```

The points are about 24 meters apart. That is well within what expected due to both GPS error and behavioral patterns affecting the signal we get in the data, and well within the 40 meters buffer we used to calculate revisits. We can confidently confirm that that is the nest.

Now, we want to select a non-nest among the other revisited locations. We want to pick a location that we are confident is not a nest but that gets as close as possible to the revisitation parameters of a true nest. This will give us the most power to discern nests from non-nests.

One way of going about this task would be to select the second most visited location in the set. But how can we be sure that the second most visited location is not also a nest, maybe corresponding to a second nesting attempt?

Some species are known to be able to nest twice in the same breeding season if they start the first clutch early enough or if their first attempt fails. In this case, one way to be sure that the location we select is truly a non-nest is to choose one that temporally overlaps with the true nesting attempt at the known location. This is where having set `discard_overlapping = FALSE` in `find_nests()` comes in handy, as it made sure that locations visited simultaneously to the true nest were still retained in our output.

In the case of the Jacksonville stork, the nesting attempt at the true nest was estimated to start on March 24$^{th}$, 2010, and end on July 11$^{th}$, 2010. The second and third locations were both visited in a time range that does not overlap with that interval. The first instance of a location that was visited simultaneously with the true nest is the fourth location in the set. Since wood storks are able to have repeated nesting attempts, we can conservatively choose the fourth location as a non-nest to compare the nest to, rather than the second.

The function `get_explodata()` automates the process we described so far. It takes as input the list of candidate nests output by `find_nests()`, and a data frame of coordinates for the known nests (`known_coords`). The argument `pick_overlapping` allows the user to decide whether the criterion of temporal overlap to a true nest gets applied or not when selecting a non-nest. For species that do not re-nest, this might not be necessary. The function is also designed to handle two possible sources of error that could arise from automating the process instead of supervising it:

1. The true nest location is not in the output. This can happen if, for example, gaps in the data due to tag malfunctioning led to missing most of the visits to the nest.
2. Multiple locations surrounding the nest were selected as candidate nests, and the closest one to the nest in terms of linear distance is not the one with the highest revisitation parameters.

Selecting the true nest as simply the closest location to the known nest is prone to both of these error sources. The argument `buffer` in the function `get_explodata()` helps solve both of these issues. The function selects the true nest as the top visited location among those that fall within a `buffer` distance from the known nest. If there is none, nothing is returned. If multiple locations are available within the buffer, the most visited one, not the closest one, gets selected. If the coordinates in `known_coords` represent the actual location of the nest, we recommend setting `buffer` in `get_explodata()` to the same value used for the argument `buffer` in `find_nests()`. In cases where, for example, a single pair of coordinates is available for an entire colony rather than for the exact location of a nest within it, the value of `buffer` can be increased to match the spatial extent of the colony to ensure the true nest does not fall outside of the buffer.

Here is the output of `get_explodata()` for the Jacksonville stork:

```
output_stork1 <- ws_output_1$nests %>%
  filter(burst == "721290-2010")

(explodata_stork1 <- get_explodata(candidate_nests = output_stork1,
                                   known_coords = jax_known_nest,
                                   buffer = 40,
                                   pick_overlapping = TRUE))
#>         burst loc_id      long      lat first_date  last_date
#> 1 721290-2010    594 -81.64500 30.40483 2009-11-26 2010-08-16

#> 2 721290-2010   1246 -81.64517 30.40517 2010-01-22 2010-08-08
```

```
#>   attempt_start attempt_end tot_vis days_vis consec_days perc_days_vis
#> 1    2010-03-24  2010-07-11    1066      143          37         54.17
#> 2    2010-01-24  2010-05-13      60       32           4         16.08
#>   perc_top_vis nest
#> 1       100.00  yes
#> 2        42.86   no
```

The result of the automated process matches our expectations.

##### Case B: coordinates of true nest are unknown

For the second individual in the wood stork example dataset, no prior information is available on the location of the nest. However, we can identify the likely location of the nest by visually inspecting revisited locations in the output of `find_nests()`.

A useful tool available within `nestR` for visual exploration of the data is available with the function `explore_nests()`. This function launches a Shiny app that allows the user to interactively explore the results of `find_nests()` by visualizing them on a map. The app can be opened with the RStudio browser or any other web browser. Input parameters taken as arguments by `find_nests()` can be manipulated on the spot and the corresponding results are displayed over a satellite imagery basemap.

By launching `explore_nests()` on the wood stork data, selecting the burst 1134370-2013, and inputting the same parameters we used when running `find_nests()`, we notice that the first location of the set (the most visited one), falls right on the Rookery Branch colony in Everglades National Park. This, together with the fact that the location was visited a total of 589 times, for 61 days (of which 37 consecutive), 89.71% of the days between the first and last visit, and 100% of the time on the day with maximum attendance, makes us confident that the location represents a true nest.

```
explore_nests(woodstorks)
```

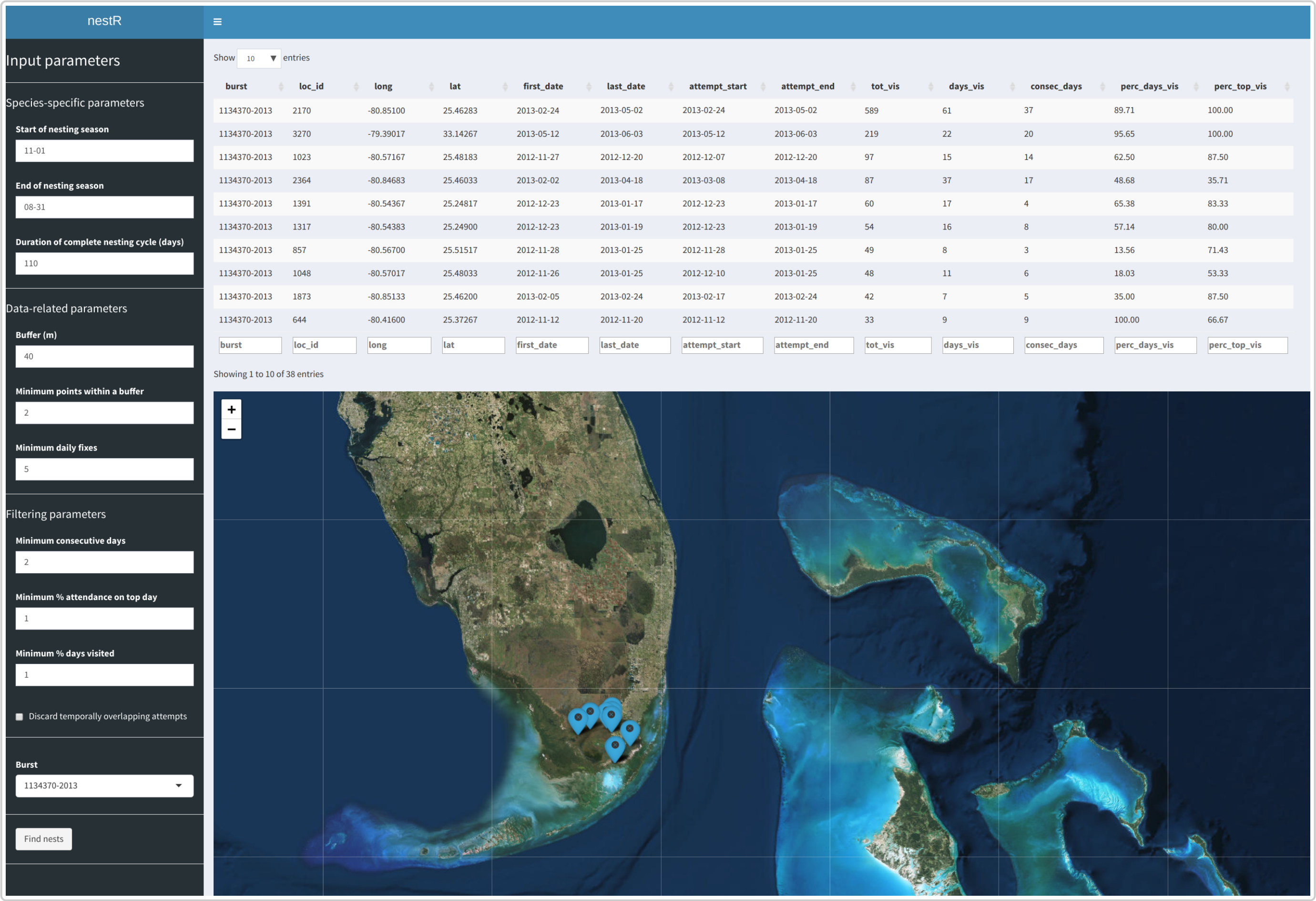


```
ws_output_1$nests %>%
  filter(burst == "1134370-2013") %>%
  slice(1)
#>          burst loc_id    long      lat first_date  last_date attempt_start
#> 1 1134370-2013   2170 -80.851 25.46283 2013-02-24 2013-05-02    2013-02-24
#>   attempt_end tot_vis days_vis consec_days perc_days_vis perc_top_vis
#> 1  2013-05-02     589       61          37         89.71          100
```

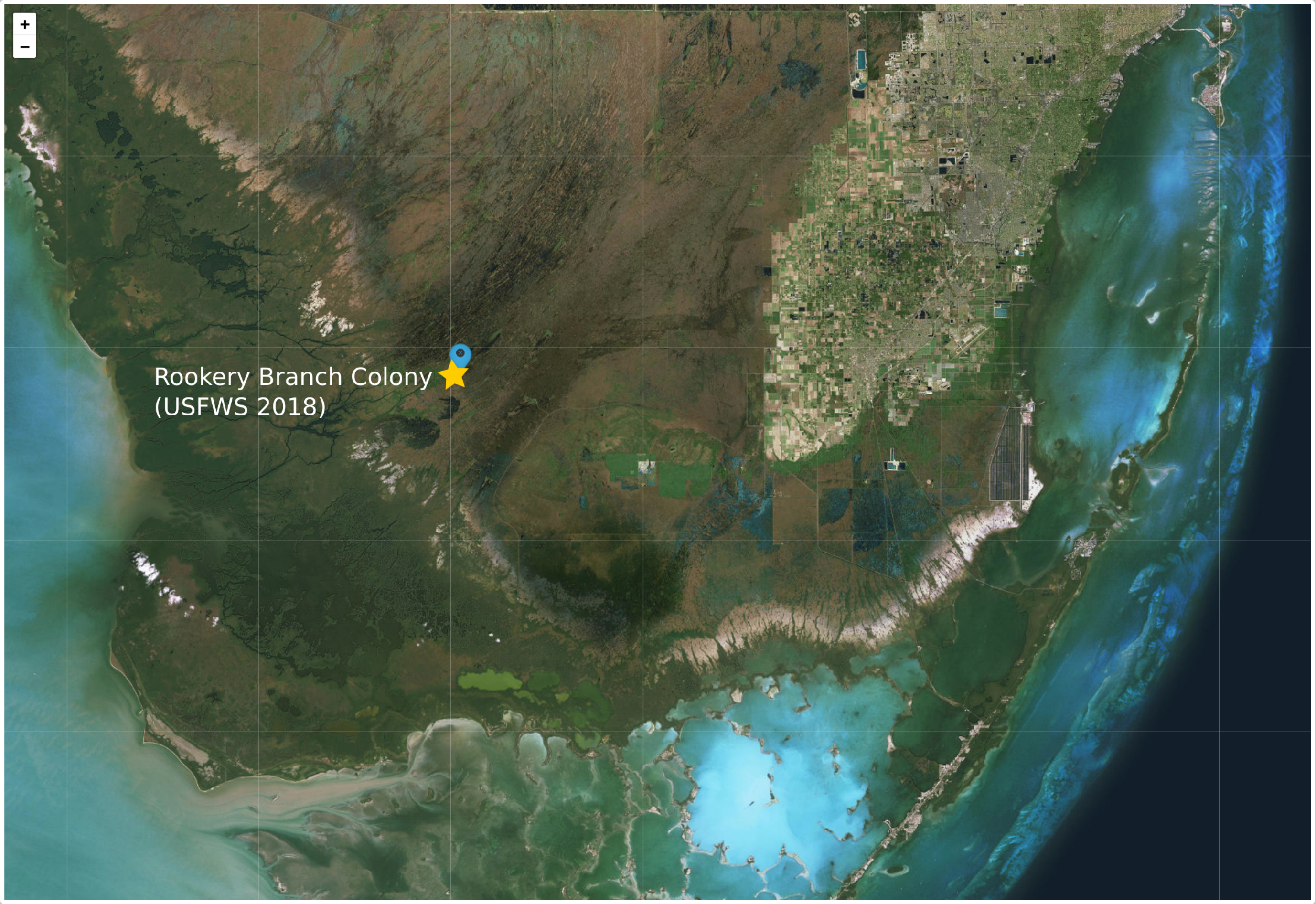


The function `get_explodata()` allows the user to input information on true nests in an alternative way that works well in this case: instead of providing the nest coordinates by passing `known_coords` to the function, the user can provide the location ID instead, by using the argument `known_ids`. We can thus run `get_explodata()` on wood stork 1134370-2013 by specifying the location ID of the point we trust to be a nest:

```
output_stork2 <- ws_output_1$nests %>%
  filter(burst == "1134370-2013")

id_known <- data.frame(burst = "1134370-2013",
                       loc_id = 2170)

(explodata_stork2 <- get_explodata(candidate_nests = output_stork2,
                                   known_ids = id_known,
                                   pick_overlapping = TRUE))
#>          burst loc_id      long      lat first_date  last_date
#> 1 1134370-2013   2170 -80.85100 25.46283 2013-02-24 2013-05-02
#> 2 1134370-2013   2364 -80.84683 25.46033 2013-02-02 2013-04-18
#>   attempt_start attempt_end tot_vis days_vis consec_days perc_days_vis
#> 1    2013-02-24  2013-05-02     589       61          37         89.71
#> 2    2013-03-08  2013-04-18      87       37          17         48.68
#>   perc_top_vis nest
#> 1       100.00  yes
#> 2        35.71   no
```

##### Find set of parameter values to tell apart nests and non-nests

Once we have a dataset of nests and non-nests, we can compare values of revisitation parameters between those, and use that to inform the choice of parameter values to filter nests among revisited locations.

Several possible approaches are available to tackle this objective. Here, we present one possible implementation, based on Classification and Regression Trees (CART). Functions to run a customized CART algorithm on the revisited location data are available within `nestR`, but we encourage the user to explore other implementation options or other analytical tools that may also be appropriate for the task.

To run the CART, we are going to use a larger version of the exploratory dataset we built in the previous step for 2 wood storks. This is a simulated dataset including 200 bursts with one nest and one non-nest each.

```
data(explodata_ws)
head(explodata_ws)
#>     burst loc_id      long      lat first_date   last_dat attempt_start
#> 1    B100     27 -80.70704 31.49420 2009-11-30 2010-07-12    2010-01-06
#> 101  B100    166 -80.60423 31.51261 2010-01-05 2010-06-15    2010-03-22
#> 2    B101     37 -79.84679 26.35810 2013-02-10 2013-08-09    2013-03-06
#> 102  B101    135 -79.67794 26.64990 2013-04-19 2013-01-17    2013-05-03
#> 3    B102     57 -80.73609 31.60196 2009-09-28 2010-08-12    2010-04-07
#> 103  B102    127 -80.48651 31.34487 2010-02-18 2010-09-27    2009-12-12
#>     attempt_end tot_vis days_vis consec_days perc_days_vis perc_top_vis
#> 1    2010-07-24     845      119          58     66.230057     86.58949
#> 101  2010-02-25     151       41          14     14.657008     66.79314
#> 2    2013-03-07     893       83          39     74.805390     99.54293
#> 102  2013-07-24      82       35          13     -6.065098     63.56319
#> 3    2010-09-24     780      120          19     83.416199    100.00000
#> 103  2010-02-10      59       17          10     64.774015     17.93633
#>     nest
#> 1    yes
#> 101   no
#> 2    yes
#> 102   no
#> 3    yes
#> 103   no
```

We are going to use half of the data to train the CART and the other half to test it. Classification accuracy should improve with the size of the training dataset. In cases where data availability is limited, the choice of how much data to use to train versus test the CART is based on a trade-off between classification accuracy and the ability to cross-validate the tree and estimate error rates.

```
(ws_cart <- discriminate_nests(explodata = explodata_ws, train_frac = 0.5))
```

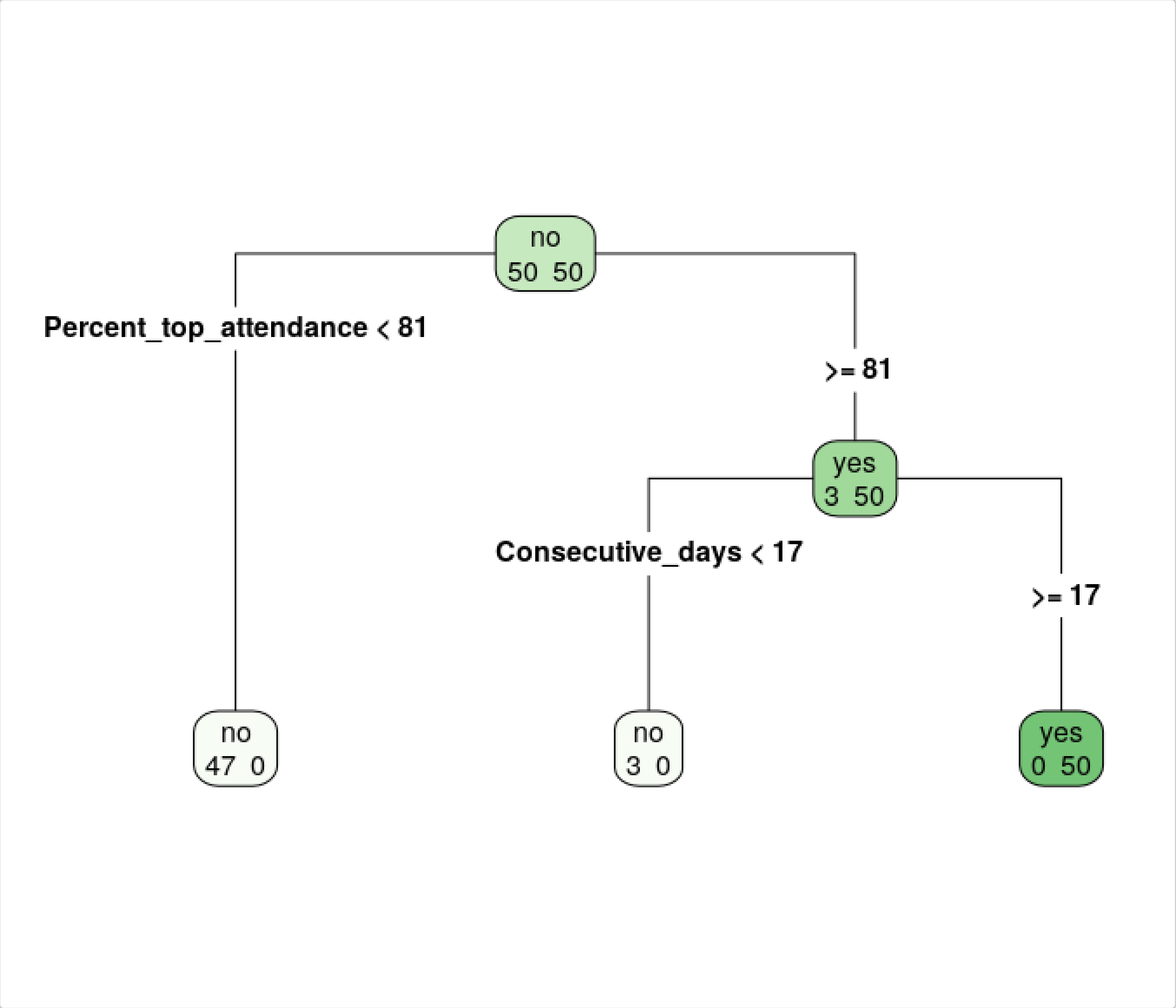


```
#> $`Type I error (false positives)`
#> [1] 0
#>
#> $`Type II error (false negatives)`
#> [1] 0.01
```

The resulting tree shows that two criteria can help us tell nests apart from non-nests: (1) a location needs to be visited for at least 81% of the time on the day the attendance is maximum and for at least 17 consecutive days to be a nest. These criteria translate in the following set of parameters:

- `min_top_att >= 81`, `min_consec >= 17`

#### Step 3: Identifying nests among revisited locations

After having identified one or more sets of parameter values that discriminate between nests and non-nests, we can make an informed decision on what values to use as input in the filtering arguments of `find_nests()`. Let's go ahead and run the function on the original wood stork dataset with the set of filtering parameter values suggested by the CART:

```
ws_output_2 <- find_nests(gps_data = woodstorks,
                          sea_start = "11-01",
                          sea_end = "08-31",
                          nest_cycle = 110,
                          buffer = 40,
                          min_pts = 2,
                          min_d_fix = 5,
                          min_consec = 17,
                          min_top_att = 81,
                          min_days_att = 1,
                          discard_overlapping = TRUE)
```

Note that for all other arguments in the function (those related to the species biology and to data characteristics) we are keeping the same values as we did the first time. However, this time we are setting `discard_overlapping = TRUE`. We no longer need to retain overlapping nesting attempts in the output.

```
ws_output_2$nests
```

```
#>           burst loc_id      long      lat first_date  last_date
#> 1 1134370-2013   2170 -80.85100 25.46283 2013-02-24 2013-05-02
#> 2 1134370-2013   3270 -79.39017 33.14267 2013-05-12 2013-06-03
#> 3  721290-2010    594 -81.64500 30.40483 2009-11-26 2010-08-16
#>   attempt_start attempt_end tot_vis days_vis consec_days perc_days_vis
#> 1    2013-03-04  2013-05-02     589       61          37         89.71
#> 2    2013-05-12  2013-06-03     219       22          20         95.65
#> 3    2010-04-26  2010-08-13    1066      143          37         54.17
#>   perc_top_vis
#> 1          100
#> 2          100
#> 3          100
```

Good news! The function identified the known nest correctly for both burst 1134370-2013 and 721290-2010. However, it also identified a second location for burst 1134370-2013. This looks like a second, probably failed, nesting attempt. We know that after leaving Rookery Branch, this wood stork migrated north to the Atlantic coast where its summer range is, and we cannot exclude that it attempted to nest a second time there.

The data on nesting attempts obtained as output of `find_nests()` will be the starting point for the second part of the workflow, where we will estimate reproductive outcome.

```
ws_nests <- ws_output_2
```

### Part II: Estimating reproductive outcome

#### Background

Nesting birds keep returning to their nest to feed their offspring until fledging. Histories of nest revisitation obtained from GPS data can be used to estimate the outcome of nesting attempts. Specifically, we can estimate if an attempt was successful or not according to whether it lasted as long as the duration of a complete nesting cycle for the focal species. The underlying assumption is that the nest stops being revisited after an attempt fails, which is true for many bird species.

Two elements complicate the picture: first, GPS fixes are not recorded in real time, which means some visits to a nest may be missed. Second, nest attendance can vary throughout the nesting cycle as a function of developmental stage of the chicks, so that visits may become less frequent and/or shorter with time. This means that the probability of observing a nest visit in the GPS data, given that it happened, may decrease with time during the nesting attempt.

Functions in `nestR` allow the estimation of reproductive outcome from GPS-derived nest revisitation histories while taking into account the probability of visit detection and allowing both detection and nest survival probability to vary in time.

#### Formatting nest revisitation histories

The function `format_attempts()` organizes the data obtained as output from `find_nests()` into the correct format for input into the next function. The output of `format_attempts()` is a list of two matrices. Both matrices have the same size, with as many rows as the number of nesting attempts identified by `find_nests()` and as many columns as the number of days in a complete nesting cycle. The first one is a matrix of fixes: for each attempt, it stores the number of GPS fixes that were taken on each day. The second one is a matrix of visits: for each attempt, it stores the number of nest visits that were recorded on each day.

```
ws_attempts <- format_attempts(nest_info = ws_nests, nest_cycle = 110)
```

```
# Matrix of fixes
class(ws_attempts$fixes)
#> [1] "matrix"
dim(ws_attempts$fixes)
#> [1]   3 110
```

```
# Matrix of visits
class(ws_attempts$visits)
#> [1] "matrix"
dim(ws_attempts$visits)
#> [1]   3 110
```

These two matrices will be used as input in the function that estimates the outcome of nesting attempts.

#### Nest survival model specification

The function `estimate_outcomes()` fits a Bayesian hierarchical model to the histories of nest revisitation and estimates the probability that each nest is still active (i.e., “alive”) on the last day of the attempt. Under the hood, `estimate_outcomes()` uses JAGS (Just Another Gibbs Sampler) to run the MCMC (Markov Chain Monte Carlo), via the package `rjags`. This gives the user the option of accessing the MCMC diagnostics from the `coda` package. The user can control parameters of the MCMC by passing them to the argument `mcmc_params`.

The model specification includes two processes: the survival process, which is not directly observable, and the observation process, which is the signal we get in the revisitation data. Much the same as in a Bayesian implementation of a Cormack-Jolly-Seber capture-mark-recapture model, the latent nest survival variable ($z_t$) is modelled at the daily scale as a function of survival status at the previous time-step ($z_{t-1}$) and daily survival probability ($\phi$):

$$z_t \sim Bern(z_{t-1} \times \phi_{t-1})$$

Observed visits on a given day ($Y_t$) are modelled as a function of current nest survival status ($z_t$), probability of visit detection on that day ($p_t$), and number of GPS fixes available on that day ($N_t$):

$$Y_t \sim Binom(z_t \times p_t, N_t)$$

Where the probability of detection is:

$$p_t = Pr(visit\ detected | z_t = 1, N_t)$$

Reproductive outcome is defined as the probability that value of $z_{nest\_cycle} = 1$, i.e., the probability that the nest was still surviving after `nest_cycle` days.

The user can choose among four models:

- `null` a null model, where both survival and detection probability are constant,
- `phi_time` a model where survival varies through time,
- `p_time` a model where detection probability varies through time, and
- `phi_time_p_time` a model where both survival and detection probability are a function of time.

In all four models, the parameters $\phi$ and $p$ are modeled using a binomial GLM as a function of the day of the attempt, $t$, i.e.:

$$logit(\phi_t) = \beta_{\phi 0} + \beta_{\phi 1} \times t$$

$$logit(p_t) = \beta_{p0} + \beta_{p1} \times t$$

In the case of the time-invariant models, $\beta_1$ is simply set to 0. The MCMC algorithm monitors the $\beta$ parameters (on the logit scale) for both $\phi$ and $p$, as well as the derived values of $\phi$ and $p$ (back-transformed). As we will see below, plotting and summary functions are available that take advantage of this information.

The model is fully specified by putting uninformative priors on the $\beta$ parameters, in this case a normal distribution with a mean of 0 and precision (i.e., $1/\sigma^2$) of 1e-5. I.e.:

$$\beta \sim N(\mu = 0, \tau = 1.0 \times 10^{-5})$$

Notice that we assume that daily survival ($\phi$) and detection ($p$) are the same for all nests in the population.

#### Model fitting and results

In the case of wood storks, we expect the probability of detection to decrease with time. Wood stork attendance at the nest is maximum during the initial phase of nesting, when birds are building their nest and forming pairs, and stays very high during incubation. We expect to observe nest visits on most of the days during this phase. After hatching, the male and female start alternating in leaving the nest to go on foraging trips. Still, the nest is never left unattended and at least one parent is at the nest for periods of multiple hours a day. The probability of detection should slightly decrease at this point but still remain high enough that visits are observed on the majority of days. When chicks grow big enough that they can thermoregulate on their own and increase their energetic demands, foraging trips become more and more frequent and the nest is left unattended for long chunks of time. Therefore, in the last few weeks before fledging the probability of detecting a nest visit is expected to drop. Therefore, the model with p varying through time is appropriate in the case of wood storks. We run `estimate_outcomes()` on `ws_attempt` using the model where $p$ varies with time:

```
# Using default MCMC parameters (mcmc_params)

ws_outcomes <- estimate_outcomes(fixes = ws_attempts$fixes,
                                 visits = ws_attempts$visits,
                                 model = "p_time")

str(ws_outcomes)
```

The output of `estimate_outcomes()` is a list including the following objects of class `mcarray`:

- `p` stores the population-level value of $p$ on each day of the attempt.
- `p.b0` is the value of the intercept of $p$.
- `p.b1` is the value of the slope of $p$.
- `phi` stores the population-level value of $\phi$ on each day of the attempt.
- `phi.b0` is the value of the intercept of $\phi$.
- `phi.b1` is the value of the slope of $\phi$.
- `z` stores the daily survival for each attempt.
- `names` stores the attempt ID, in the form `burst_loc_id`.
- `model` is the name of the chosen model structure.

We can now plot survival and detection probability through time at the population level, as well as daily survival estimated individually for each nest. Survival is constant through time as specified by the chosen model. As expected, detection probability decreases with time.

```
# Plot population-level daily survival
plot_survival(ws_outcomes)
```

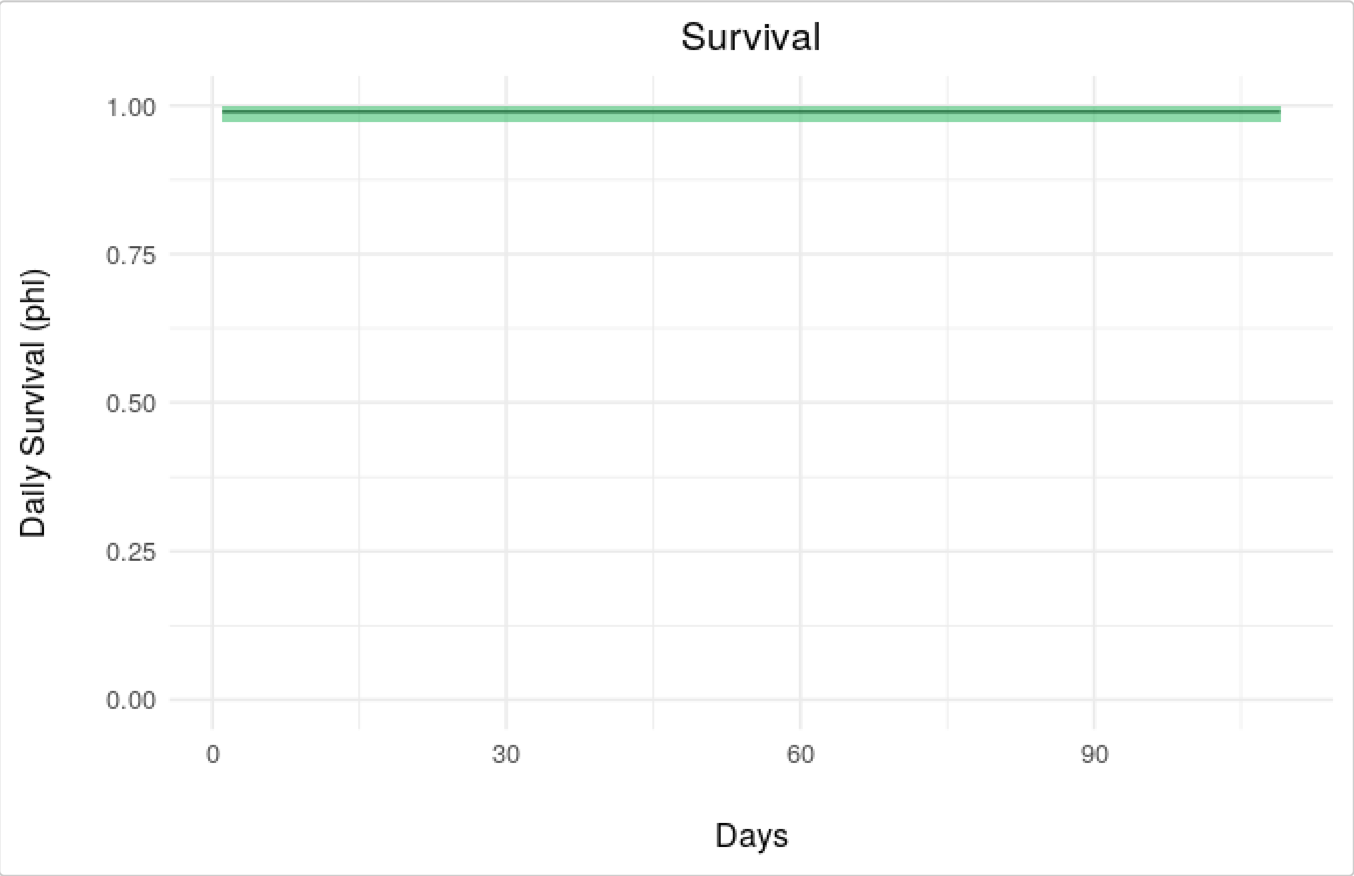


```r
# Plot population-level detection
plot_detection(ws_outcomes)
```

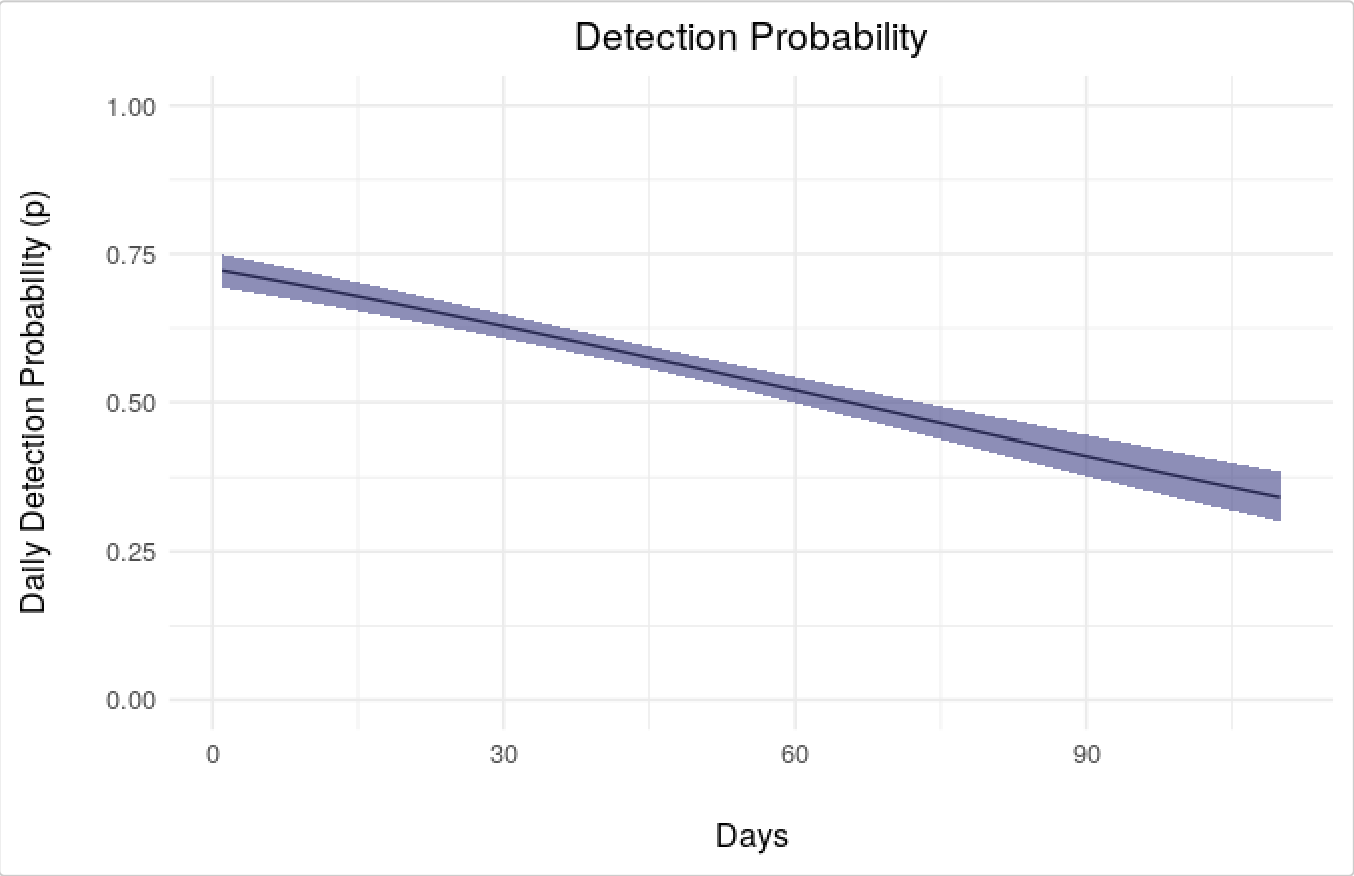

The MCMC algorithm also monitors the $z_t$ for all bursts, so we are able to specifically visualize the survival status for each nest.

```
# Plot estimated survival for burst 1
plot_nest_surv(ws_outcomes, who = 1)
```

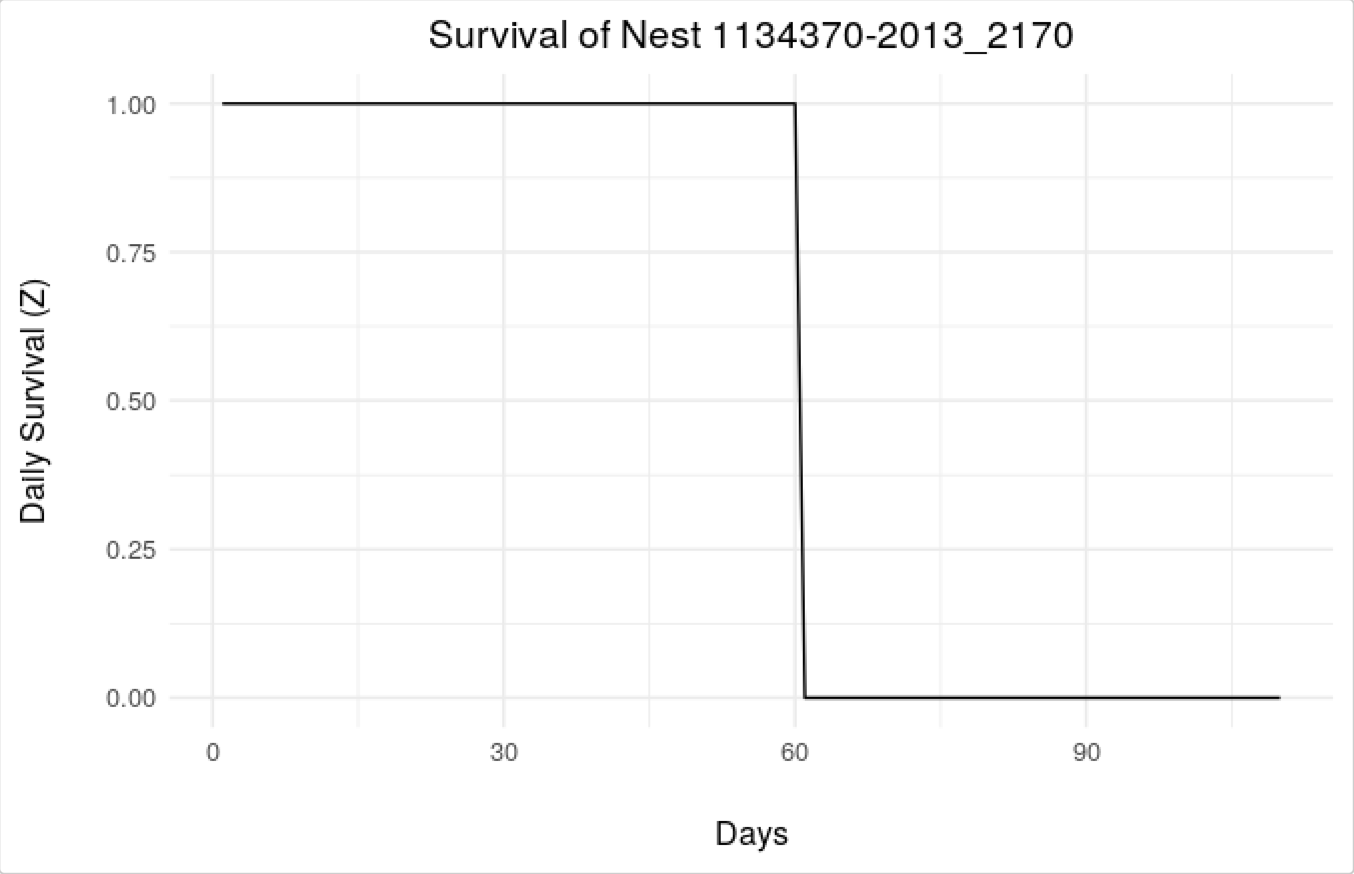


```
# Plot estimated survival for burst 2
plot_nest_surv(ws_outcomes, who = 2)
```

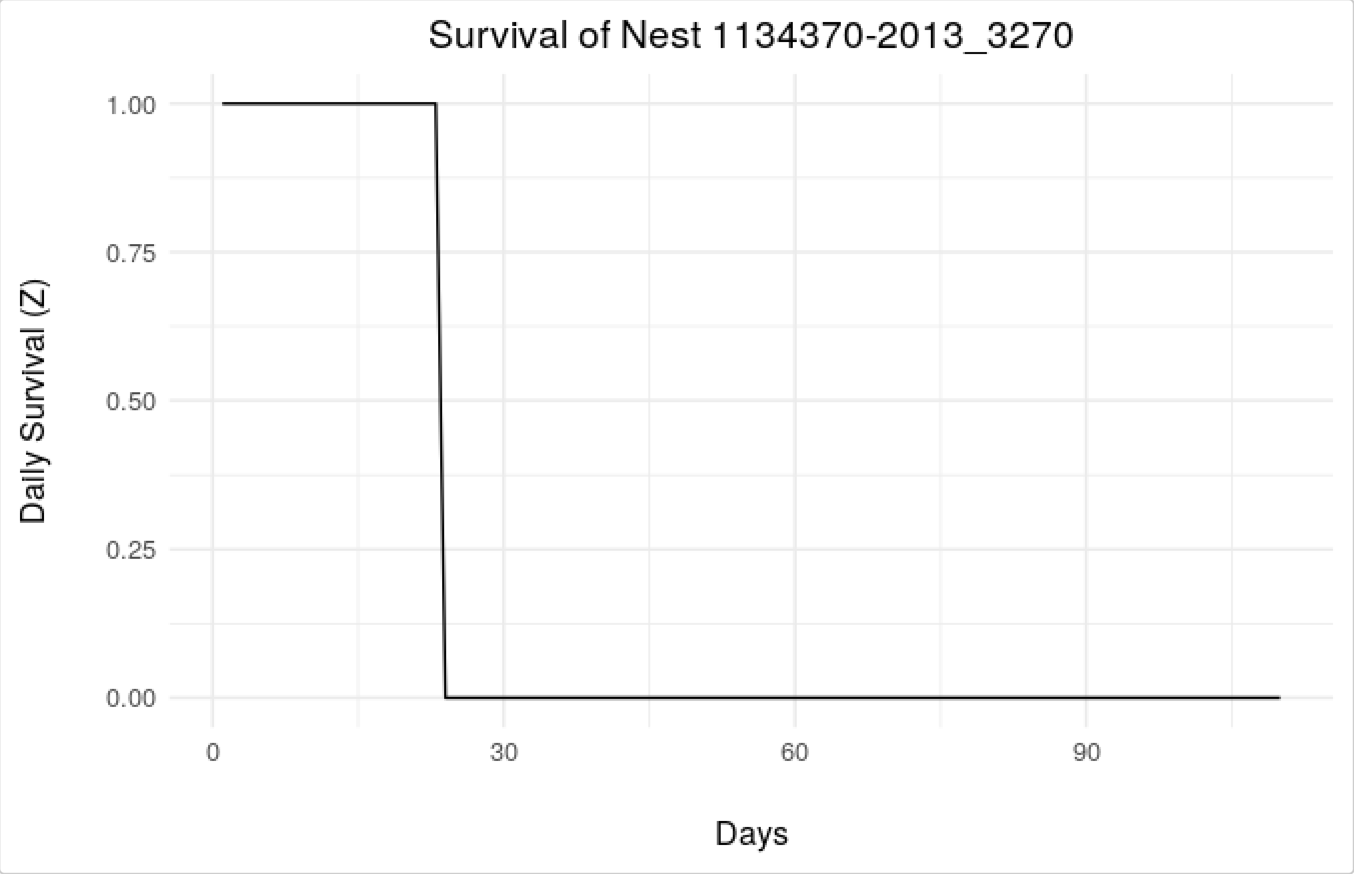


```
# Plot estimated survival for burst 3
plot_nest_surv(ws_outcomes, who = 3)
```

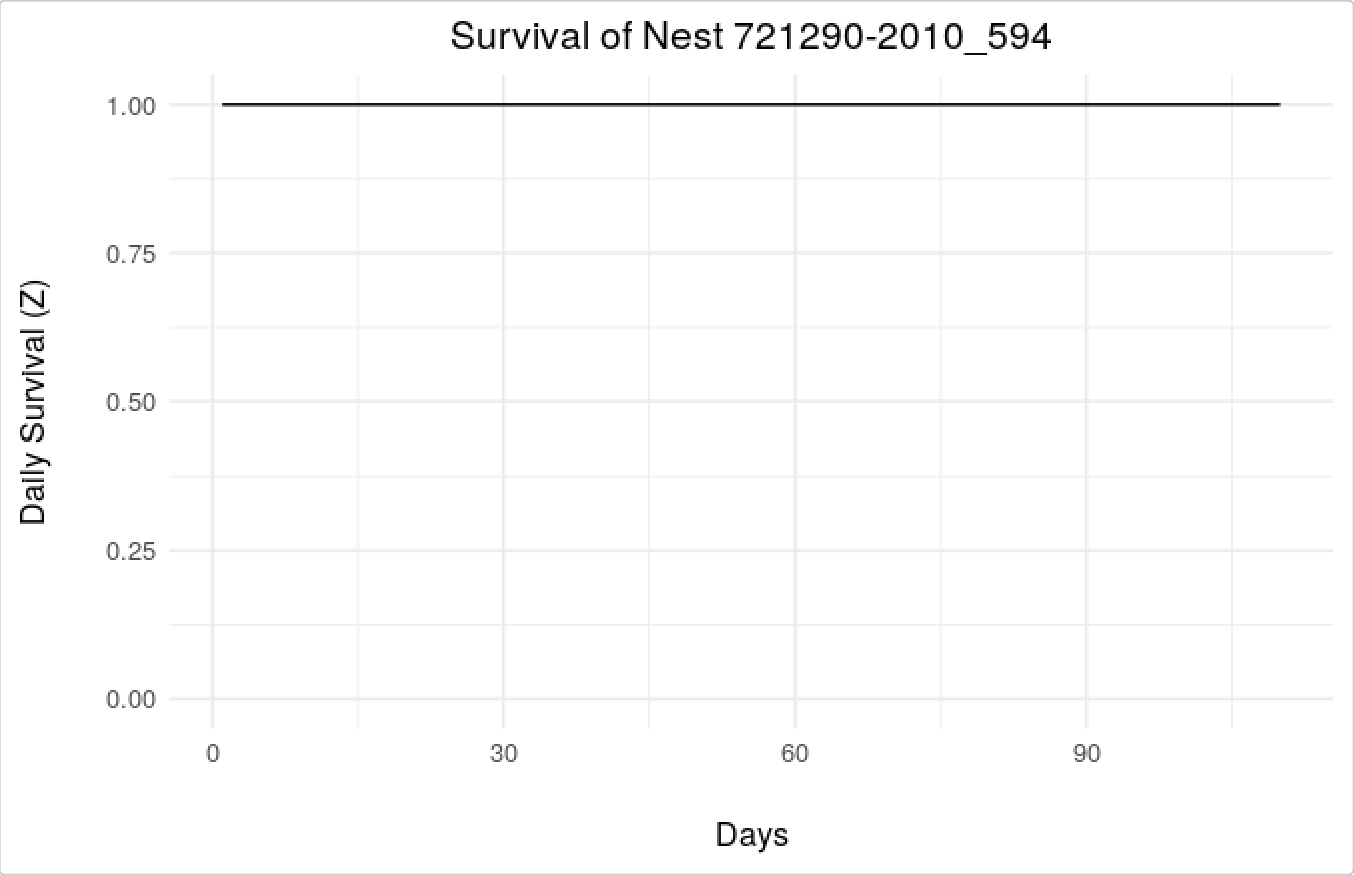

The last three plots inform us of the fate of the nesting attempts: two out of three appeared to have failed on day 60 and day 23, respectively, while the third was successful.

Finally, the function `summarize_outcomes()` formats the output of the MCMC into a `list` including:

- `phi`, a `data.frame` storing the mean value of $phi$ and the upper and lower credible interval limits.
- `p`, a `data.frame` storing the mean value of $p$ and the upper and lower credible interval limits.
- `outcomes`, a `data.frame` listing each nesting attempt, the probability that it was successful, and the day until which the nest survived.

```
summarize_outcomes(ws_outcomes, ci = 0.95)
#> $phi
#>         lwr      mean       upr
#> 1 0.9712795 0.9897539 0.9987782
#>
#> $p
#>      b0_lwr   b0_mean   b0_upr      b1_lwr     b1_mean      b1_upr
#> 1 0.8259129 0.9687598 1.107882 -0.01738344 -0.01478975 -0.01217242
#>
#> $outcomes
#>               burst pr_succ_lwr pr_succ_mean pr_succ_upr last_day_lwr
#> 1 1134370-2013_2170           0            0           0           60
#> 2 1134370-2013_3270           0            0           0           23
#> 3   721290-2010_594           1            1           1          110
#>   last_day_mean last_day_upr
#> 1            60           60
#> 2            23           23
#> 3           110          110
```

The `coda` diagnostics are available after converting the output of `estimate_outcomes()` into `mcmc.list`. For example, if you wanted to look at the diagnostic plots for the $\beta$ parameters on detection ($p$):

```
ws_pb0_coda <- as.mcmc.list(ws_outcomes$p.b0)
ws_pb1_coda <- as.mcmc.list(ws_outcomes$p.b1)

plot(ws_pb0_coda)
```

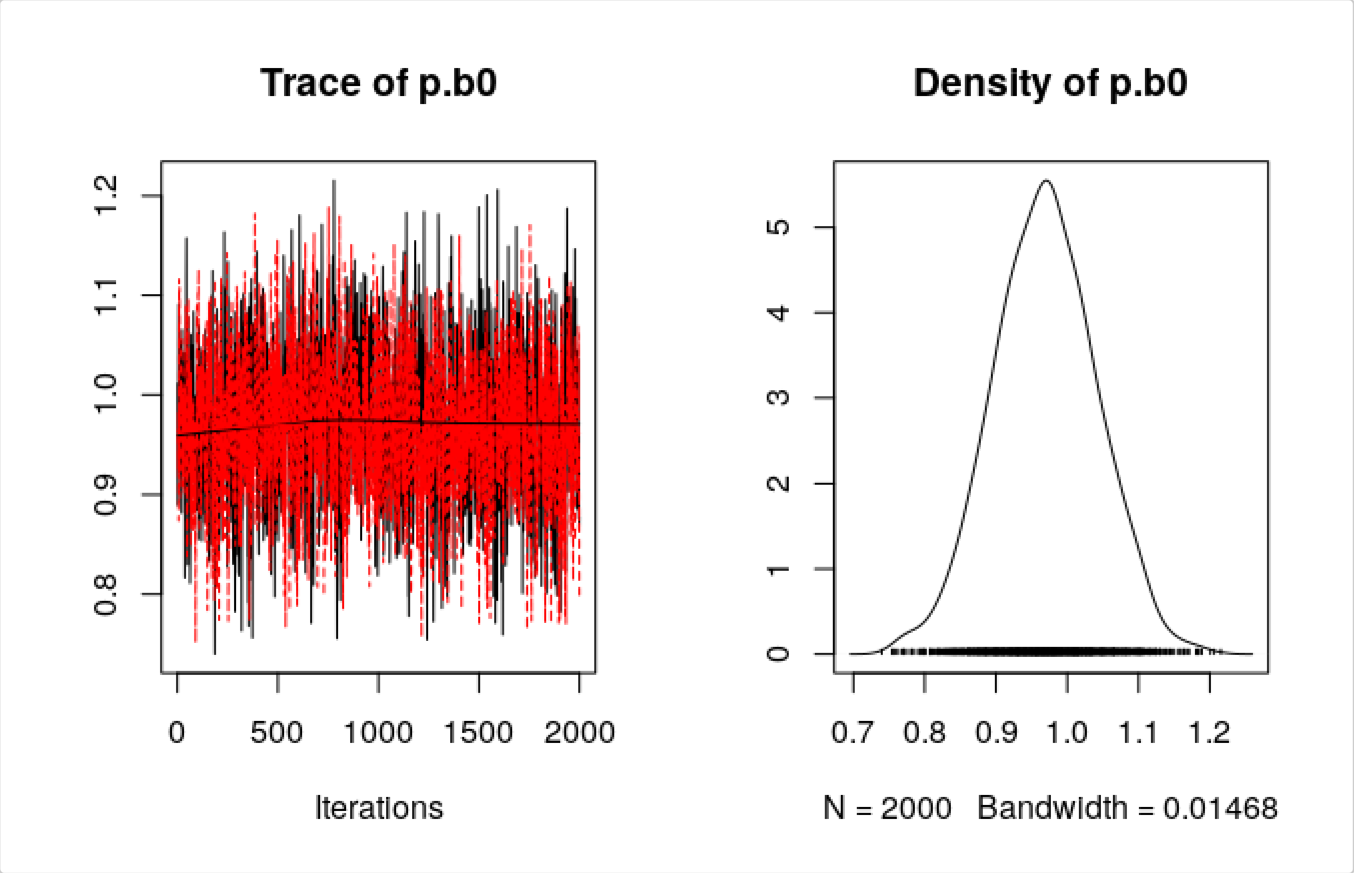


```
plot(ws_pb1_coda)
```

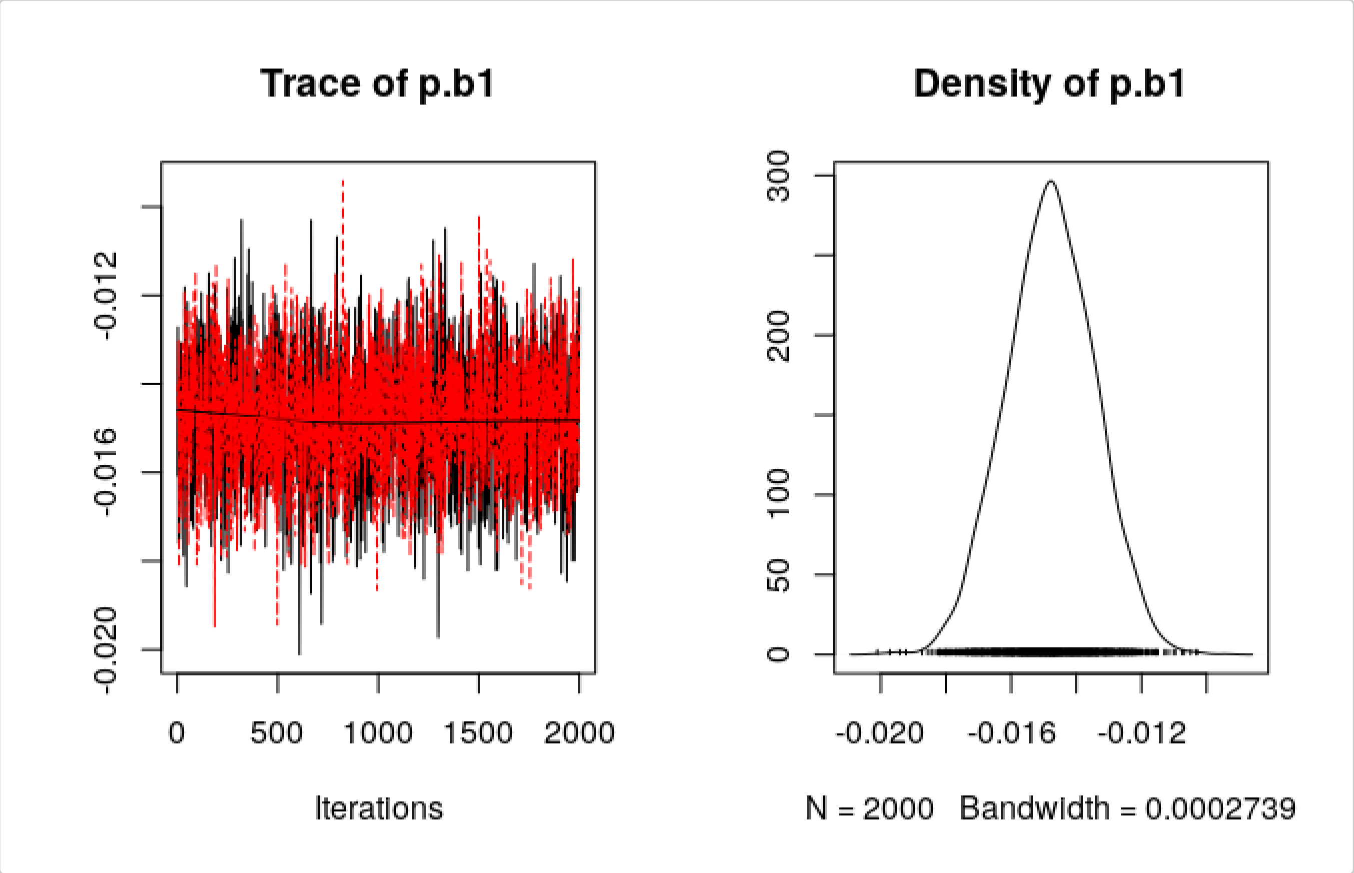

### Case studies: applying the method to different species and data

#### Lesser kestrels

Lesser kestrels nest between mid-April and late-July in southern Italy. Their nesting cycle is approximately 65 days long from egg-laying to fledging. However, the kestrels in our dataset were captured on the nest and tagged during the early chick-rearing phase. This means that we are missing the initial part of nesting attempts (from egg-laying to shortly after hatching, ~30 days). To account for this in our analysis, we subtract 30 days from the actual duration of a full-length nesting attempt, and set the value of `nest_cycle` to the longest attempt duration that we expect to be able to observe in our data. To allow some room for variability in the onset of nesting, we constrain the nesting season between March 31st and August 31st. This temporal range will definitely include any nesting attempts in our study population. The GPS tags were programmed to collect a fix every 15 minutes, so we should bump up the value of `min_d_fix` to a higher value than we did for wood storks. We run a first screening with loose filtering constraints:

```
lk_output_1 <- find_nests(gps_data = kestrels,
                          sea_start = "03-31",
                          sea_end = "08-31",
                          nest_cycle = 35,
                          buffer = 40,
                          min_pts = 2,
                          min_d_fix = 15,
                          min_consec = 2,
                          min_top_att = 1,
                          min_days_att = 1,
                          discard_overlapping = FALSE)
```

Here are the results for the first kestrel:

```
lk_output_1$nests %>% filter(burst == "16680-2017") %>% head()
```

```
#>        burst loc_id     long      lat first_date  last_date attempt_start
#> 1 16680-2017   1282 16.55029 40.83296 2017-06-15 2017-07-31    2017-06-26
#> 2 16680-2017    207 16.55209 40.82840 2017-06-15 2017-07-30    2017-06-15
#> 3 16680-2017   1761 16.54268 40.89704 2017-07-05 2017-07-15    2017-07-08
#> 4 16680-2017    629 16.55166 40.82864 2017-06-17 2017-07-30    2017-06-17
#> 5 16680-2017   1413 16.55018 40.83261 2017-06-17 2017-07-29    2017-07-10
#> 6 16680-2017    617 16.53573 40.89567 2017-06-20 2017-06-27    2017-06-20
#>   attempt_end tot_vis days_vis consec_days perc_days_vis perc_top_vis
#> 1  2017-07-30     236       47          47        100.00        37.04
#> 2  2017-07-19     207       44          27         95.65        79.17
#> 3  2017-07-15      18        6           2         54.55        31.58
#> 4  2017-07-21      16       13           3         29.55        10.00
#> 5  2017-07-29      16       12           2         27.91        11.11
#> 6  2017-06-27      14        7           7         87.50        17.39
```

Here are the results for the second kestrel:

```r
lk_output_1$nests %>% filter(burst == "16682-2017") %>% head()
#>        burst loc_id     long      lat first_date  last_date attempt_start
#> 1 16682-2017   1514 16.38192 41.00030 2017-07-05 2017-07-25    2017-07-05
#> 2 16682-2017     23 16.41666 40.81702 2017-06-16 2017-07-04    2017-06-16
#> 3 16682-2017   1473 16.40184 41.03227 2017-07-10 2017-07-14    2017-07-10
#> 4 16682-2017    144 16.42090 40.81645 2017-06-16 2017-07-31    2017-06-27
#> 5 16682-2017    571 16.42425 40.81612 2017-06-25 2017-07-30    2017-06-27
#> 6 16682-2017   2349 16.38337 41.01260 2017-07-22 2017-07-25    2017-07-24
#>   attempt_end tot_vis days_vis consec_days perc_days_vis perc_top_vis
#> 1  2017-07-25     127       21          21        100.00        30.00
#> 2  2017-07-04      44       13           6         68.42        75.00
#> 3  2017-07-14      43        4           3         80.00        30.36
#> 4  2017-07-31      41       12           5         26.09        21.88
#> 5  2017-07-30      36       11           4         30.56        18.18
#> 6  2017-07-25      35        3           2         75.00        37.04
```

Having tagged the birds after hatching means we do not have GPS data for the first part of the nesting attempt, which can potentially make the detection of nests more difficult because nest attendance usually decreases after hatching and visits become less frequent. However, on the bright side, it also means that we have information about the exact location of the nests. We can use this information to get some exploratory data on nests vs. non-nests:

```r
# Load data on known nest locations
data(lk_known_nests)

(lk_explodata <- get_explodata(candidate_nests = lk_output_1$nests,
                               known_coords = lk_known_nests,
                               buffer = 40))
#>        burst loc_id     long      lat first_date  last_date attempt_start
#> 1 16680-2017    207 16.55209 40.82840 2017-06-15 2017-07-30    2017-06-15
#> 2 16682-2017     23 16.41666 40.81702 2017-06-16 2017-07-04    2017-06-16
#> 3 16680-2017   1282 16.55029 40.83296 2017-06-15 2017-07-31    2017-06-26
#> 4 16682-2017    144 16.42090 40.81645 2017-06-16 2017-07-31    2017-06-27
#>   attempt_end tot_vis days_vis consec_days perc_days_vis perc_top_vis nest
#> 1  2017-07-19     207       44          27         95.65        79.17  yes
#> 2  2017-07-04      44       13           6         68.42        75.00  yes
#> 3  2017-07-30     236       47          47        100.00        37.04   no
#> 4  2017-07-31      41       12           5         26.09        21.88   no
```

In this simplified version of the workflow, with only two individuals, it is easy to just visually pick the best values to discriminate nests from non-nests. In general, manual tuning of filtering parameter values is viable and encouraged whenever there is good knowledge of the species' ecology and a clear behavioral signal is visible in the data. This can become challenging for very large datasets that are not easy to explore visually or in which there is much variability. In those situations, the user can use statistical tools to inform the choice of parameter values and has to be willing to accept some degree of error. On the other hand, using large amounts of data to inform the choice of parameter values will likely help with generalization, and it will improve our ability to find nests in trajectories for which no prior information is available (species and data characteristics being equal).

```r
lk_output_2 <- find_nests(gps_data = kestrels,
                          sea_start = "03-31",
                          sea_end = "08-31",
                          nest_cycle = 35,
                          buffer = 40,
                          min_pts = 2,
                          min_d_fix = 15,
                          min_consec = 6,
                          min_top_att = 70,
                          min_days_att = 65,
                          discard_overlapping = TRUE)

lk_output_2$nests
```

```
#>        burst loc_id     long      lat first_date  last_date attempt_start
#> 1 16680-2017    207 16.55209 40.82840 2017-06-15 2017-07-30    2017-06-15
#> 2 16682-2017     23 16.41666 40.81702 2017-06-16 2017-07-04    2017-06-16
#>   attempt_end tot_vis days_vis consec_days perc_days_vis perc_top_vis
#> 1  2017-07-19     207       44          27         95.65        79.17
#> 2  2017-07-04      44       13           6         68.42        75.00
```

The output accurately reflects our prior data on nest locations for the two kestrels in the dataset.

```r
lk_nests <- lk_output_2
```

The kestrel colonies were periodically revisited after tagging to keep track of the progress of nesting attempts. This means that we can compare the output of the model with the data we have and verfiy that the estimation of reproductive outcome was correct.

We can proceed with the estimation of reproductive outcome. Again, we use 35 days as `nest_cycle` instead of the actual 65 days to account for the fact that the first part of the attempts was missed.

```r
lk_attempts <- format_attempts(nest_info = lk_nests, nest_cycle = 35)
```

We can now run the nest survival model. We do not expect visit detection probability to change with time, because the frequency of visits should not vary much after the early chick-rearing phase.

```r
lk_outcomes <- estimate_outcomes(fixes = lk_attempts$fixes,
                                 visits = lk_attempts$visits,
                                 model = "null")
```

Here are the summary plots for survival and detection probability at the population level, as well as the outcome of the two nesting attempts in the dataset:

```r
gridExtra::grid.arrange(plot_survival(lk_outcomes),
```

```
                       plot_detection(lk_outcomes),
                       ncol = 2)
```

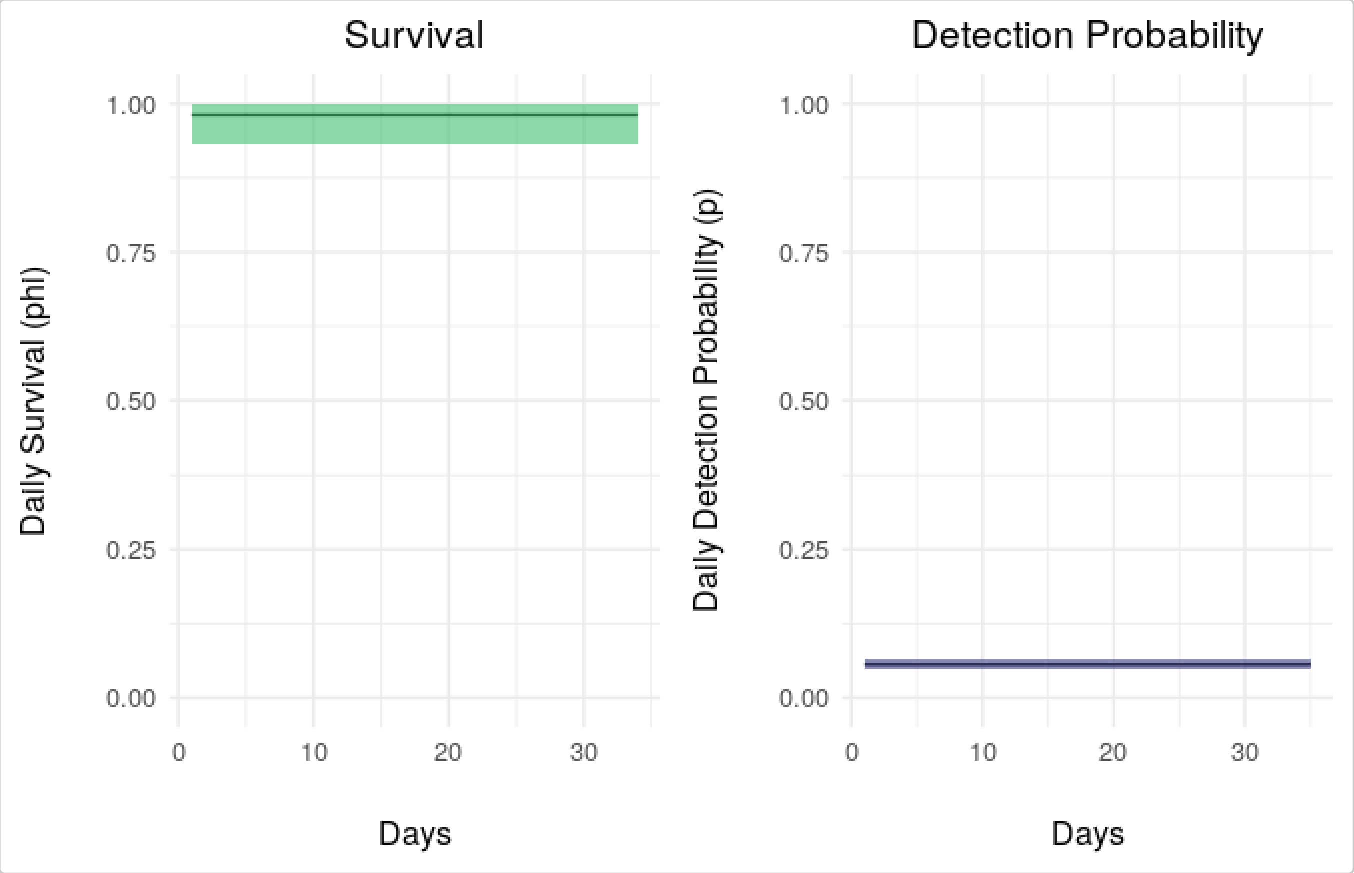


```
gridExtra::grid.arrange(plot_nest_surv(lk_outcomes, who = 1),
                       plot_nest_surv(lk_outcomes, who = 2),
                       ncol = 2)
```

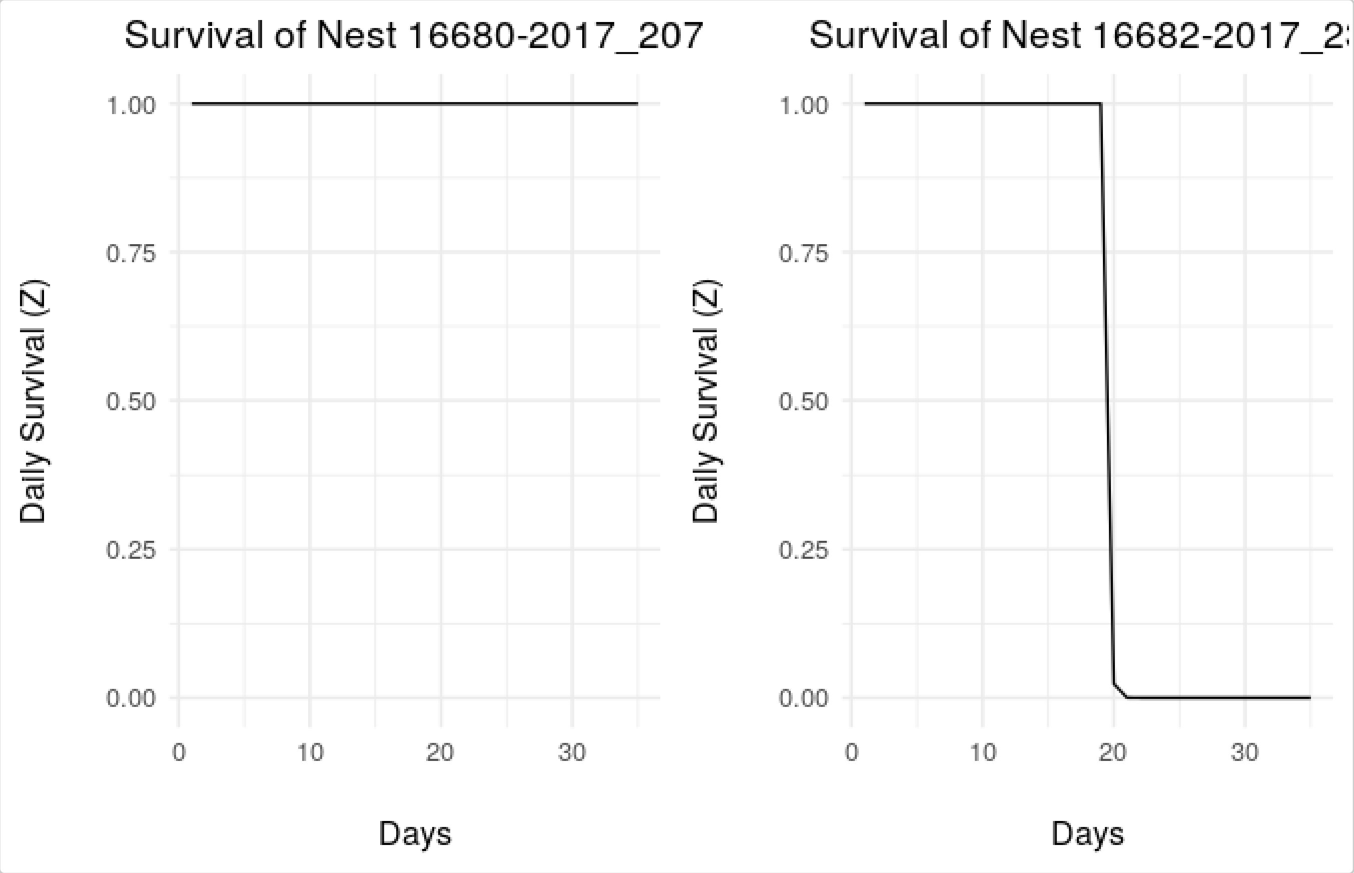


According to the model, kestrel 16680-2017 was successful, while kestrel 16682-2017 failed at around 19 days (plus the 30 days we subtracted, therefore at approximately 50 days into the actual attempt).

```
summarize_outcomes(lk_outcomes, ci = 0.95)
#> $phi
#>         lwr      mean       upr
#> 1 0.9301438 0.9809687 0.9994558
#>
#> $p
#>          lwr       mean        upr
#> 1 0.04869226 0.05630568 0.06435249
#>
#> $outcomes
#>            burst pr_succ_lwr pr_succ_mean pr_succ_upr last_day_lwr
#> 1 16680-2017_207           1            1           1           35
#> 2  16682-2017_23           0            0           0           19
#>   last_day_mean last_day_upr
#> 1      35.00000           35
#> 2      19.02375           19
```

#### Mediterranean gulls

Mediterranean gulls nest between mid-April and late-July in Italy. Similarly to lesser kestrels, their nesting cycle is approximately 60 days long from egg-laying to fledging. However, the gulls in our dataset were captured on the nest and tagged during incubation. This means that we are missing the initial part of nesting attempts (from egg-laying to early incubation, approximately 20 days). Like we did for kestrels, we subtract the duration of the missing part from the actual duration of a full-length nesting attempt, and set the value of `nest_cycle` to 40 days. Probably because the 15-minute time resolution and the low fix failure rate make the individual datasets very large, we easily incurred in memory issues when running `find_nests()` on this dataset. To counteract this issue, we reduce the number of points used to compute the distance matrix by using strict temporal limits for the nesting season (April 15th to August 1st). Next time we will buy a computer with more RAM. We run the first screening with loose filtering constraints:

```
data(gulls)

mg_output_1 <- find_nests(gps_data = gulls,
                          sea_start = "04-15",
                          sea_end = "08-01",
                          nest_cycle = 40,
                          buffer = 40,
                          min_pts = 2,
                          min_d_fix = 15,
                          min_consec = 2,
                          min_top_att = 1,
                          min_days_att = 1,
                          discard_overlapping = FALSE)
```

Here is the output for the first gull:

```
mg_output_1$nests %>% filter(burst == "URI05-2016") %>% head()
```

```
#>         burst loc_id     long      lat first_date  last_date attempt_start
#> 1 URI05-2016     79 12.32500 44.23812 2016-05-18 2016-05-28    2016-05-18
#> 2 URI05-2016   1285 15.89975 41.49545 2016-06-08 2016-06-27    2016-06-08
#> 3 URI05-2016   1311 16.01907 41.42352 2016-06-06 2016-06-27    2016-06-14
#> 4 URI05-2016   1863 16.07903 41.39988 2016-06-06 2016-07-01    2016-06-21
#> 5 URI05-2016   1752 16.07228 41.37127 2016-06-21 2016-06-29    2016-06-21
#> 6 URI05-2016   1013 15.90350 41.49407 2016-06-05 2016-06-26    2016-06-05
#>   attempt_end tot_vis days_vis consec_days perc_days_vis perc_top_vis
#> 1  2016-05-28     233       11          11        100.00        86.84
#> 2  2016-06-27     193       12          10         60.00        85.00
#> 3  2016-06-27     110       11           4         50.00        87.88
#> 4  2016-07-01     104       12           5         46.15        71.79
#> 5  2016-06-29      59        7           4         77.78        66.67
#> 6  2016-06-26      34        7           2         31.82        41.94
```

And here is the output for the second gull:

```
mg_output_1$nests %>% filter(burst == "URI29-2016") %>% head()
#>         burst loc_id     long      lat first_date  last_date attempt_start
#> 1 URI29-2016    359 12.32513 44.23807 2016-05-24 2016-07-12    2016-05-26
#> 2 URI29-2016   2070 12.32515 44.23770 2016-05-26 2016-07-09    2016-06-04
#> 3 URI29-2016   2230 12.39975 44.20573 2016-06-27 2016-07-08    2016-06-27
#> 4 URI29-2016   3396 12.32748 44.25205 2016-07-04 2016-07-12    2016-07-04
```

```
#> 5 URI29-2016    116 12.32477 44.23832 2016-05-24 2016-07-01    2016-05-24
#> 6 URI29-2016   3264 12.34268 44.21670 2016-06-14 2016-07-11    2016-06-16
#>   attempt_end tot_vis days_vis consec_days perc_days_vis perc_top_vis
#> 1  2016-07-04    1222       49          41         98.00        90.14
#> 2  2016-07-09      72       28          12         62.22        15.79
#> 3  2016-07-08      58        9           3         75.00        20.55
#> 4  2016-07-12      58        6           4         66.67        31.43
#> 5  2016-07-01      56       22           8         56.41        22.22
#> 6  2016-07-11      56       17           4         60.71        18.92
```

Since the gulls were also captured on the nests, we know their location. We can use that information to compare parameters between nests and non-nests:

```
data(mg_known_nests)

(mg_explodata <- get_explodata(candidate_nests = mg_output_1$nests,
                               known_coords = mg_known_nests,
                               buffer = 40))
#>        burst loc_id     long      lat first_date  last_date attempt_start
#> 1 URI05-2016     79 12.32500 44.23812 2016-05-18 2016-05-28    2016-05-18
#> 2 URI29-2016    359 12.32513 44.23807 2016-05-24 2016-07-12    2016-05-26
#> 3 URI05-2016    102 12.32447 44.23810 2016-05-18 2016-05-23    2016-05-20
#> 4 URI29-2016   2070 12.32515 44.23770 2016-05-26 2016-07-09    2016-06-04
#>   attempt_end tot_vis days_vis consec_days perc_days_vis perc_top_vis nest
#> 1  2016-05-28     233       11          11        100.00        86.84  yes
#> 2  2016-07-04    1222       49          41         98.00        90.14  yes
#> 3  2016-05-23       9        4           2         66.67        17.65   no
#> 4  2016-07-09      72       28          12         62.22        15.79   no
```

We tune filtering parameters according to what we found in the exploratory data:

```
mg_output_2 <- find_nests(gps_data = gulls,
                    sea_start = "04-15",
                    sea_end = "08-01",
                    nest_cycle = 40,
                    buffer = 40,
                    min_pts = 2,
                    min_d_fix = 15,
                    min_consec = 10,
                    min_top_att = 80,
                    min_days_att = 90,
                    discard_overlapping = TRUE)

mg_output_2$nests
```

```
#>        burst loc_id     long      lat first_date  last_date attempt_start
#> 1 URI05-2016     79 12.32500 44.23812 2016-05-18 2016-05-28    2016-05-18
#> 2 URI29-2016    359 12.32513 44.23807 2016-05-24 2016-07-12    2016-06-02
#>   attempt_end tot_vis days_vis consec_days perc_days_vis perc_top_vis
```

```
#> 1  2016-05-28     233       11          11            100        86.84
#> 2  2016-07-11    1222       49          41             98        90.14
```

The output includes both the correct nesting locations.

```
mg_nests <- mg_output_2
```

Let's proceed with the estimation of reproductive outcome. Like for kestrels, we have information on the outcome of these two nesting attempts for Mediterranean gulls, and we will be able to verify our results.

We set the size of the matrix to account for the missing initial part of the attempts, setting `nest_cycle` to 40 days.

```
mg_attempts <- format_attempts(nest_info = mg_nests, nest_cycle = 40)
```

Part of the incubation phase was still captured in the data, so we do expect the detection probability to vary through time in this case. We pick the model with time-varying $p$:

```
mg_outcomes <- estimate_outcomes(fixes = mg_attempts$fixes,
                                 visits = mg_attempts$visits,
                                 model = "p_time")
```

Let's take a look at the summary plots:

```
gridExtra::grid.arrange(plot_survival(mg_outcomes),
                        plot_detection(mg_outcomes),
                        ncol = 2)
```

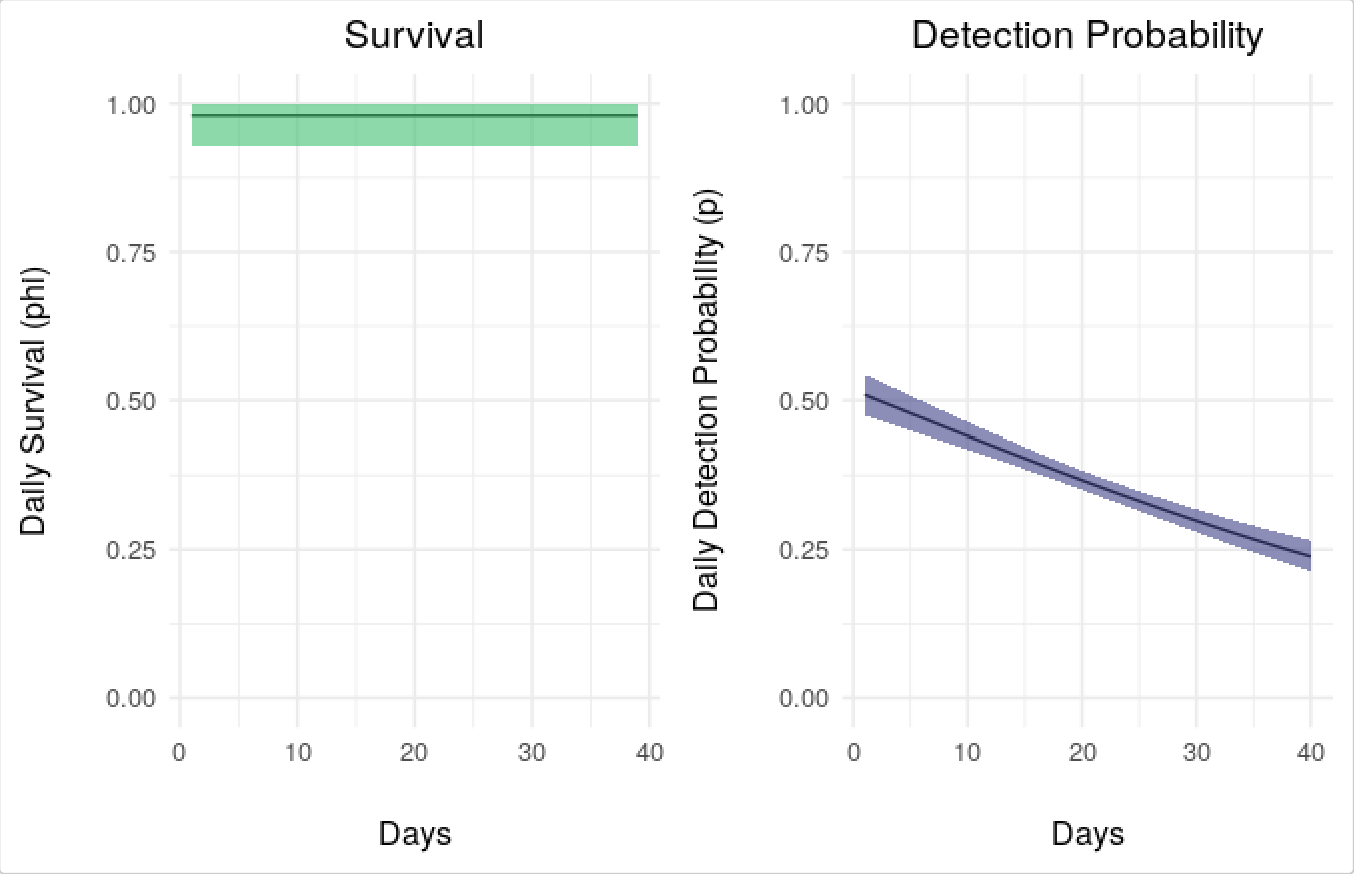


```
gridExtra::grid.arrange(plot_nest_surv(mg_outcomes, who = 1),
                        plot_nest_surv(mg_outcomes, who = 2),
                        ncol = 2)
```

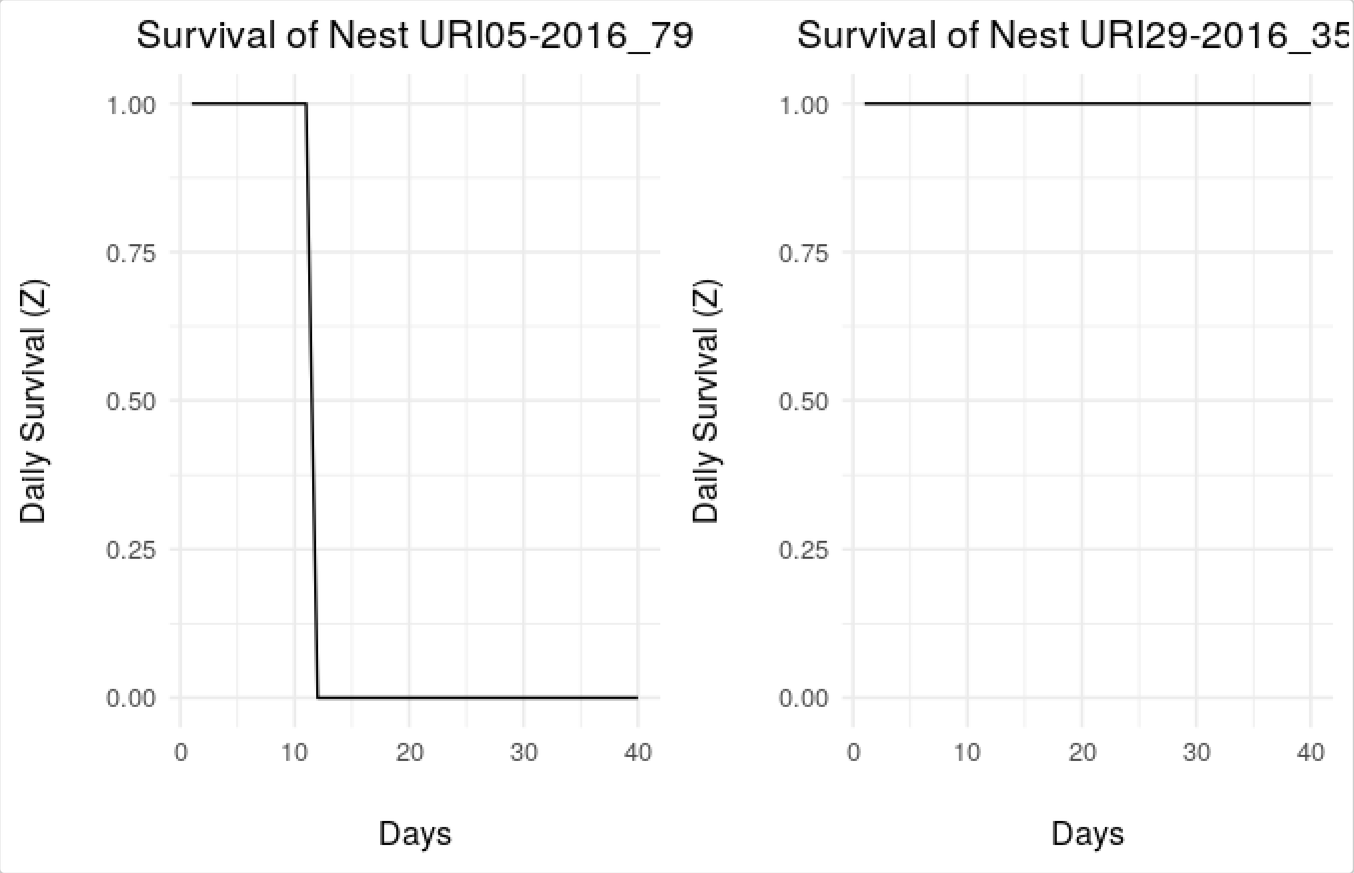


The estimation of reproductive outcome matches our prior information. Gull URI29-2016 successfully completed its nesting attempt, while gull URI05-2016 failed at 11 days (plus the 20 days that we are missing, therefore at about 30 days from the start of the attempt).

```
summarize_outcomes(mg_outcomes, ci = 0.95)
#> $phi
#>         lwr      mean       upr
#> 1 0.9268635 0.9798267 0.9995114
#>
#> $p
#>        b0_lwr    b0_mean    b0_upr      b1_lwr     b1_mean      b1_upr
#> 1 -0.07169135 0.06807269 0.2105537 -0.03665198 -0.03076287 -0.02442803
#>
#> $outcomes
#>            burst pr_succ_lwr pr_succ_mean pr_succ_upr last_day_lwr
#> 1  URI05-2016_79           0            0           0           11
#> 2 URI29-2016_359           1            1           1           40
#>   last_day_mean last_day_upr
#> 1            11           11
#> 2            40           40
```

#### Conclusions

In this vignette, we illustrated how to use functions in the package `nestR` to identify nesting locations along movement trajectories of birds and, based on the history of nest revisitation, to estimate the outcome of nesting attempts. The package provides tools to deal with both situations where prior information on nests is available and where it is not. Functions are designed with flexibility in mind, to accommodate the needs of users dealing with different types of data or focusing on different species. Being able to estimate individual reproductive outcome from tracking data provides an unprecedented opportunity to investigate links between reproductive fitness and factors affecting it in cases where on-ground data is not available.
