## Additional file 2 for "Analysis of movement recursions to detect reproductive events and estimate their fate in central place foragers"

### Package ‘nestR’

December 2, 2019

**Type** Package

**Title** Estimation of Bird Nesting from Tracking Data

**Version** 1.1.0

**Maintainers@R**

**Description** A package to locate nests and estimate avian reproductive outcome from GPS-tracking data. It includes functions to identify nest sites along bird movement trajectories based on patterns of location revisitation, dynamically explore results, estimate the outcome of identified nesting attempts, and plot and summarize outputs.

**License** MIT + file LICENSE

**Encoding** UTF-8

**LazyData** true

**Depends** dplyr,
lubridate,
data.table,
geosphere,
rjags,
shiny,
shinydashboard,
leaflet,
kableExtra,
gridExtra,
purrr

**Imports** forcats,
rpart,
rpart.plot

**RoxygenNote** 6.1.1

**Suggests** knitr,
rmarkdown

**VignetteBuilder** knitr

### R topics documented:

**Index** **24**

---

`attempt_limits` *Determine start and end dates of nesting attempt*

---

#### Description

`attempt_limits` Determines the start and end dates of the potential nesting attempt based on the patterns of revisitation of the candidate nest.

#### Usage

```
attempt_limits(x, min_consec, nest_cycle)
```

#### Arguments

| | |
|---|---|
| x | data.frame of daily revisitation history of a candidate nest |
| min_consec | Integer. Minimum number of consecutive days a location needs to be visited to be considered as a candidate nest. See Details |
| nest_cycle | Integer. Duration (in days) of a complete nesting cycle |

#### Details

Used with lapply inside function revisit_stats.

The function uses a moving window of size nest_cycle to find the most likely time range of the nesting attempt.

Attendance is expected to be maximum during the initial phase of nesting, so we assume that the longest series of consecutive days corresponds to the beginnig of the nesting attempt. Therefore, the moving window starts at the beginning of the first series of consecutive days longer than min_consec. This helps discard any early visits to the nest before the actual start of the attempt.

After that, the function slides a moving window of size nest_cycle to the data until it hits the last visit. The window that encompasses the highest number of visits is selected as most likely delimiting the nesting attempt.

If the last nest visit is on the end date of the window or later, the end of the attempt is set at the end date of the window (which is the start date + nest_cycle). This helps discard any later visits to the nest after the attempt is already concluded.

If the last nest visit is earlier than the ending date of the window, the end of the attempt is set at the date of the last visit.

#### Value

Returns data.frame with start and end dates of the attempt and number of nest visits within it.

---

candidate_summary *Summarize number of points within candidate buffers*

---

#### Description

candidate_summary counts the number of points grouped within each candidate buffer and arranges them in descending order.

#### Usage

```
candidate_summary(cands)
```

#### Arguments

| | |
|---|---|
| cands | data.frame of associations between points and nest candidates returned by get_candidates |

##### Value

Returns tibble counting number of points within each candidate buffer.

---

check_input *Check input data format*

---

##### Description

check_input checks that the data input in find_nests is in the correct format.

##### Usage

```
check_input(gps_data)
```

##### Arguments

| | |
|---|---|
| gps_data | data.frame of GPS data |

##### Details

The function checks that the input data includes burst, date-time, and lat/long coordinates.

---

choose_overlapping *Handle overlapping attempts*

---

##### Description

choose_overlapping selects top candidate nesting attempt among those that are temporally overlapping.

##### Usage

```
choose_overlapping(attempts)
```

##### Arguments

| | |
|---|---|
| attempts | data.frame of revisitation patterns of candidate nests |

##### Details

Within the function nest_finder, choose_overlapping is used when discard_overlapping = TRUE.

If the list of nest candidates includes temporally overlapping nesting attempts, only the candidate with the most visits is kept and the others get discarded. This is based on the rationale that an individual cannot simultaneously nest at more than one location. The location that is visited the most is assumed to be the most likely true nest.

##### Value

Returns data.frame of revisitation patterns filtered to only include non-temporally overlapping candidate nests

---

compare_buffers *Perform sensitivity analysis of buffer size for nest detection*

---

##### Description

compare_buffers compares nest identification results obtained using buffers of different size. This function is useful to perform sensitivity analyses of nest-detection results as the chosen buffer size varies. This helps both to demonstrate robustness of results independent of buffer size as well as to choose an appropriate buffer size for the species and data at hand. Users can specify a set of buffers that they wish to compare. Then, compare_buffers applies the same procedure as find_nests to the data using each of the different buffers. For the sake of keeping computation time at a manageable level, if the full dataset is large, it is recommended to only use a subset of data to test buffers on. If data on known nest locations are provided, compare_buffers also computes the following performance metrics for each buffer size:

##### Usage

```
compare_buffers(gps_data, buffers, known_coords, sp_tol = 100, min_pts,
  sea_start, sea_end, nest_cycle, min_d_fix, min_consec, min_top_att,
  min_days_att, discard_overlapping = TRUE)
```

##### Arguments

| | |
|---|---|
| gps_data | data.frame of movement data. Needs to include burst, date, long, lat |
| buffers | A vector of buffer sizes |
| known_coords | data.frame of coordinates for known nests. Needs to include burst, long, lat. |
| sp_tol | Integer. Spatial tolerance value: maximum distance tolerated between the estimated location of a nest and its actual (known) location. Defaults to 100 meters. |
| min_pts | Minimum number of points within a buffer |
| sea_start | Character string. Earliest date to be considered within the breeding season. Month and day, format "mm-dd" |
| sea_end | Character string. Latest date to be considered within the breeding season. Month and day, format "mm-dd" |
| nest_cycle | Duration of nesting cycle |
| min_d_fix | Minimum number of fixes for a day to be retained if no nest visit was recorded |
| min_consec | Minimum number of consecutive days visited |
| min_top_att | Minimum percent of fixes at a location on the day with maximum attendance |
| min_days_att | Minimum percent of days spent at a location between first and last visit |
| discard_overlapping | If results include temporally overlapping attempts, select only one among those? Defaults to TRUE. |

##### Details

- Positive predictive value, i.e., the proportion of identified nests that are known to be true nests;
- Sensitivity, i.e., the proportion of known nests that were successfully identified;
- False negative rate, i.e., the proportion of known nests that we failed to identify.

The false positive rate, i.e., the proportion of spurious nests among those identified, is not computed because it would require the assumption that any nests that are detected but are not known are spurious, which is not reasonable in most situations. Users can evaluate which buffer size is more appropriate based on the performance metrics. Regardless of whether data on known nest locations are provided, `compare_buffers` computes the total number of nests found for each buffer size as well as the number of nests found for each burst. If the user provides data on known nests, they can also specify a value of spatial tolerance to use as a cutoff to establish whether a nest was correctly identified: since coordinates of detected nests rarely match exactly the actual (known) coordinates of the nest, users can define which distance between real and estimated location they are willing to accept at most. If no data on true nests are available, these numbers can provide an indication of whether a given buffer size likely results in over- or underestimation of the number of nests. `compare_buffers` takes almost exactly the same arguments as `find_nests`, except for the argument 'buffer' which is here replaced with 'buffers' and for the additional arguments 'known_coords' and 'sp_tol'.

##### Value

Returns a `list` of variable length including:

- As many data frames of nest locations as the number of buffers specified (formatted like 'nests' in the output of `find_nests`);
- A `data.frame` reporting the total number of nests identified with each buffer size;
- A `data.frame` reporting the total number of nests identified with each buffer size for each burst;
- If 'known_coords' is provided, a `data.frame` reporting the positive predictive value obtained with each buffer size;
- If 'known_coords' is provided, a `data.frame` reporting the sensitivity obtained with each buffer size;
- If 'known_coords' is provided, a `data.frame` reporting the false negative rate obtained with each buffer size;

---

#### `date_handler` *Process dates and subset data within nesting season*

---

##### Description

`date_handler` finds the ymd starting and ending dates of the nesting season based on the dates given in input (as Julian day or a character string indicating month and day), and returns the subset of data that falls within the range. Inputting month and day is the recommended option.

#### Usage

```
date_handler(dat, sea_start, sea_end)
```

#### Arguments

| | |
|---|---|
| dat | `data.frame` of movement data for a single burst. Needs to include burst, date, long, lat |
| sea_start | Character string for month and day ("mm-dd") in which the nesting season starts |
| sea_end | Character string for month and day ("mm-dd") in which the nesting season ends |

#### Value

Returns subset of data comprised within the nesting season

---

### discriminate_nests *Find set/s of parameter values to discriminate nests*

---

#### Description

`discriminate_nests` uses CART to find sets of parameter values that best distinguish nests from non-nests among revisited locations.

#### Usage

```
discriminate_nests(explodata, train_frac)
```

#### Arguments

| | |
|---|---|
| explodata | `data.frame` of nests and non-nests as output by `get_explodata` |
| train_frac | Numeric. The fraction of data to use for training |

#### Details

Given a dataset of revisited locations flagged as either nests or non-nests, `discriminate_nests` uses Classification and Regression Trees (CART) to find the set (or sets) of revisitation parameters that best distinguishes between nests and non-nests.

The function fits a CART model on the training fraction of the data, prunes the tree, and performs cross-validation using the testing fraction of the data.

The user can specify how much of the data is used for training versus testing the algorithm. If all the data is used for training (`train_frac = 1`), cross-validation is not possible and error rates are not estimated.

The CART uses the following model formula:

`nest ~ consec_days + perc_days_vis + perc_top_vis`

The original tree is automatically pruned based on minimum error criterion: the tree is pruned back to the point where the cross-validated relative error (X-rel error) is at its minimum. If multiple trees compete at the minimum X-rel error, the smallest tree is picked.

#### Value

A `list` with the Type I and II estimated error rates (where applicable) and a plot of the CART output.

---

`dist_mat` *Calculate distance matrix between points in the data*

---

#### Description

`dist_mat` calculates pairwise distances between all points in the data.

#### Usage

```
dist_mat(dat)
```

#### Arguments

| | |
|---|---|
| `dat` | `data.frame` of movement data for a single burst. Needs to include burst, date, long, lat |

#### Details

Distances are calculated using the function `distGeo`. Takes advantage of `data.table` for a fast implementation. Adapted from a post on StackOverflow.

#### Value

Returns `data.table` with distance matrix.

---

`dummy` *Dummy function for BJS to test git*

---

#### Description

BJS wrote this function just to test git

#### Usage

```
dummy()
```

dummy2 *Second dummy function for BJS to test git*

#### Description

BJS wrote this function just to test git

#### Usage

```
dummy2()
```

estimate_outcomes *Estimate nesting outcomes*

#### Description

estimate_outcomes fits a Bayesian hierarchical model to the histories of nest revisitation to estimate the outcome of nesting attempts.

#### Usage

```
estimate_outcomes(fixes, visits, model = "null",
  mcmc_params = list(burn_in = 1000, n_chain = 2, thin = 5, n_adapt =
  1000, n_iter = 10000))
```

#### Arguments

| | |
|---|---|
| fixes | A matrix of the number of GPS fixes on each day of the nesting attempt as returned by format_visits. |
| visits | A matrix of the number of nest visits on each day of the nesting attempt as returned by format_visits. |
| model | Type of model to be run. One of "null", "p_time", "phi_time", or "phi_time_p_time". |
| mcmc_params | List of MCMC parameters. burn_in, n_chain, thin, n_adapt, n_iter |

#### Details

Data can be passed from format_visits to the function arguments fixes and visits.

The function runs a JAGS MCMC with uninformative priors for the estimation of nest survival. Parameters for the MCMC can be specified by the user by passing them as a list to mcmc_params.

The user can choose among four possible models: a null model with constant p and phi; a model where p varies with time; a model where phi varies with time; and a model where both p and phi vary with time.

#### Value

A list of mcarray objects.

#### `explore_nests` *Launch Shiny app for nesting data exploration*

##### Description

`explore_nests` launches a Shiny app for dynamic exploration and visualization of nests from movement data

##### Usage

```
explore_nests(gps_data)
```

##### Arguments

| | |
|---|---|
| `gps_data` | `data.frame` of movement data. Needs to include burst, date, long, lat |

##### Details

The function takes as input `gps_data` (see `find_nests` or vignette for details on data format) and launches a Shiny app that allows the user to dynamically explore repeatedly visited locations for one burst at a time while interactively tuning input parameters (see `find_nests` for the complete list). Under the hood, `explore_nests` runs `find_nests`, and then displays the results on a map.

##### Value

nothing

#### `find_nests` *Find nest locations from GPS data*

##### Description

`find_nests` finds nest locations from GPS data based on patterns of location revisitation

##### Usage

```
find_nests(gps_data, buffer, min_pts, sea_start, sea_end, nest_cycle,
  min_d_fix, min_consec, min_top_att, min_days_att,
  discard_overlapping = TRUE)
```

##### Arguments

| | |
|---|---|
| gps_data | data.frame of movement data. Needs to include burst, date, long, lat |
| buffer | Size of the buffer to compute location revisitation |
| min_pts | Minimum number of points within a buffer |
| sea_start | Character string. Earliest date to be considered within the breeding season. Month and day, format "mm-dd" |
| sea_end | Character string. Latest date to be considered within the breeding season. Month and day, format "mm-dd" |
| nest_cycle | Duration of nesting cycle |
| min_d_fix | Minimum number of fixes for a day to be retained if no nest visit was recorded |
| min_consec | Minimum number of consecutive days visited |
| min_top_att | Minimum percent of fixes at a location on the day with maximum attendance |
| min_days_att | Minimum percent of days spent at a location between first and last visit |
| discard_overlapping | If results include temporally overlapping attempts, select only one among those? Defaults to TRUE. |

##### Details

Data passed to the argument gps_data needs to be split in individual-years labelled each as a separate burst. We recommend dividing the data so that seasonal nesting activities are contained within single bursts. Cutting data at a day that is not likely to overlap with nesting is best.

Data must include the following columns: a burst identifier (burst), date-time (date), and long/lat coordinates (long, lat).

Patterns of revisitation to repeatedly visited locations are used to identify potential nesting locations. Due to both movement and GPS error, recorded points around recurrently visited locations are expected to be scattered around the true revisited location. To account for this scattering, the user defines a buffer value which will be used to group points falling within a buffer distance from each other.

When grouping, several points peripheral to a true revisited location may compete in grouping points around them. We term these 'competing points'. If the buffers of two points do not overlap, those points are not competing. Among competing points, only one point is selected, chosen as the one that incorporates the most other points within its buffer. A top candidate is selected for each cluster of competing points, i.e., one representative for each cluster around a revisited location.

To speed up calculations, the user can define min_pts as the minimum number of points that need to fall within the buffer for a point to be considered as a potential nest candidate. This discards isolated points from consideration as revisited locations.

The arguments sea_start and sea_end are used to delimit the nesting season. The user can pass either a Julian day or a date. If inputting dates, the year can be a dummy year which will get automatically updated each time to the correct year for the current burst. If working with a species for which the temporal limits of the nesting season are not well-defined, the user can input a range of dates that covers the entire year. Nonetheless, we recommend ensuring that sea_start and sea_end are set so that nesting attempts are not split between bursts. For example, for a species that nests from October to September, enter October 1st as start date and September 30th as end date and not, for example, January 1st-December 31st.

The argument `nest_cycle` is the duration (in days) of a complete nesting attempt, i.e., the time necessary for an individual to successfully complete reproduction.

Once recurrently visited locations are identified, the function computes, for each of them:

- the first and last day when the location was visited;
- the total number of visits;
- the number of days in which it was visited;
- the percent of days visited between the days of first and last visit;
- the attendance (% of fixes at the location) on the day with the most visits;
- the longest series of consecutive days visited;
- the estimated start and end dates of the nesting attempt.

On days when no visit was recorded, two cases are possible: either the nest was truly not visited, or visits were missed. On days with few fixes, there is a higher chance of missing a visit given that it happened. Missed visit detections can interrupt an otherwise continuos strike of days visited. To counteract possible issues due to missed visit detections, the user can define `min_d_fix` as the minimum number of fixes that have to be available in a day with no visits for that day to be retained when counting consecutive days visited. If a day with no visits and fewer fixes than `min_d_fix` interrupts a sequence of consecutive days visited, it does not get considered and the sequence gets counted as uninterrupted.

The remaining arguments are used to filter results. The user can set minimum values for each of the following revisitation statistics:

- the longest series of consecutive days visited (`min_consec`);
- the attendance (% of fixes at the location) on the day with the most visits (`min_top_att`);
- the percent of days visited between the days of first and last visit `min_days_att`;

Among candidate nests, only those whose values for the above parameters exceed the user-defined minima are returned.

If the results include any temporally overlapping nesting attempts, the user can opt to only keep one among those. If `discard_overlapping` is set to `TRUE` (default), only the candidate nest with the most visits is kept among temporally overlapping ones, and the others get discarded. This is based on the rationale that an individual cannot simultaneously nest at more than one location. The location that is visited the most is assumed to be the most likely true nest. On the other hand, setting `discard_overlapping` to `FALSE` retains all candidate nests in the results.

After identifying all nests that correspond to the criteria defined in input, the function appends a new column to the original GPS data that flags fixes recorded at a nest with the location identifier of that nest. The result is a history of nest visits for each burst.

#### Value

Returns a `list` of two elements: first, ‘nests‘, a `data.frame` of nest locations and associated revisitation stats; second, ‘visits‘, a `data.frame` of nest revisitation histories.

---

`foo` *foo: A package for computating the notorious bar statistic.*

---

#### Description

The foo package provides three categories of important functions: foo, bar and baz.

#### Foo functions

The foo functions ...

---

`format_attempts` *Format data for nesting outcome estimation*

---

#### Description

`format_attempts` takes as input the output of `find_nests` and formats it for input in `estimate_outcomes`.

#### Usage

```
format_attempts(nest_info, nest_cycle)
```

#### Arguments

| | |
|---|---|
| `nest_info` | Output of `find_nests` |
| `nest_cycle` | Duration of nesting cycle |

#### Details

The history of nest revisitation in the ‘visits‘ data frame in output from `find_nests` gets formatted as a matrix indicating, for each day, the number of GPS points at the nest. This is the ‘visits‘ matrix that `format_attempts` will output. Concurrently, another matrix is created, ‘fixes‘, indicating the number of GPS points available on each day.

#### Value

A `list` with two matrices: ‘fixes‘, a matrix of GPS fixes available on each day of the attempt; and ‘visits‘, a matrix of nest visits on each day of the attempt.

get_candidates *Get top candidate nests from possible competitors*

#### Description

get_candidates uses a distance matrix returned by dist_mat and a user-defined buffer to select candidate nest sites.

#### Usage

```
get_candidates(dm, buffer, min_pts = 2)
```

#### Arguments

| | |
|---|---|
| dm | Distance matrix returned by dist_mat |
| buffer | Buffer distance (in meters) used to group points |
| min_pts | Minimum number of points within the buffer for a point to be retained. Defaults to 2 |

#### Details

Due to both movement and GPS error, recorded points around recurrently visited locations are expected to be scattered around the true revisited location. The buffer is meant to account for this scattering, by grouping points that fall within the buffer distance.

When grouping, several points peripheral to the true revisited location may compete in grouping points around them. We term these 'competing points'. If the buffers of two points do not overlap, those points are not competing. Based on the assumption that the true revisited location is the one that incorporates the most points within its buffer, get_candidates compares the number of points that fall within the buffers of competing points and selects the one that includes the most.

A top candidate is selected for each cluster of competing points, i.e., one representative for each cluster around a revisited location. If there are multiple revisited locations with non-competing points, independent top candidates are all returned.

To speed up calculations, the user can define min_pts as the minimum number of points that need to fall within the buffer for a point to be considered as a potential candidate. This discards isolated points from consideration as revisited locations.

#### Value

Returns data.frame relating original location identifiers (loc_id) to the identifier of the corresponding candidate nest (group_id).

#### get_candidates_multi *Get top candidate nests from possible competitors (multiple buffers)*

##### Description

get_candidates_multi is an adaptation of get_candidates for multiple buffers. It is called by compare_buffers.

##### Usage

```
get_candidates_multi(dmat, buffers, min_pts = 2)
```

##### Arguments

| | |
|---|---|
| dmat | Distance matrix returned by dist_mat |
| buffers | Vector of buffer size (in meters) used to group points |
| min_pts | Minimum number of points within the buffer for a point to be retained. Defaults to 2 |

##### Value

Returns a list of data frames relating original location identifiers (loc_id) to the identifier of the corresponding candidate nest (group_id) for each of the specified buffers.

#### get_explodata *Extract nests and non-nests from revisited locations*

##### Description

get_explodata uses known nest locations to extract an equal number of true nests and non-nests from a set of revisited locations for exploration of parameter values.

##### Usage

```
get_explodata(candidate_nests, known_coords, known_ids, buffer,
  pick_overlapping = TRUE)
```

##### Arguments

| | |
|---|---|
| candidate_nests | data.frame of candidate nests output by find_nests |
| known_coords | data.frame of coordinates for known nests. Needs to include burst, long, lat. |
| known_ids | data.frame of location IDs for known nests. Needs to include burst and loc_id. |
| buffer | Integer. Buffer distance (in meters) used to select true nest location among candidates when known_coords is provided. |

pick_overlapping
: Logical. If TRUE (default), the non-nest is picked among those whose time range overlaps with the true nesting attempt.

#### Details

If no prior information is available to the user about which parameter values to use to filter nests among revisited locations, some exploration of the data is necessary. The objective of the exploration phase is to identify the set of parameter values that best discriminates between nests and non-nests in the species or population at hand.

Our suggested procedure consists in running a first coarse screening of revisited locations, using loose thresholds of behavioral parameters for filtering. For example, using `min_consec = 1`, `min_top_att = 1`, and `min_days_att = 1` in `find_nests`. In most cases, previous knowledge about the biology of the species should allow the user to already provide values for `sea_start`, `sea_end`, and `nest_cycle`. The output of this first screening is a set of revisited locations at the `buffer` distance of choice, which should include some nests as well as other repeatedly visited locations that are not nests. Comparing the values of behavioral parameters at nests versus non-nests can inform the choice of parameter values for later analysis.

This procedure requires knowledge of true nest locations for at least a subset of the data. Given some data on known nest locations and the coarse-screening output of `find_nests`, the function `get_explodata` identifies true nests and an equal number of non-nests to compare them to.

Comparing behavioral parameter values at nests versus non-nests will allow the user to find the set of parameter values that best discriminates between them. The resulting set of parameters can then be used to find nests in new data or in a subset of data for which no prior information on nests is available.

The user can pass data on known nests as either coordinates or location IDs. Ideally, prior and independent information on nest locations is available for a subset of the data. In this case, we recommend that coordinates are passed to the function argument `known_coords`. When coordinates of true nests are not known a-priori but the user is able to visually inspect revisited locations and identify those that are true nests (for example because they fall within known colonies), providing location IDs for true nests in `known_ids` is an alternative option.

When passing `known_coords`, the user is required to also specify a value for `buffer`. Because of GPS error, the coordinates of the point representing the true nest in `candidate_nests` might not exactly match those of the known nest location. If coordinates of true nests are provided rather than location IDs, the function selects the true nest among the set of candidates by choosing the candidate with the most visits among those that fall within a `buffer` distance from the known nest location. We recommend using for this argument the same value used for argument `buffer` in `find_nests`.

#### Value

A `data.frame` including an equal number of true nests and non-nests and their revisitation parameters.

#### hello — *Hello, World!*

##### Description

Prints 'Hello, world!'.

##### Usage

```
hello()
```

##### Examples

```
hello()
```

#### initialize_z — *Set initial known-state values prior to JAGS MCMC*

##### Description

This function is used in `estimate_outcomes` Used to initialize the latent variable `z` in the JAGS MCMC. Sets `z` = 1 for all time steps in between the first sighting and the last sighting.

##### Usage

```
initialize_z(ch)
```

##### Arguments

| | |
|---|---|
| `ch` | Capture history, i.e., matrix of nest visits |

##### Value

A matrix of initial states for each nesting attempt

#### julian_to_date *Conversion of Julian day to date*

##### Description

julian_to_date returns a date corresponding to Julian day and year in input.

##### Usage

```
julian_to_date(jd, yr)
```

##### Arguments

| | |
|---|---|
| jd | Integer. Julian day to convert. |
| yr | Integer. Year the Julian date refers to. |

##### Value

Returns a date in ymd format.

##### Examples

```
julian_to_date(1, 2000)
# Returns January 1st of 2000

# Test what happens with leap years

julian_to_date(100, 1991)
# 1991 was not a leap year; Julian day 100 is April 10th

julian_to_date(100, 1992)
# 1992 was a leap year; Julian day 100 is April 9th
```

#### perf_metrics *Compute nest-detection performance metrics*

##### Description

perf_metrics calculates nest detection performance metrics based on the results of find_nests and data on known nest locations. The user defines the spatial tolerance threshold between the estimated location of a nest and its real (known) location. The performance metrics are defined as follows:

##### Usage

```
perf_metrics(gps_data, nest_info, known_coords, min_consec, sp_tol = 100)
```

##### Arguments

| | |
|---|---|
| gps_data | Original `data.frame` of movement data. Needs to include burst, date, long, lat |
| nest_info | Output of `find_nests` |
| known_coords | `data.frame` of coordinates for known nests. Needs to include burst, long, lat |
| min_consec | The minimum number of consecutive days visited that was used when running `find_nests` |
| sp_tol | Integer. Spatial tolerance value: maximum distance tolerated between the estimated location of a nest and its actual (known) location. Defaults to 100 meters. |

##### Details

- Positive predictive value, i.e., the proportion of identified nests that are known to be true nests;
- Sensitivity, i.e., the proportion of known nests that were successfully identified;
- False negative rate, i.e., the proportion of known nests that we failed to identify.

The false positive rate, i.e., the proportion of spurious nests among those identified, is not computed because it would require the assumption that any nests that are detected but are not known are spurious, which is not reasonable in most situations.

---

#### plot_detection *Plot detection probability over time from MCMC run*

---

##### Description

`plot_detection` makes a plot of mean nest detection probability over time as estimated by `estimate_outcomes`.

##### Usage

```
plot_detection(mcmc_obj, ci = 0.95)
```

##### Arguments

| | |
|---|---|
| mcmc_obj | List of `mcarrays` output by `estimate_outcomes`. |
| ci | Numeric. Credible interval level. |

##### Value

A plot of nest detection probability over time.

#### plot_nest_surv *Plot nest survival over time from MCMC run*

##### Description

`plot_nest_surv` makes a plot of nest survival over time for a chosen nesting attempt as estimated by `estimate_outcomes`.

##### Usage

```
plot_nest_surv(mcmc_obj, who = 1, ci = 0.95)
```

##### Arguments

| | |
|---|---|
| mcmc_obj | List of `mcarrays` output by `estimate_outcomes`. |
| who | Integer. Which nesting attempt to plot. |
| ci | Numeric. Credible interval level. |

##### Value

A plot of individual nest survival over time.

#### plot_survival *Plot survival over time from MCMC run*

##### Description

`plot_survival` makes a plot of mean nest survival over time as estimated by `estimate_outcomes`.

##### Usage

```
plot_survival(mcmc_obj, ci = 0.95)
```

##### Arguments

| | |
|---|---|
| mcmc_obj | List of `mcarrays` output by `estimate_outcomes`. |
| ci | Numeric. Credible interval level. |

##### Value

A plot of nest survival over time.

---

`revisit_stats` *Calculate revisitation patterns*

---

#### Description

`revisit_stats` calculates patterns of revisitation at candidate nests.

#### Usage

```
revisit_stats(dat, sub, sea_start, sea_end, min_d_fix, min_consec,
  nest_cycle)
```

#### Arguments

| | |
|---|---|
| `sub` | `data.frame`. Subset of movement data corresponding to candidate nests |
| `sea_start` | Integer (if Julian day) or date in which the nesting season starts |
| `sea_end` | Integer (if Julian day) or date in which the nesting season ends |
| `min_d_fix` | Integer. Minimum number of fixes in a day for that day to be counted as not visited if a visit was not observed |

#### Details

This is a wrapper function that calls `visit_rle`, `rle_to_consec`, and `attempt_limits`.

For each candidate nest, the function computes the first and last day when the location was visited, the total number of visits, the number of days in which it was visited, the percent of days with a visit, the attendance on the day with the most visits (percent locations at the nest over the total number of fixes on that day), the longest series of consecutive days visited, and the estimated start and end dates of the nesting attempt.

On days when no visit was recorded, two cases are possible: either the nest was truly not visited, or visits were missed. On days with few fixes, there is a higher chance of missing a visit given that it happened. Missed visit detections can interrupt an otherwise continuos strike of days visited. To counteract possible issues due to missed visit detections, the user can define `min_d_fix` to set a minimum number of fixes that have to be available in a day with no visits for that day to be retained when counting consecutive days visited. If a day with no visits and fewer fixes than `min_d_fix` interrupts a sequence of consecutive days visited, it does not get considered and the sequence gets counted as uninterrupted.

#### Value

Returns `data.frame` with revisitation statistics for each candidate nest.

#### rle_to_consec *Calculate consecutive days visited*

##### Description

rle_to_consec calculates the longest sequence of consecutive days a candidate nest was visited.

##### Usage

```
rle_to_consec(rl_df)
```

##### Arguments

| | |
|---|---|
| rl_df | data.frame of run-length encoding of nest visits |

##### Details

Used with lapply inside function revisit_stats. Computes longest series of consecutive days a candidate nest was visited. Takes as input the output of visit_rle.

##### Value

Returns maximum number of consecutive days when the candidate nest was visited.

#### summarize_outcomes *Summary of outcomes from MCMC run*

##### Description

summarize_outcomes returns summary statistics of estimated nesting attempt outcomes.

##### Usage

```
summarize_outcomes(mcmc_obj, ci = 0.95)
```

##### Arguments

| | |
|---|---|
| mcmc_obj | List of mcarrays output by estimate_outcomes. |
| ci | Numeric. Credible interval level. |

#### Details

The function takes as input a list of `mcarrays` output by `estimate_outcomes` and returns a list including the following:

- mean, lower and upper credible interval values (at the level specified by the user) for both survival and detection probability, where applicable (depending on which model formula was chosen);
- for each attempt, mean, lower and upper credible interval values of the probability of success and of the failure date. If the estimated probability of success is 1, the failure date corresponds to the duration of a complete nesting attempt.

#### Value

A list with (a) the population-level survival and (b) detection parameters and (c) the individual nest fates

---

`visit_rle` *Run-length encoding of nest visits*

---

#### Description

`visit_rle` calculates the run-length encoding of visits based on the daily history of revisitation of a candidate nest.

#### Usage

```
visit_rle(x)
```

#### Arguments

| | |
|---|---|
| `x` | `data.frame` of daily revisitation history of a candidate nest |

#### Details

Used with `lapply` inside function `revisit_stats`. Performs run-length encoding of nest visits and formats as `data.frame` for later use.

#### Value

Returns `data.frame` of run-length encoding of nest visits.

### Index

attempt_limits, 2

candidate_summary, 3
check_input, 4
choose_overlapping, 4
compare_buffers, 5, *15*

date_handler, 6
discriminate_nests, 7
dist_mat, 8, *14, 15*
distGeo, *8*
dummy, 8
dummy2, 9

estimate_outcomes, 9
explore_nests, 10

find_nests, *5, 6*, *10*, 10, *18*
foo, 13
foo-package *(*foo*)*, 13
format_attempts, 13
format_visits, *9*

get_candidates, *3*, 14, *15*
get_candidates_multi, 15
get_explodata, 15

hello, 17

initialize_z, 17

julian_to_date, 18

perf_metrics, 18
plot_detection, 19
plot_nest_surv, 20
plot_survival, 20

revisit_stats, 21
rle_to_consec, 22

summarize_outcomes, 22

visit_rle, 23
