## Additional file 1 for "Analysis of movement recursions to detect reproductive events and estimate their fate in central place foragers"

| Function name | Function purpose |
|---|---|
| compare_buffers() | Performs a sensitivity analysis on the effect of buffer size on nest-detection performance. |
| discriminate_nests() | Uses CART to identify one or more sets of parameter values that best discriminate between known nests and non-nests. The parameter values identified can then be manually entered in find_nests() to filter nests among revisited locations. |
| estimate_outcomes() | Runs Bayesian nest survival analysis to estimate the fate of breeding attempts. |
| explore_nests() | Launches Shiny app to interactively explore revisited locations on a map and visualize results based on different values of input parameters. |
| find_nests() | Finds revisited locations that satisfy revisitation parameter values provided in input. |
| format_attempts() | Formats GPS data into a presence/absence time series at the nest that is used as input for nest survival analysis. |
| get_explodata() | Selects one revisited non-nest location for each known nest provided. The resulting dataset of known nests and non-nests can be used in input in discriminate_nests(). |
| perf_metrics() | Computes performance metrics of the nest-detection algorithm (positive predictive value, sensitivity, and false negative rate). |
| plot_detection() | Plots detection probability through time at the population level based on results of nest survival analysis. |
| plot_nest_surv() | Plots nest survival through time for each individual attempt based on results of nest survival analysis. |
| plot_survival() | Plots nest survival through time at the population level based on results of nest survival analysis. |
| summarize_outcomes() | Returns summary statistics of reproductive fate based on results of nest survival analysis. |

Table 1 – Functions of R package nestR (https://github.com/picardis/nestR) and what they are used for. Only exported functions are shown. The complete package documentation (help files for both exported and non-exported functions) is in Additional file 2.
